# Retroviral Infection of Human Neurospheres and Use of Stem Cell EVs to Repair Cellular Damage

**DOI:** 10.1101/2020.12.31.424849

**Authors:** Heather Branscome, Pooja Khatkar, Sarah Al Sharif, Dezhong Yin, Sheela Jacob, Maria Cowen, Yuriy Kim, James Erickson, Christine A. Brantner, Nazira El-Hage, Lance A. Liotta, Fatah Kashanchi

**Affiliations:** Laboratory of Molecular Virology, School of Systems Biology, George Mason University, Manassas, VA; American Type Culture Collection (ATCC), Manassas, VA; ATCC Cell Systems, Gaithersburg, MD; GW Nanofabrication and Imaging Center, George Washington University, Washington, DC; Department of Immunology and Nano-Medicine, Herbert Wertheim College of Medicine, Florida International University, Miami, FL; Center for Applied Proteomics and Molecular Medicine, George Mason University, Manassas, VA

**Keywords:** Neurospheres, neural progenitor cells (NPCs), HIV-1, stem cells, extracellular vesicles (EVs)

## Abstract

HIV-1 remains an incurable infection that is associated with substantial economic and epidemiologic impacts. HIV-associated neurocognitive disorders (HAND) are commonly linked with HIV-1 infection; despite the development of combination antiretroviral therapy (cART), HAND is still reported to affect at least 50% of HIV-1 infected individuals. It is believed that the over-amplification of inflammatory pathways, along with release of toxic viral proteins from infected cells, are primarily responsible for the neurological damage that is observed in HAND; however, the underlying mechanisms are not well-defined. Therefore, there is an unmet need to develop more physiologically relevant and reliable platforms for studying these pathologies. In recent years, neurospheres derived from induced pluripotent stem cells (iPSCs) have been utilized to model the effects of different neurotropic viruses. Here, we report the generation of neurospheres from iPSC-derived neural progenitor cells (NPCs) and we show that these cultures are permissive to retroviral (e.g. HIV-1, HTLV-1) replication. In addition, we also examine the potential effects of stem cell derived extracellular vesicles (EVs) on HIV-1 damaged cells as there is abundant literature supporting the reparative and regenerative properties of stem cell EVs in the context of various CNS pathologies. Consistent with the literature, our data suggests that stem cell EVs may modulate neuroprotective and anti-inflammatory properties in damaged cells. Collectively, this study demonstrates the feasibility of NPC-derived neurospheres for modeling HIV-1 infection and, subsequently, highlights the potential of stem cell EVs for rescuing cellular damage induced by HIV-1 infection.

## Introduction

HIV-1 remains an incurable infection and as of 2018 approximately 37.9 million people were estimated to be living with HIV-1, with an average of 1.7 million new infections occurring annually [1]. In addition to its deleterious impact on the immune system, HIV-1 also contributes to neurocognitive impairments. HIV-associated neurocognitive disorders (HAND) are commonly associated with HIV-1 infection and comprise a range of neurological dysfunctions, including asymptomatic neurocognitive impairment, mild neurocognitive disorder, and HIV-associated dementia [2]. Despite the development of combination antiretroviral therapy (cART), which effectively suppresses viral replication and has vastly improved the quality of life for those treated, HAND is still reported to affect up to 50% of infected individuals [3, 4]. Therefore, HAND represents a considerable cause of morbidity in these populations.

Although the exact mechanisms that contribute to its persistence are poorly understood, it is believed that the over-amplification of inflammatory pathways, along with the release of toxic viral proteins from infected cells, are primarily responsible for the neurological damage that is observed in HAND [5,6,7]. While neurons are not directly infected by HIV-1, they remain highly susceptible to the cellular damage and injury mediated by viral proteins such as gp120 and Tat [8]. Additionally, the release of pro-inflammatory cytokines, including TNF-α, IL-1β and IL-6, from infected macrophages and microglia can also promote neuronal damage and neuronal apoptosis and it has also been suggested that exposure of astrocytes to these factors can further exacerbate neurotoxic effects [5, 6]. Consequently, there is a gap of knowledge surrounding the neuropathology of HAND and the mechanisms that contribute to its persistence.

The complexity of the Central Nervous System (CNS) relative to other systems in the body has largely limited the progress of CNS disease modeling and drug development. Moreover, the global incidence of CNS diseases, including neurocognitive and neurodegenerative disorders as well as neurological infections, has substantially risen throughout the last several decades. This, in turn, imposes a substantial socioeconomic burden. For example, a recent report found that approximately 16.5% of global deaths are caused by neurological diseases and the estimated costs associated with these diseases are expected to range from $600 to $700 million [9, 10]. This underscores the outstanding need for the development of novel research platforms which can allow for improved modeling of CNS pathologies, including those related to HIV-1 infection.

Improvements in biotechnology, coupled with progress in stem cell research, have revolutionized the biomedical field during the last decade. Specifically, induced pluripotent stem cells (iPSCs) represent a relatively novel platform that has allowed for advanced disease modeling and drug discovery [11]. Using this technology, adult somatic cells can be reprogrammed into an embryonic-like state via the overexpression of transcription factors (e.g. Oct3/4, SOX2, c-Myc, Klf4) responsible for maintaining pluripotency. Takahashi and Yamanaka first demonstrated this in 2006 with the successful reprogramming of mouse somatic cells into iPSCs and again, in 2007, when they generated iPSCs from human fibroblasts [12, 13]. In the context of the CNS, iPSCs have the potential to differentiate into a wide variety of neuronal and glial cell types including dopaminergic, cortical, and motor neurons as well as astrocytes, oligodendrocytes, microglia, and neural precursors [14, 15, 16, 17]. Along these lines, iPSCs have been employed to model a number of different CNS pathologies ranging from Alzheimer’s, Parkinson’s, and Huntington’s disease, to amyotrophic lateral sclerosis, multiple system atrophy, and Zika, West Nile virus, and HIV-1 [18, 19, 20, 21, 22, 23, 24, 25].

In recent years, human iPSCs have been used to derive neural stem cells (NSCs) and/or neural progenitor cells (NPCs); these neuronal cultures have subsequently been used to efficiently generate three-dimensional (3D) neurospheres *in vitro.* In general, 3D models are expected to be superior to traditional 2D culture models of NPCs due to their resemblance to the *in vivo* cellular environment which permits a more accurate representation of tissue-specific architecture and cell to cell communication [26, 27, 28]. Along these lines, it has been demonstrated that NPC-derived neurospheres are similar to those derived from primary fetal human NPCs with respect to migration, proliferation, and differentiation [29]. Additionally, it has previously been shown that neurospheres partially maintain regionally specific expression of developmental control genes *in vitro* and, moreover, that cells derived from these spheres retain the ability to differentiate into cellular subtypes representative of their regional origin [30, 31, 32]. Therefore, the use of 3D neurospheres is expected to serve as a complementary approach to *in vivo* studies relating to neurogenesis and neural development.

More recently, others have employed the use of 3D neurospheres to model the effects of different neurotropic viruses. For example, Garcez et al. investigated the consequences of Zika virus (ZIKV) infection and found that human iPSC-derived neurospheres exposed to ZIKV exhibited morphological abnormalities and had a higher induction of apoptosis relative to mock infection [33]. In a follow-up study which included proteomic and mRNA transcriptional profiling of infected neurospheres, these authors generated further data suggesting that ZIKV impaired pathways related to cell cycle and neuronal differentiation [34]. Additionally, D’Aiuto et al. utilized human iPSCs to generate 3D neurospheres to evaluate herpes simplex virus type 1 (HSV-1) infection [35]. The results from these studies demonstrated that neurospheres remained viable after four weeks in culture, were capable of differentiating into not only mature neurons (glutamatergic, GABAergic, dopaminergic) but also glial cells, and could be applied in a high throughput fashion for drug screening. It is also worth noting that while this manuscript was in-process, Dos Reis et al. published a similar study that utilized NPC-derived 3D organoids co-cultured with HIV-1 infected microglia. The results from these experiments not only confirmed the replication potential of HIV-1 in these cultures but were also consistent with the expected neurodegenerative and pro-inflammatory phenotypes associated with HIV-1 infection [36]. Taken together, this research strongly suggests that neurospheres are physiologically relevant models that can be used to study not only the cellular and molecular interactions in both healthy and diseased states but can also serve as a reliable platform for drug testing. Furthermore, they set an essential precedent for the modeling of other neurotropic viruses.

As described earlier, the development of CNS therapeutics has met relatively limited success. Factors relating to overall disease complexity, lengthy developmental periods and stringent regulatory review, coupled with the challenge of drug delivery across the blood-brain-barrier (BBB) have hindered progress in this field [37, 38]. While stem cell therapy has been studied in the context of CNS pathologies, there is now an emerging interest in the potential therapeutic application of stem cell-derived extracellular vesicles (EVs) as these vesicles have been shown to mediate many of the reparative and regenerative properties associated with their donor stem cells [39]. Briefly, EVs are nano-sized vesicles that are generally classified by their size and biogenesis. Two of the most widely studied subtypes of EVs are exosomes, which range approximately 50 to 150 nm in size and originate from the endosomal pathway, and microvesicles (MVs), which range approximately 50 to 1000 nm in size and are released directly from the plasma membrane [40]. EVs are enriched with a diverse array of biological cargo which includes RNAs, proteins, cytokines, and lipids and this cargo is believed to drive their potent functional effects [41, 42]. Accordingly, EVs from both mesenchymal stem cells (MSCs) and iPSCs have been broadly studied for their functional effects and there is rich literature to support their potential therapeutic application in various pathologies as reviewed elsewhere [43, 44].

With respect to the CNS, there is accumulating evidence which strongly indicates that stem cell EVs exert both immunomodulatory and neuroprotective effects to promote functional recovery in damaged tissue [45, 46, 47, 48]. However, it is important to note that this field is still in its infancy. Moreover, whether stem cell EVs may confer beneficial effects in the context of CNS infection, including HIV-1 and HAND, remains to be addressed. Thus, studies of this nature are imperative to better characterize the potential neuroprotective properties of stem cell EVs and to determine the mechanisms by which these EVs may repair cellular damage that is induced by different viral infections.

Here, we report the generation of neurospheres from NPCs derived from normal human iPSCs. These cultures not only exhibit reproducibility with respect to morphology and size but also demonstrate the ability to be stably maintained in culture to permit differentiation into different CNS cell types. Importantly, our data suggests that these cultures are permissive to HIV-1 infection. Additionally, we include data to show that neurospheres co-cultured with HTLV-1 infected cells may also be susceptible to HTLV-1 infection. Lastly, we evaluated the potential reparative effects of both MSC and iPSC EVs on HIV-1 damaged cells in both 2D cultures and 3D neurospheres. Collectively, this data highlights NPC-derived neurospheres as relevant and robust models for future studies relating to HIV-1 infection and HAND and, subsequently, demonstrates the potential application of stem cell EVs for rescuing cellular damage induced by HIV-1 infection.

## Results

### Generation and Characterization of NPC-derived Neurospheres

The ability of NPCs to aggregate and assemble into physiologically relevant 3D structures that are also permissive to viral replication, including Zika, HSV-1, and more recently HIV-1, has been published by others [27, 33, 34, 35, 36, 49]. To further expand upon these studies, we sought to evaluate the susceptibility of NPC-derived neurospheres to retroviral infection. For these studies we utilized commercially available NPCs that had been established from normal human iPSCs. The diagram in Fig. 1a summarizes the overall workflow and timeline associated with the generation of NPCs and neurospheres. Briefly, adult somatic cells were directly reprogrammed into iPSCs using a non-integrating Sendai-based reprogramming kit following established protocols [50]. To authenticate the quality of iPSCs, their germ line differentiation potential was confirmed by pluripotency testing (PluriTest); furthermore, flow cytometry was performed to confirm the positive expression (>85%) of the pluripotency stem cell markers SSEA4 and TRA-1-60. iPSCs then underwent embryoid body (EB) formation to assemble into spherical colonies that were subsequently differentiated into NPCs. NPCs were briefly scaled up in 2D culture prior to the induction of neurosphere formation.

**Figure 1.**
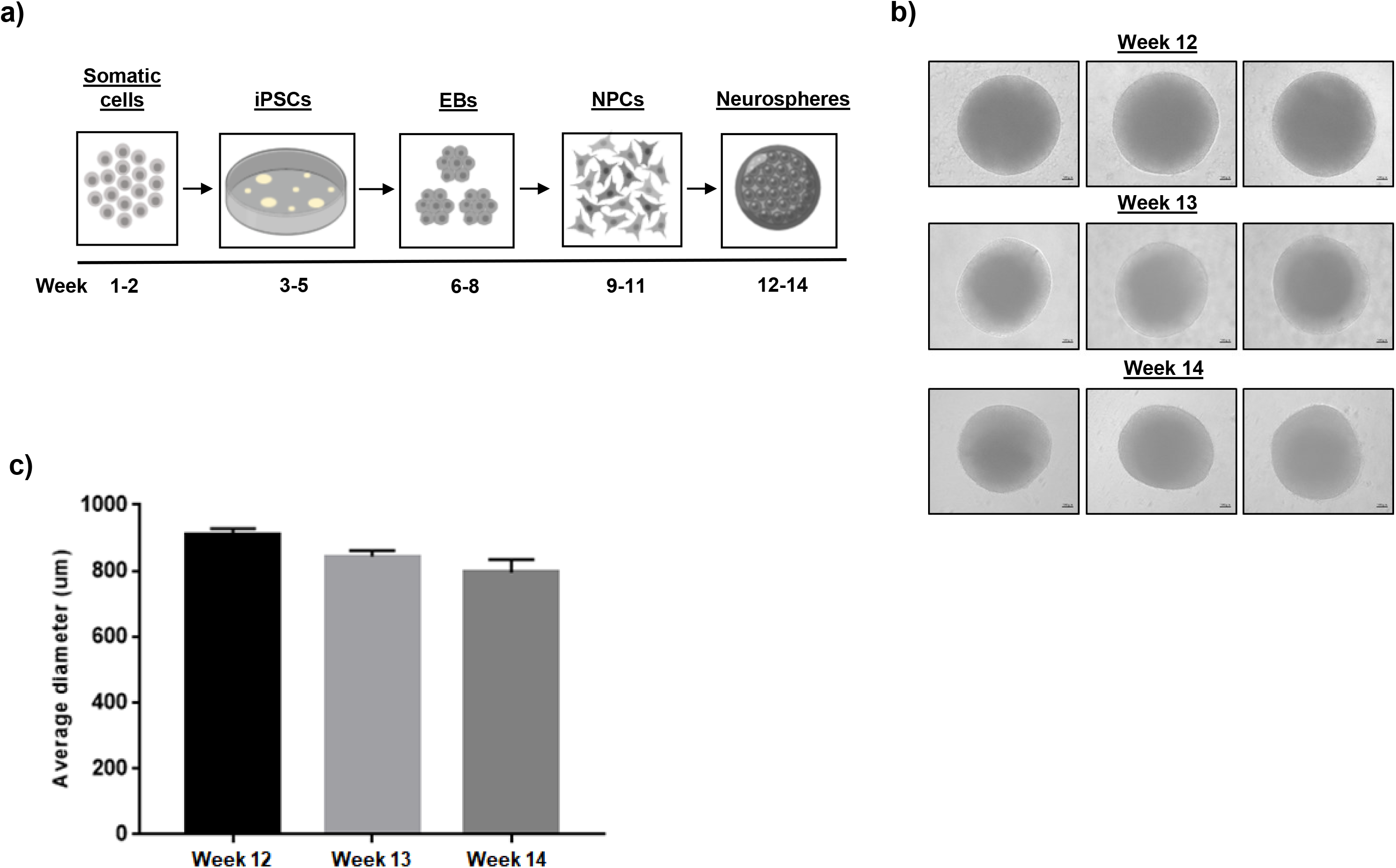
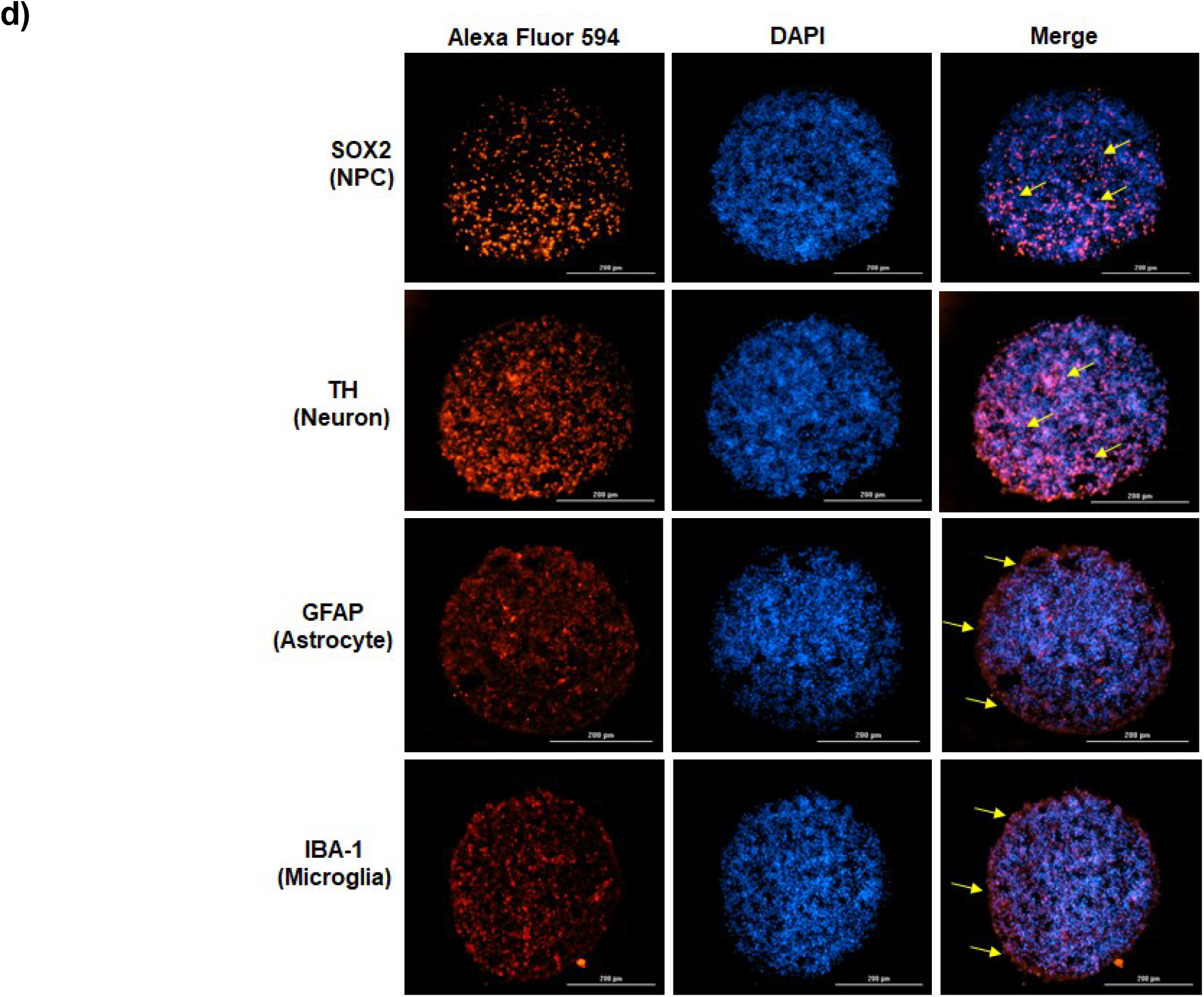
Generation and characterization of NPC-derived neurospheres. **(a)** General workflow and approximate timeline associated with the development of neurospheres using human somatic cells as the starting material. Figure created with BioRender.com. **(b)** Representative phase contrast images show the appearance and morphology of NPC-derived neurospheres during the course of the two week differentiation period (weeks 12 through 14). n=3. Scale bar = 100 µm. **(c)** Average diameter (µm) of NPC-derived neurospheres (two measurements per sphere) during the course of the differentiation period. n=3 neurospheres. **(d)** After the two week differentiation period (weeks 12 through 14), ICC staining was carried out on differentiated, cross-sectioned neurospheres. Nuclei were counter-stained with DAPI. Representative images show the relative expression of NPCs (SOX2), neurons (TH), astrocytes (GFAP), and microglia-like cells (IBA-1). Scale bar= 200 µm. n = 2 for ICC imaging.

In an effort to promote the controlled and consistent formation of neurospheres, NPCs were seeded at a high density in individual U-shaped wells of an ultra-low attachment 96-well plate. After 48 to 72 hours of incubation a single, well-defined neurosphere was visible in each well. Representative phase contrast images in Fig. 1b show that this methodology resulted in uniformly shaped spheres that maintained their morphology and structure throughout the two week differentiation process (weeks 12 through 14). Using calibrated imaging software, the average diameter of each sphere in Fig. 1b was measured and graphed. As shown in Fig. 1c, the average diameter of the neurospheres remained between 700 to 800 µm. This quantitative analysis indicates that NPC-derived neurospheres maintain a relatively consistent size over time and, moreover, demonstrates the reproducibility of our protocols.

As part of routine quality control testing, the NPCs used in these experiments were previously characterized via immunocytochemistry (ICC), which confirmed positive expression of the NSC markers Nestin and Pax6. Additionally, these NPCs have displayed robust expression of the early neuronal marker Tuj1, the dopaminergic marker tyrosine hydroxylase (TH), and the astrocytic marker GFAP when cultured in the appropriate differentiation medium (Supplementary Fig. 1a). This data is consistent with the reports of others who have shown that NPCs are capable of differentiating into not only mature neurons but also glial cells [51, 52, 53, 54, 55]. Given these results, we rationalized that NPC-derived neurospheres would retain their inherent differentiation capacity in 3D cultures.

Cross-sectioning and ICC staining of neurospheres that had been cultured in differentiation medium for two weeks indicated that neurospheres were composed of a mixed population of cells which included NPCs (SOX2^+^), dopaminergic neurons (TH^+^), astrocytes (GFAP^+^), and microglia-like (IBA-1^+^) cells. Representative images of stained cross-sections (Fig. 1d) suggest that both NPCs and neurons are relatively diffuse and uniformly distributed throughout the spheres. On the other hand, glial cells appeared to be more co-localized towards the periphery of the spheres, as indicated by expression of GFAP and IBA-1. Additionally, H&E histological staining illustrated the relatively compact and dense structure of cross-sectioned neurospheres (Supplementary Fig. 1b). Collectively, this data highlights the ability of iPSC-derived NPCs to self-assemble into neurospheres that maintain their ability to differentiate into both neurons and glial cells.

### Infection of Neurospheres

To assess the susceptibility of NPC-derived neurospheres to HIV-1 infection, neurospheres were cultured in the presence of HIV-1 dual tropic 89.6 (MOI:10). In addition, since HIV-1 infected individuals utilize combined antiretroviral therapy (cART), neurospheres were also treated with a cocktail consisting of equal parts lamivudine, tenofovir disoproxil fumarate, emtricitabine, and indinavir to model the effects of cART [1]. After a period of fourteen days in culture, supernatants were collected and neurospheres were harvested for downstream assays. To determine whether exposure of neurospheres to HIV-1 resulted in effective viral replication, total cellular protein was isolated and western blot was performed to assess the expression of different viral proteins. As shown in Fig. 2a, uncleaved Gag polyprotein Pr55, p24 capsid protein, and accessory protein Nef were all detected in neurospheres exposed to HIV-1 89.6.

**Figure 2.**
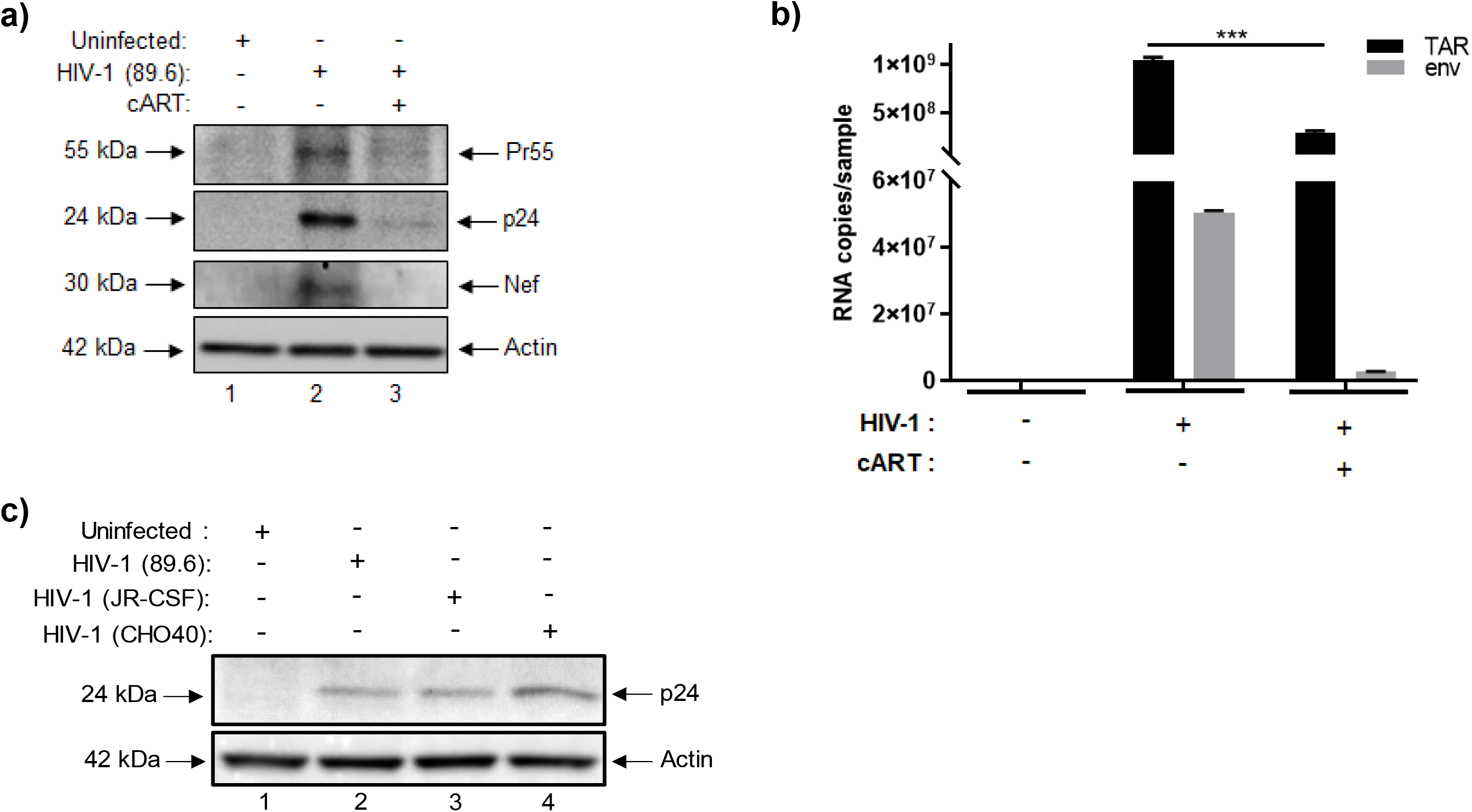
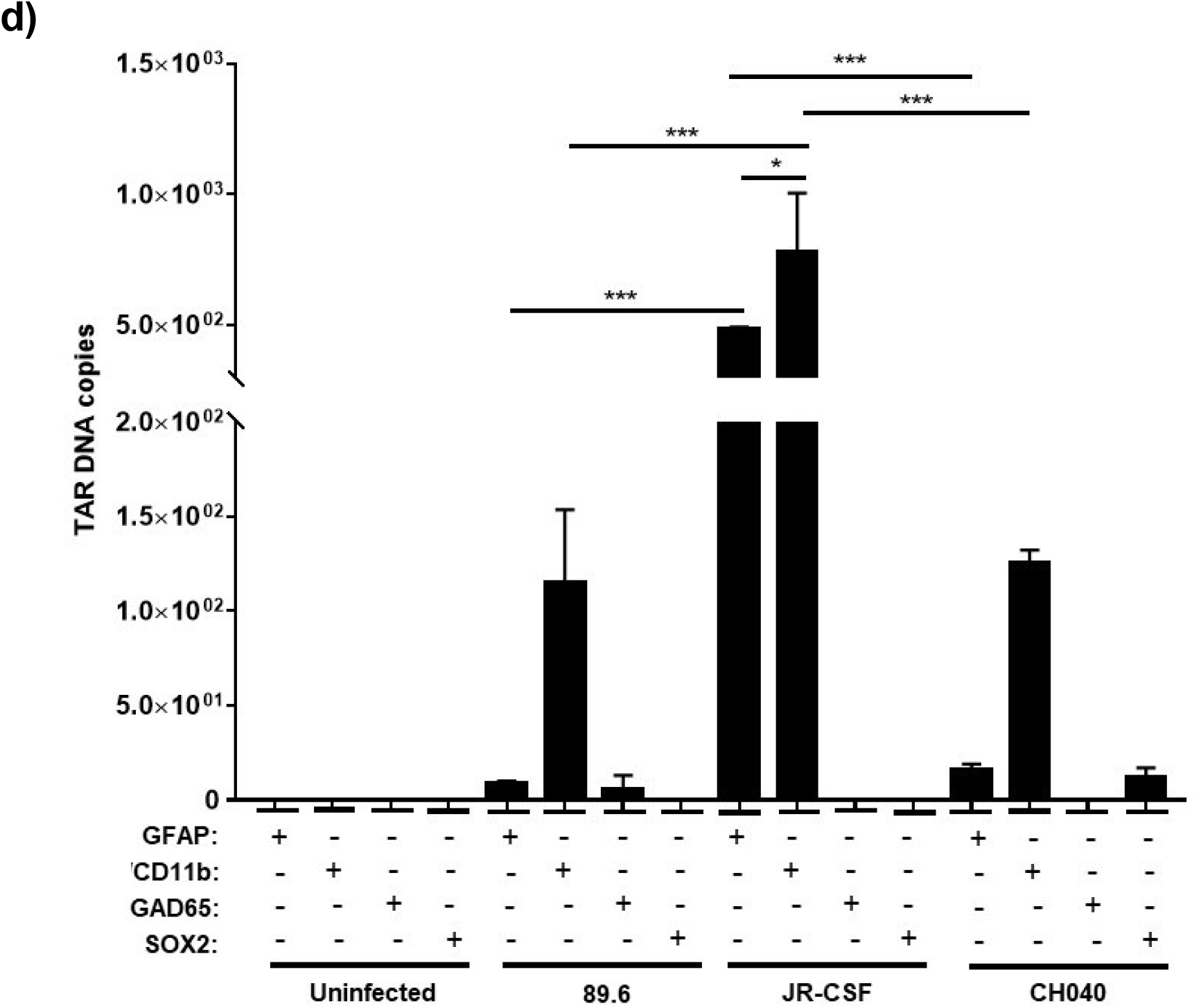
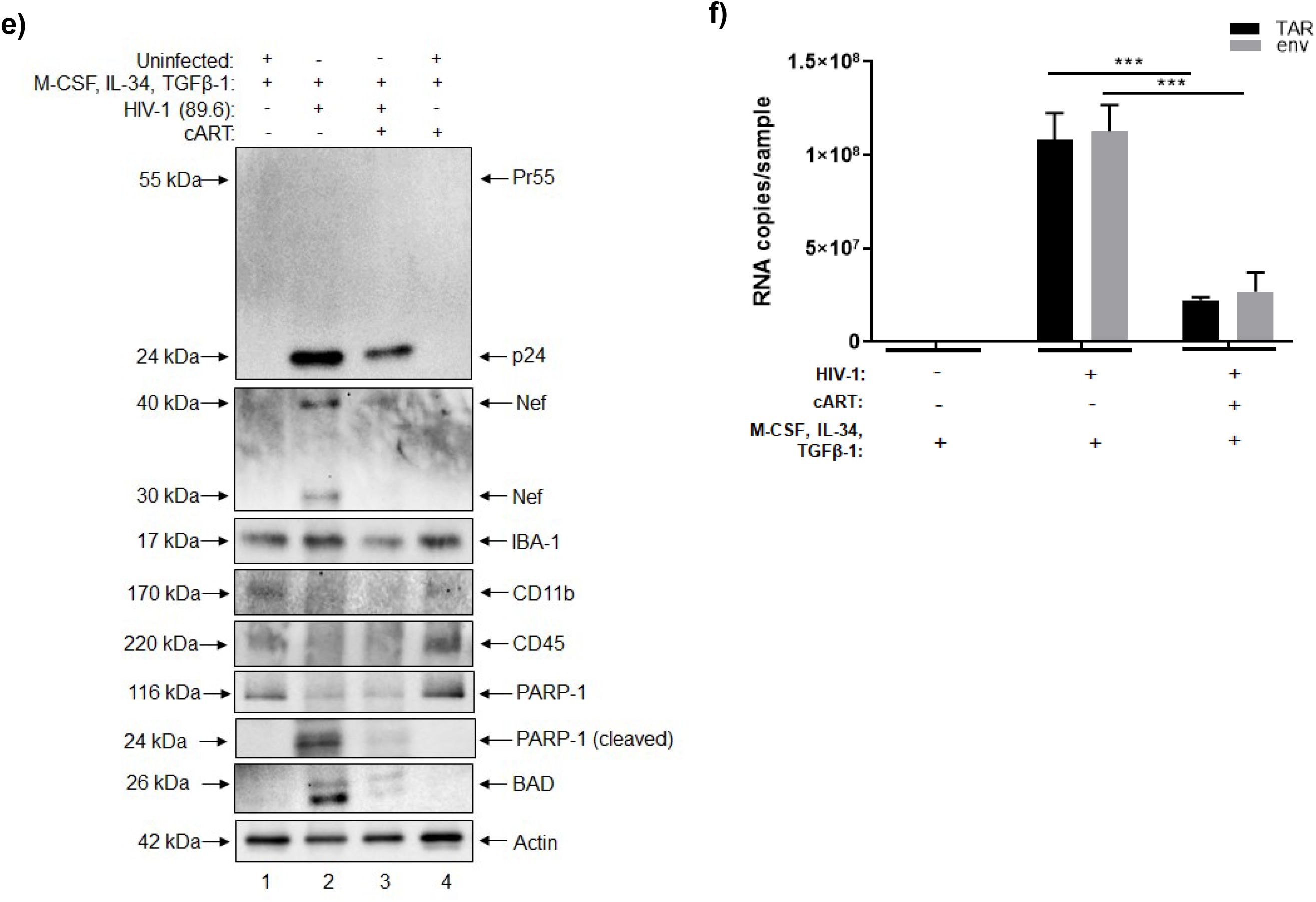
HIV-1 infection of NPC-derived neurospheres. Differentiated neurospheres were exposed to dual-tropic HIV-1 89.6, JR-CSF, or CHO40 (MOI:10). **(a)** Western blot was performed to examine the relative expression of Pr55, p24, and Nef in HIV-1 89.6 neurospheres cultured with or without cART (lamivudine, tenofovir disoproxil fumarate, emtricitabine, indinavir) after fourteen days. **(b)** RT-qPCR was performed to examine the relative copy numbers of TAR and *env* RNA in HIV-1 89.6 neurospheres with or without cART after fourteen days. n=3. ***p < 0.0001. **(c)** Western blot was performed to compare the relative expression of p24 in HIV-1 89.6, JR-CSF, and CHO40 infected neurospheres after seven days. **(d)** Neurospheres were gently disaggregated and immunoprecipitation was performed using antibodies specific for astrocytes (GFAP), microglia-like cells (CD11b), neurons (GAD65), and NSCs ( SOX2) followed by direct PCR to quantify copies of TAR DNA in different cellular subpopulations of HIV-1 89.6, JR-CSF, and CHO40-infected neurospheres. n=3. * p < 0.05, ***p < 0.0001. Differentiated neurospheres were treated with IL-34 (100 ng/mL), M-CSF (25 ng/mL), TGFβ-1 (50 ng/mL) every other day for a period of seven days and were then exposed to dual tropic HIV-1 89.6 for a period of fourteen days with or without cART (lamivudine, tenofovir disoproxil fumarate, emtricitabine, indinavir). **(e)** Western blot was performed to examine the relative expression of viral proteins (Pr55, p24, Nef), microglia markers (IBA-1, CD11b, CD45), and apoptotic proteins (PARP-1, BAD). **(f)** RT-qPCR was performed to quantify the relative copy numbers of TAR and *env* RNA. n=3. *** p < 0.0001.

The cART cocktail used for these experiments consisted of a mixture of nucleoside reverse transcriptase inhibitors (NRTIs) and a protease inhibitor; as expected, culturing of HIV-1 neurospheres with cART reduced the expression of Pr55, p24, and Nef to almost undetectable levels (Fig. 2a). RT-qPCR for HIV-1 TAR and *env* confirmed the presence of these RNAs in HIV-1 89.6-exposed neurospheres with approximately 10^9^ and 10^7^ copies being detected, respectively, relative to the control (uninfected) neurospheres (Fig. 2b). Treatment of HIV-1 neurospheres with cART decreased the expression of both TAR and *env* RNA relative to the HIV-1 sample, with a significant reduction in the expression of TAR.

To validate these results, we also exposed neurospheres to two other HIV-1 lab adapted strains, JR-CSF (MOI:10) and CHO40 (MOI:10) with or without cART. After seven days in culture, supernatants were collected and neurospheres were harvested for downstream assays. Total cellular proteins were isolated and, again, western blot showed the expression of p24 capsid protein in neurospheres that had been exposed to each viral isolate (Fig. 2c). Moreover, the level of p24 expression appeared relatively consistent among 89.6, JR-CSF, and CHO40-exposed neurospheres. Collectively, this data shows that HIV-1 viral replication occurs in NPC-derived neurospheres in a reproducible manner.

We were also interested in examining the potential tropism of HIV-1 in neurospheres. While CD4^+^T cells are the most abundantly HIV-1 infected cells in the body there is ample evidence to suggest that microglia and astrocytes are not only capable of being infected by HIV-1, but may also serve as viral reservoirs, as reported elsewhere [56, 57, 58, 59, 60, 61, 62]. For this experiment, NPCs were gently disaggregated with trypsin and immunoprecipitation (IP) was performed using antibodies for either astrocytes (GFAP^+^), microglia-like cells (CD11b^+^), mature neurons (GAD65^+^), or NSCs (SOX2^+^). We next performed direct PCR of the IP material without purification of DNA, as RNA isolation from a relatively small number of infected cells would not yield a sufficient amount of sample material for RT-qPCR. Results from qPCR confirmed the presence of TAR DNA in microglia-like cells in the 89.6, JR-CSF, and CHO40 infected neurospheres (Fig. 2d). While the highest numbers of TAR DNA were associated with astrocytes (∼480 copies) and microglia-like cells (∼780 copies) in the JR-CSF samples, lower amounts were also detected in microglia-like cells from the 89.6 (∼ 114 copies) and CHO40 (∼ 125 copies) samples. In addition, astrocytes from the JR-CSF neurospheres (∼480 copies) had substantially higher levels of TAR DNA than astrocytes from the 89.6 and CHO40-infected neurospheres (< 20 copies each). However, within the JR-CSF samples, TAR DNA was still the most abundant in microglia-like cells. Consistent with the literature, this data suggests that HIV-1 tropism in neurospheres is directed towards glial cells [56, 57, 58, 59, 60, 61, 62].

Based on this data, and given that microglia are the prominent innate immune cells in the CNS and, therefore, represent a major target of HIV-1, we next attempted to increase the population of myeloid cells in neurospheres to better assess the replication potential of HIV-1. To this end, differentiated neurospheres were treated with the following cytokines: IL-34 (100 ng/mL), M-CSF (25 ng/mL), TGFβ-1 (50 ng/mL) every other day for a period of seven days, as previous studies have demonstrated the ability of these cytokines to induce microglia differentiation in iPSCs [16, 63, 64]. Neurospheres were then infected with HIV-1 dual tropic 89.6 (MOI:10) with or without cART as described above. After fourteen days, supernatants were collected and neurospheres were harvested for downstream assays. As shown in Fig. 2e, western blot revealed robust expression of p24 capsid protein with undetectable expression of Pr55, thus suggesting an efficient cleavage of the viral polyprotein in the induced neurospheres relative to the uninfected sample. We also observed increased expression of the intracellular forms (e.g. 30, 40 kDa) of the accessory protein Nef [65] relative to the uninfected sample. Again, treatment of HIV-1 neurospheres with cART decreased the expression of both p24 and Nef.

Cytokine-treated neurospheres were also assessed via western blot to evaluate the expression of different microglia markers (e.g. IBA-1, CD11b, CD45). Data in Fig. 2e indicates that IBA-1 expression was relatively consistent among each sample, whereas slightly varying levels of expression of CD11b and CD45 were observed between samples. Additionally, the expression levels of apoptotic proteins were evaluated to assess whether cytokine treatment or cART had any potential effects on apoptosis (Fig. 2e). Here, expression of the pro-apoptotic protein BAD increased after HIV-1 infection and was reduced in the presence of cART. Importantly, BAD expression was undetectable in the uninfected and cART-only samples. While low levels of uncleaved PARP-1 were observed in the untreated and cART samples, we observed increased expression of the 24 kDa cleaved fragment in the HIV-1 sample.

RT-qPCR for HIV-1 TAR and *env* confirmed the expression of these RNAs in the HIV-1 cytokine-treated samples. Of note, similar amounts of both TAR and *env* (∼ 10^8^ copies each) were detected, suggesting that increased viral processivity was occurring in these cultures (Fig. 2f). Again, treatment with cART resulted in significant reductions of both TAR and *env* RNA relative to the HIV-1 sample. Collectively, this data suggests that treatment of neurospheres with IL-34, M-CSF, and TGFβ-1 effectively enhanced viral replication and processivity and that this may be attributed to the induction of a pro-myeloid phenotype. This data also indicates that cytokine treatment alone, as well as cART alone, did not promote apoptosis in neurospheres.

We next sought to assess the potential effects of HIV-1 (89.6) on the expression levels of multiple proteins or enzymes associated with different CNS cell types. To this end, western blot was performed to evaluate the relative expression microglia-like cells (CD11b^+^, CD163^+^), astrocytes (GFAP^+^, glutamine synthetase^+^), dopaminergic neurons (TH^+^, FOXA2^+^), GABAergic neurons (GAD65^+^, GAD67^+^), glutamatergic neurons (BNPI^+^) and NSCs (SOX2^+^). As shown in Fig. 3a, HIV-1 appeared to increase the expression of differentiated neurons, particularly TH^+^, GAD65^+^/GAD67^+^, and BNPI^+^ neurons, whereas treatment with cART restored the expression of these markers to levels that were comparable to the uninfected sample. A similar pattern of expression was also observed for GFAP, while the expression of CD11b appeared relatively consistent between the uninfected, HIV-1 infected, and cART samples. On the other hand, expression of CD163 appeared to slightly decrease in both the HIV-1 and cART samples relative to the uninfected sample.

**Figure 3.**
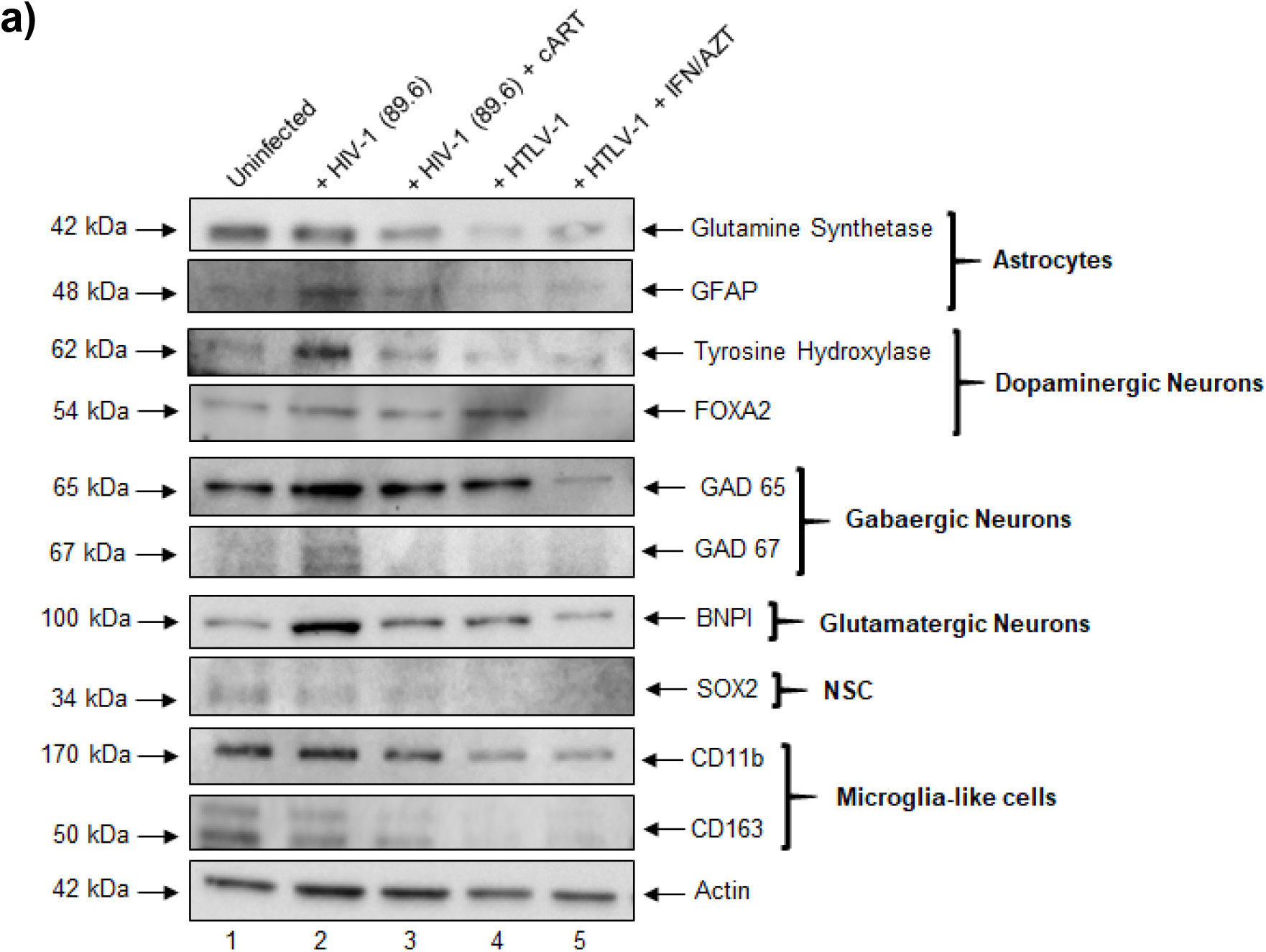
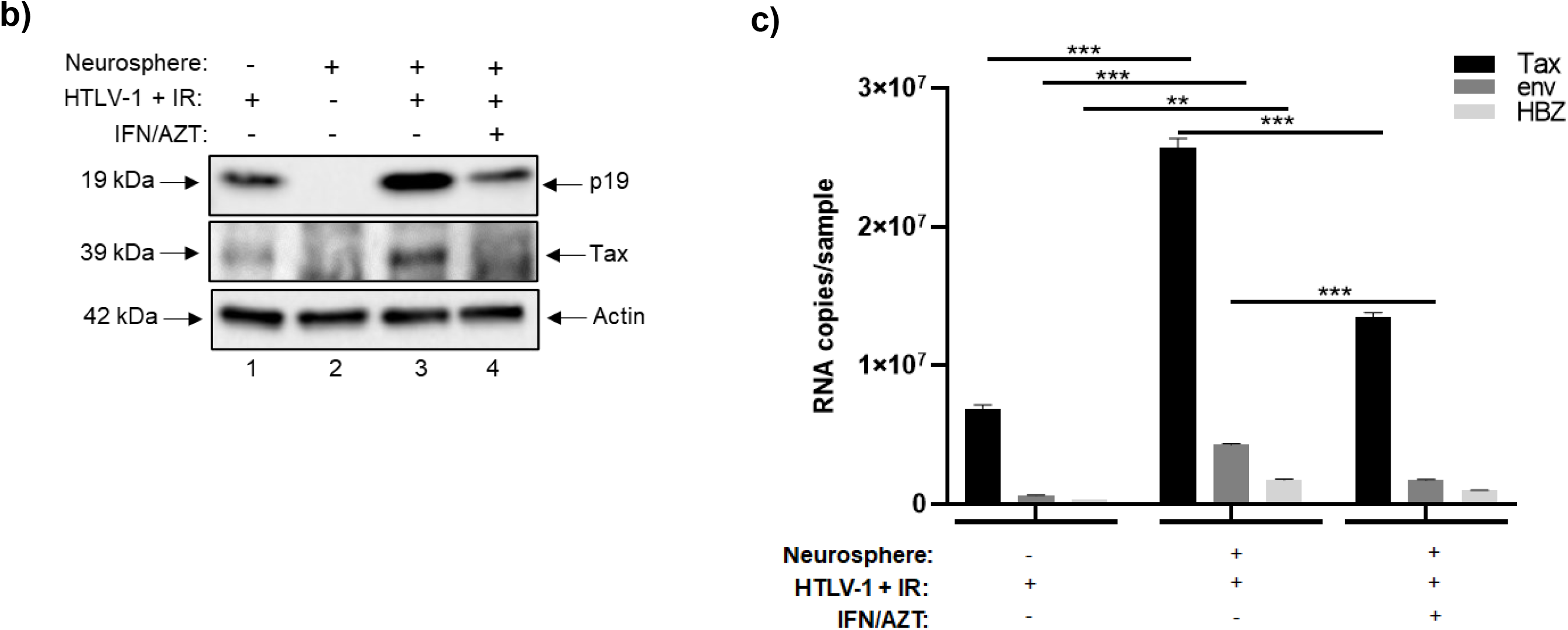
HTLV-1 co-culture and expression of cellular markers in HIV-1 and HTLV-1 infected neurospheres. **(a)** Differentiated neurospheres were exposed to dual-tropic HIV-1 89.6 with or without cART (lamivudine, tenofovir disoproxil fumarate, emtricitabine, indinavir), or were co-cultured with irradiated HTLV-1 infected cells with or without IFN/AZT for a period of fourteen days. Western blot was performed to examine the relative expression of various CNS-relevant cellular markers including those associated with microglia-like cells, astrocytes, neurons (dopaminergic, GABAergic, glutamatergic), and NSCs. **(b)** Western blot was performed to examine the relative expression of Tax and p19 in neurospheres co-cultured with HTLV-1 infected cells with or without IFN/AZT treatment. **(c)** RT-qPCR was performed to quantify the copy numbers of *tax*, *env*, and *hbz* RNA in neurospheres co-cultured with HTLV-1 infected cells with or without IFN/AZT treatment. n=3. ** p < 0.005, *** p < 0.0001.

As an additional control, we also evaluated the effects of co-culturing neurospheres with the HTLV-1 infected cell line HUT 102. HTLV-1 is the etiologic agent of HTLV-1-associated myelopathy/tropical spastic paraparesis (HAM/TSP), a neurological disorder that is characterized by chronic neuroinflammation and neurodegeneration [66, 67, 68, 69]. Prior to initiating the co-culture, HUT 102 cells were exposed to ionizing radiation (IR) to inhibit chromosomal replication and to activate viral transcription, as previously described [70, 71, 72]. Treatment with a combination of IFN/AZT was also included to model the current standard for HTLV-1 anti-viral therapy [73, 74, 75]. The irradiated HUT 102 cells served as the background control. After fourteen days of co-culture, neurospheres were harvested for downstream assays. Data in Fig. 3a shows that HTLV-1 co-culture resulted in a slight increase in the expression of FOXA2 and BNPI neurons and that treatment with IFN/AZT reversed this increase. On the other hand, HTLV-1 appeared to reduce the expression of both CD11b and glutamine synthetase relative to the uninfected sample, whereas IFN/AZT did not appear to have any impact. While the expression of NSCs appeared relatively faint in the uninfected and HIV-1 samples, its expression was undetectable in the HTLV-1 samples. These results are not entirely surprising as neurospheres were cultured in medium that was enriched to induce differentiation into more mature cell types.

To confirm whether HTLV-1 was capable of replicating in neurospheres, western blot was performed to assess the expression of HTLV-1 Tax and matrix (p19) proteins. As shown in Fig. 3b, the expression of both Tax and p19 was higher after co-culturing of neurospheres with irradiated HUT 102 cells (relative to the background control), thereby indicative of increased viral replication. As expected, treatment with IFN/AZT reduced the expression of Tax and p19 to levels that were similar to the background control. Furthermore, RT-qPCR for HTLV-1 *tax*, *env,* and *hbz* RNA showed that the expression of these RNAs significantly increased after co-culture and that treatment with IFN/AZT resulted in a significant reduction of both *tax* and *env* transcripts (Fig. 3c).

Overall, these results demonstrate the ability of HIV-1 to replicate in NPC-derived neurospheres, therefore indicating that these models may be permissive to infection. Moreover, our data suggests that the exposure of neurospheres to microglia-inducing cytokines may enhance their susceptibility to infection. While the primary focus of this manuscript remains on HIV-1, the data generated from the HTLV-1 experiments confirms the susceptibility of neurospheres to retroviral infection and lays the foundation for future experiments focused on HTLV-1 or other retroviral infections. Lastly, this data also points towards the differential effects of viral infection on the cellular composition of neurospheres. Since neurospheres are composed of a heterogeneous population of cell types, including those that are infected and uninfected, this area of research deserves further attention.

### The Effect of Stem Cell EVs on HIV-1 Infected Neurospheres

Our experiments have shown receptor mediated alterations post-infection, indicating cellular damage. Therefore, there is a need for a “holistic” reparative approach to potentially control cellular damage and regulate inflammation. There is mounting evidence that EVs from different sources of stem cells, namely MSCs and iPSCs, possess broad reparative and regenerative properties across various pathologies, including those related to the CNS [43, 44, 45, 46, 47, 48]. Whether or not these vesicles confer reparative effects in the context of infection remains relatively unexplored, especially with respect to the CNS. Our previous research pointed towards the potential reparative effects of both MSC and iPSC EVs in 2D cultures of neurons, astrocytes, and monocyte-derived macrophages (MDMs) damaged by IR [76]. Therefore, we sought to expand upon these studies by investigating the potential functional effects of MSC and iPSC EVs in HIV-1 infected neurospheres.

The EVs used for these experiments were isolated using a scalable platform which incorporated the use of tangential flow filtration (TFF) [76]. This advanced technology has the ability to produce large batches of highly concentrated material in an efficient and reproducible manner and in recent years its application for EV isolation has been well documented by others [77, 78, 79, 80]. Here, we include data from the analysis of MSC and iPSC EVs; we also include data from A549 (cancer) EVs which served as both a control for our EV isolation method as well as a non-stem cell control. NTA analysis shows that the size distribution of each EV preparation falls within the range of 50 to 250 nm (Fig. 4a). Consistent with our previous research, iPSC EVs and A549 EVs appear more heterogeneous in their size distribution relative to MSC EVs [76]. TEM of each EV preparation also confirmed the presence of cup-shaped vesicles that appeared less than 200 nm in diameter. Representative TEM images from each EV preparation are included in Fig. 4b. Interestingly, these images also showed that iPSC EVs tended to cluster together in aggregates of smaller (∼50 nm) vesicles whereas MSC EVs appeared as both larger (∼100 nm) vesicles and aggregates of smaller (∼50 nm) vesicles.

**Figure 4.**
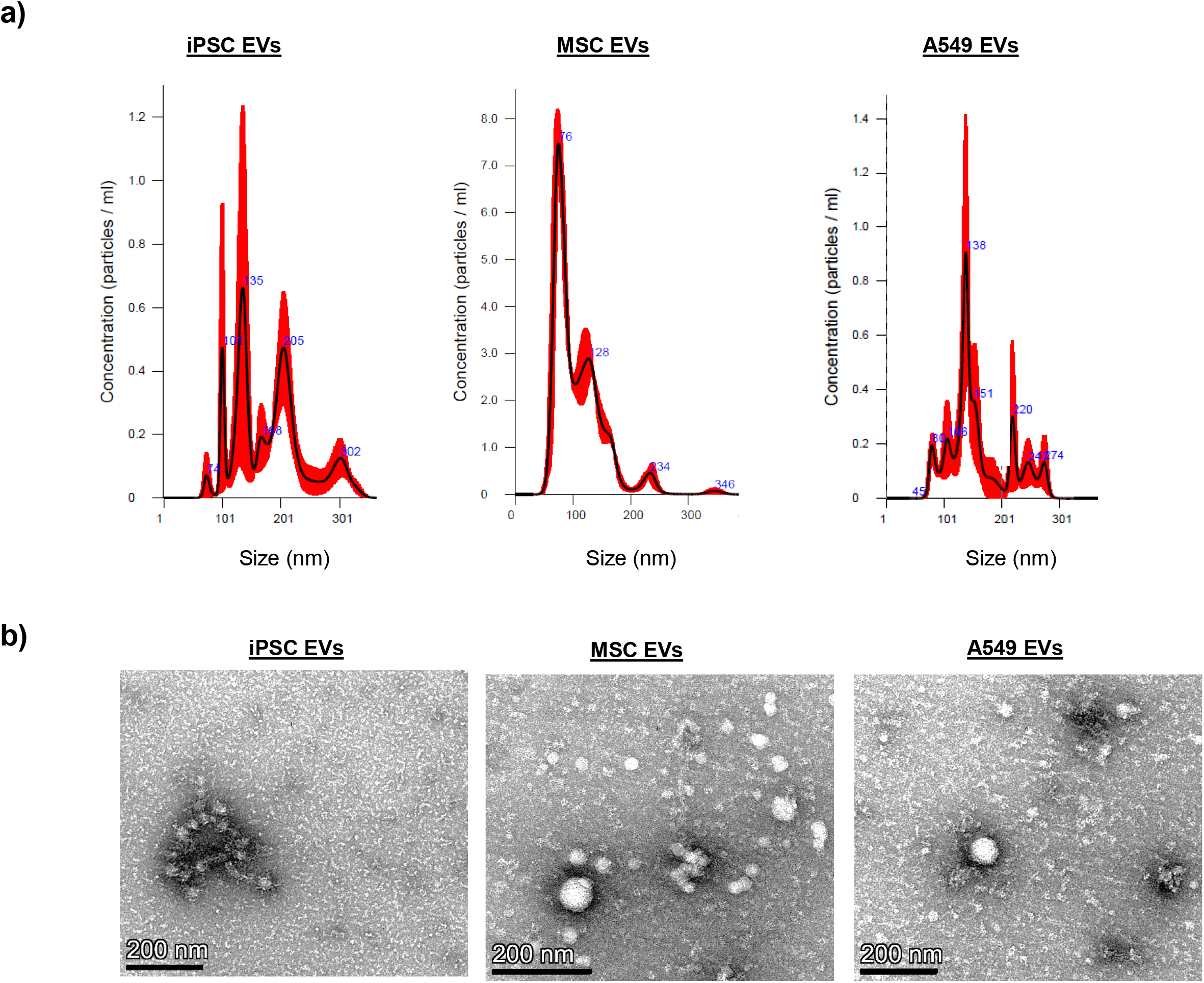
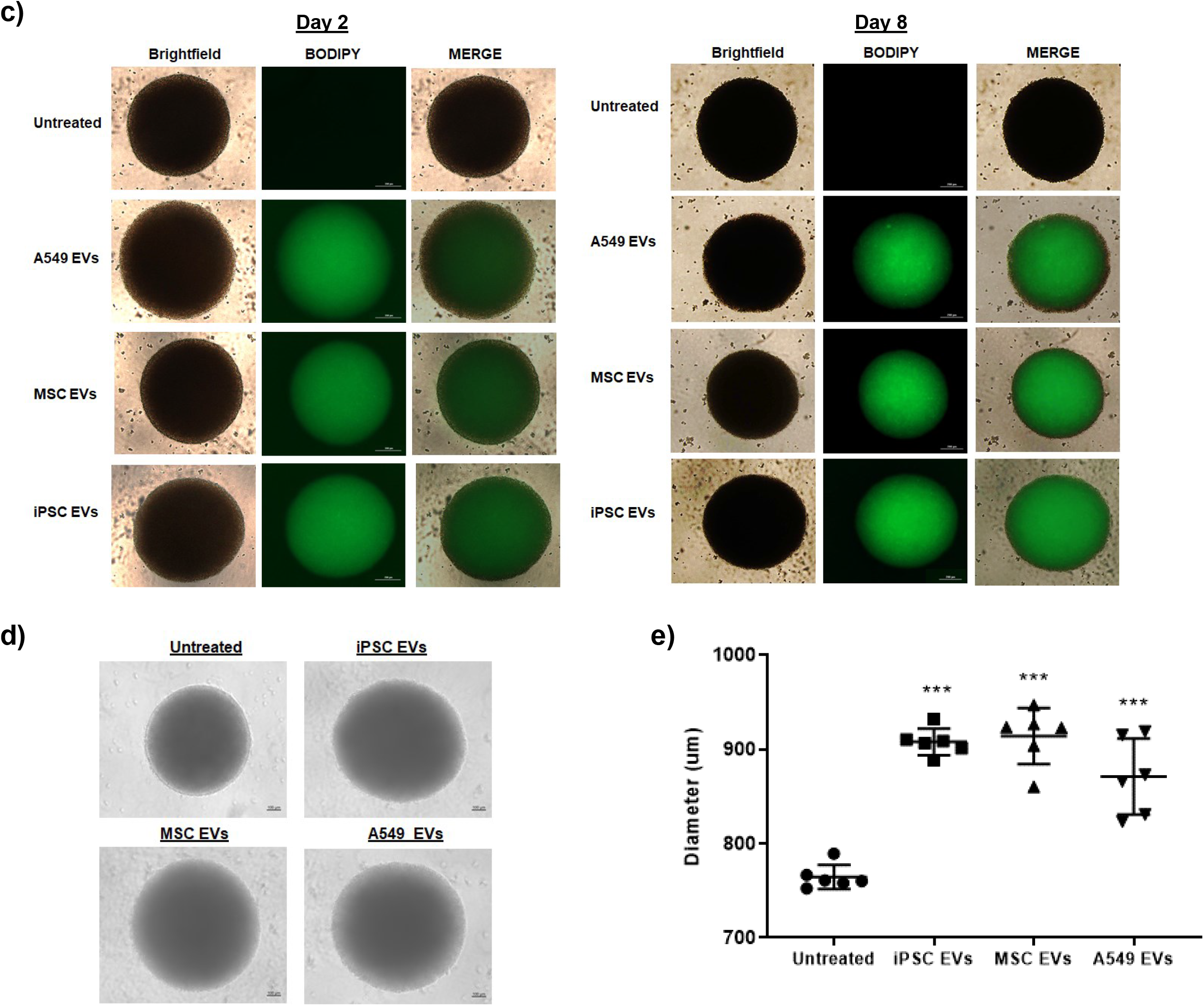
EV characterization and uptake assay. EVs from stem cells (MSCs, iPSCs) as well as A549 cancer cells were concentrated using TFF. Data from A549 EVs is included to serve as a control for NTA analysis and EV uptake. **(a)** NTA was performed and representative histograms from three independent measurements show the relative size distribution of each EV prep. **(b)** TEM (negative staining method). EVs were fixed with 4% glutaraldehyde in 0.12 M sodium cacodylate buffer and stained with 1% aqueous uranyl acetate. Imaging was performed using an FEI Talos F200X transmission electron microscope at 80KV. Scale bar = 200 nm. **(c)** EVs were fluorescently labeled with BODIPY 493/503. After removal of excess dye, EVs were added to differentiated neurospheres (day 0) at an approximate ratio of 1:250 (recipient cell to EV ratio). After 24 hours the media was completely replaced. Representative brightfield and fluorescent images show the relative uptake of EVs on days 2 and 8. Scale bar = 200 µm. n= 3. **(d)** Phase contrast images show the relative size and morphology of EV-treated neurospheres after two weeks in culture. Scale bar = 100 µm. n = 3. **(e)** The average diameter (two measurements per sphere) of EV-treated neurospheres was measured using calibrated imaging software. n = 3 neurospheres. *** p < 0.0001 relative to untreated.

To assess the efficiency of their uptake by differentiated neurospheres, EVs were fluorescently labeled with a commercially available dye which stains both neutral and nonpolar lipids (BODIPY). Labeled EVs were then added to cultures of differentiated neurospheres. As shown in Fig. 4c, fluorescent microscopy initially revealed a relatively weak and diffuse signal that persisted throughout the first 48 hours (day 2). However; over time, the fluorescent intensity increased. Representative photographs (day 8) show a strong fluorescent signal that appeared to be predominantly localized in the center of each EV-treated neurosphere. We also observed similar results when treating neurospheres with EVs labeled with the SYTO RNASelect nucleic acid stain (Supplementary Fig. 2a). Additionally, image analysis of EV-treated, cross-sectioned neurospheres pointed towards a strong and relatively even EV uptake, with fluorescent signal being detected in cells throughout the center and periphery of the spheres (Supplementary Fig. 2b). Interestingly, we also observed a change in the diameter and morphology of EV-treated neurospheres over time. Data in Fig. 4d shows representative images taken two weeks-post EV treatment. Here, EV-treated neurospheres appeared larger in size and the borders were relatively less well-defined as compared to the untreated neurospheres. Quantification of neurosphere diameter confirmed that EV-treated neurospheres were significantly larger in size relative to untreated neurospheres (Fig. 4e and Supplementary Figure 2c).

Given our observation that EV-treated neurospheres displayed a change in morphology that was characterized by a slight reduction in their well-defined spherical border, we next questioned whether EVs may contain enzymes capable of degrading the extracellular matrix within neurospheres. To address this, we performed western blot for matrix metalloproteinase 9 (MMP9), as it is one of the most well-studied members of the MMP family and, interestingly, has also been associated with reparative and regenerative functions in the CNS [81, 82]. Results from this experiment revealed MMP9 expression in both iPSC and A549 EVs, while its expression was undetected in MSC EVs (Supplementary Fig. 2d). As a follow-up, we performed gelatin zymography, as this is a reliable assay to assess the relative proteolytic activity of experimental samples [83, 84]. Imaging of the stained gel indicated the presence of active MMP9 as well as active MMP2 in each EV prep. Among the samples A549 EVs displayed the highest intensity, followed by iPSC and MSC EVs (Supplementary Fig. 2e). This data suggests that EVs are capable of not only penetrating, but also persisting in, three-dimensional cultures and, moreover, that EV-associated proteolytic enzymes may partially contribute to this effect.

After confirming EV uptake by neurospheres we focused next to examine their potential functional effects in HIV-1 infected neurospheres. To date, the anti-apoptotic effects of stem cell EVs in the context of the CNS have been well documented both *in vitro* and *in vivo* [85, 86, 87, 88, 89, 90]. To this end, we performed western blot to examine the expression levels of different pro-apoptotic proteins (Fig. 5a). Relative to the uninfected control sample, HIV-1 was associated with a slight increase in the expression of cleaved PARP-1 as well as a moderate increase in the expression of both Caspase 3 and BAD. Furthermore, the addition of cART was associated with an increased expression of cleaved Caspase 3 and BAD relative to the HIV-1 sample. However, the addition of cART in combination with EVs (iPSC, MSC EVs, and A549) decreased the expression of cleaved PARP-1 (24 kDa), Caspase 3 (cleaved and uncleaved), and BAD relative to HIV-1 + cART. As an additional control, we included treatment of uninfected neurospheres with cART (lane 3). The expression of each protein in this sample was relatively consistent with those of the uninfected control, suggesting that cART alone did not have a toxic effect on the neurospheres.

**Figure 5.**
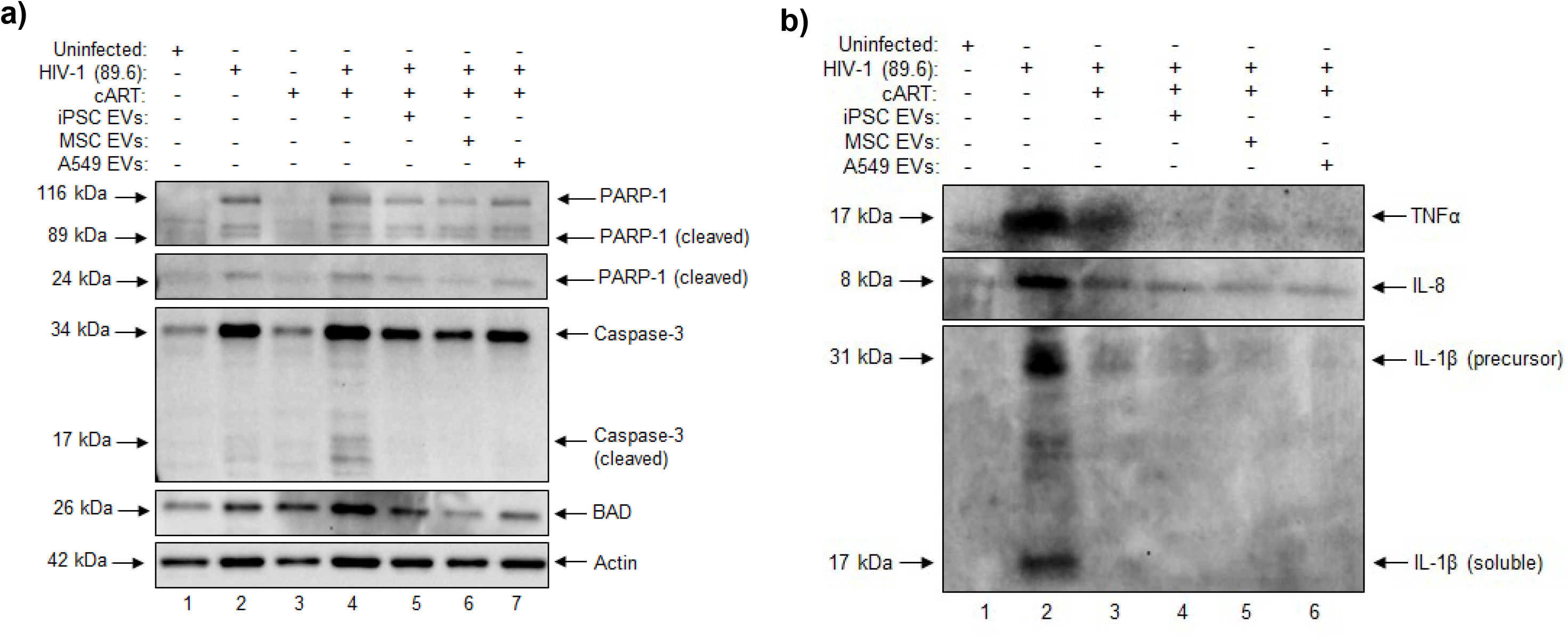
The effect of EVs on HIV-1 infected neurospheres. Differentiated neurospheres were exposed to dual-tropic HIV-1 89.6 with or without cART (lamivudine, tenofovir disoproxil fumarate, emtricitabine, indinavir) for a period of fourteen days. Additionally, the HIV-1 + cART treated samples were treated with either iPSC, MSC, or A549 EVs at an approximate ratio of 1:250 (recipient cell to EV ratio). **(a)** Western blot was performed on neurosphere lysates to evaluate the relative expression of PARP-1, Caspase-3, and BAD. **(b)** Western blot was performed on neurosphere supernatants to evaluate the relative expression of TNFα, IL-8, and IL-1β.

In addition to their anti-apoptotic effects, stem cell EVs are also associated with potent anti-inflammatory properties and several recent studies have highlighted their ability to reduce neuroinflammation in various models of CNS damage [91, 92, 93, 94, 95, 96]. To assess the potential anti-inflammatory effects of stem cell EVs on HIV-1 infected neurospheres, we performed western blot for well-known pro-inflammatory mediators (Fig. 5b). Here, substantial increases in TNFα, IL-8, and IL-1β were observed after HIV-1 infection. Treatment with cART decreased expression of soluble IL-1β to an undetectable level while only slightly decreasing the expression of TNFα and IL-8. Notably, similar results were also observed in JR-CSF and CHO40-infected neurospheres (Supplementary Fig. 3). Again, in HIV-1 89.6-infected neurospheres, the combination of cART with either iPSC or MSC EVs restored the expression of TNFα, IL-8 and IL-1β to levels that were comparable to the uninfected control. Interestingly, similar results were observed with A549 EVs. Taken together, this data indicates that EVs are capable of exerting both neuroprotective and anti-inflammatory effects in HIV-1 infected neurospheres. Overall, these findings are in consensus with the pre-existing literature on stem cell-derived EVs. Given that A549 EVs originate from cancer cells, a more thorough assessment of the functional effects associated with cancer EVs is required to evaluate the long-term consequences in recipient cells.

### RNA Analysis of Stem Cell EVs

The reparative aspects of stem cell EVs have been shown to be associated with various non-coding RNAs and/or proteins that confer protective and/or immunomodulatory functions in damaged recipient cells [97, 98, 99, 100, 101]. While several recent studies have extensively characterized the miRNA profiles of stem cell EVs, the characterization of EV-associated long non-coding RNAs (lncRNAs) remains a relatively unexplored topic [41, 102, 103, 104]. Our previous studies included the sequencing of lncRNAs (≥ 250 base pairs) from iPSC, MSC, and A549 EVs [76]. Again, the purpose of including A549 EVs was to serve as a non-stem cell control. Here, we share an expanded set of data from our initial studies to highlight the differences that exist among these different EV populations.

Overall, approximately 6.5 x10^7^, 1.9 x 10^7^, and 2.6 x 10^7^ valid reads were obtained from iPSC, MSC, and A549 EVs, respectively, with each preparation having a Q30 score greater than 85% (Supplementary Table 1). Of these reads, approximately 80.88% (iPSC EVs), 71.21% (MSC EVs), and 70.66 (A549 EVs) were mapped to the genome. As shown in Fig. 6a, the majority of lncRNAs among each EV preparation mapped to introns. iPSC EVs had the highest number of intronic transcripts (93%) followed by MSC EVs (79%) and A549 EVs (75%). The percentage of transcripts mapping to either exons or intergenic regions ranged from approximately 10 to 15% among MSC and A549 EVs; however only ∼3% of transcripts from iPSC EVs were mapped to these regions.

**Figure 6.**
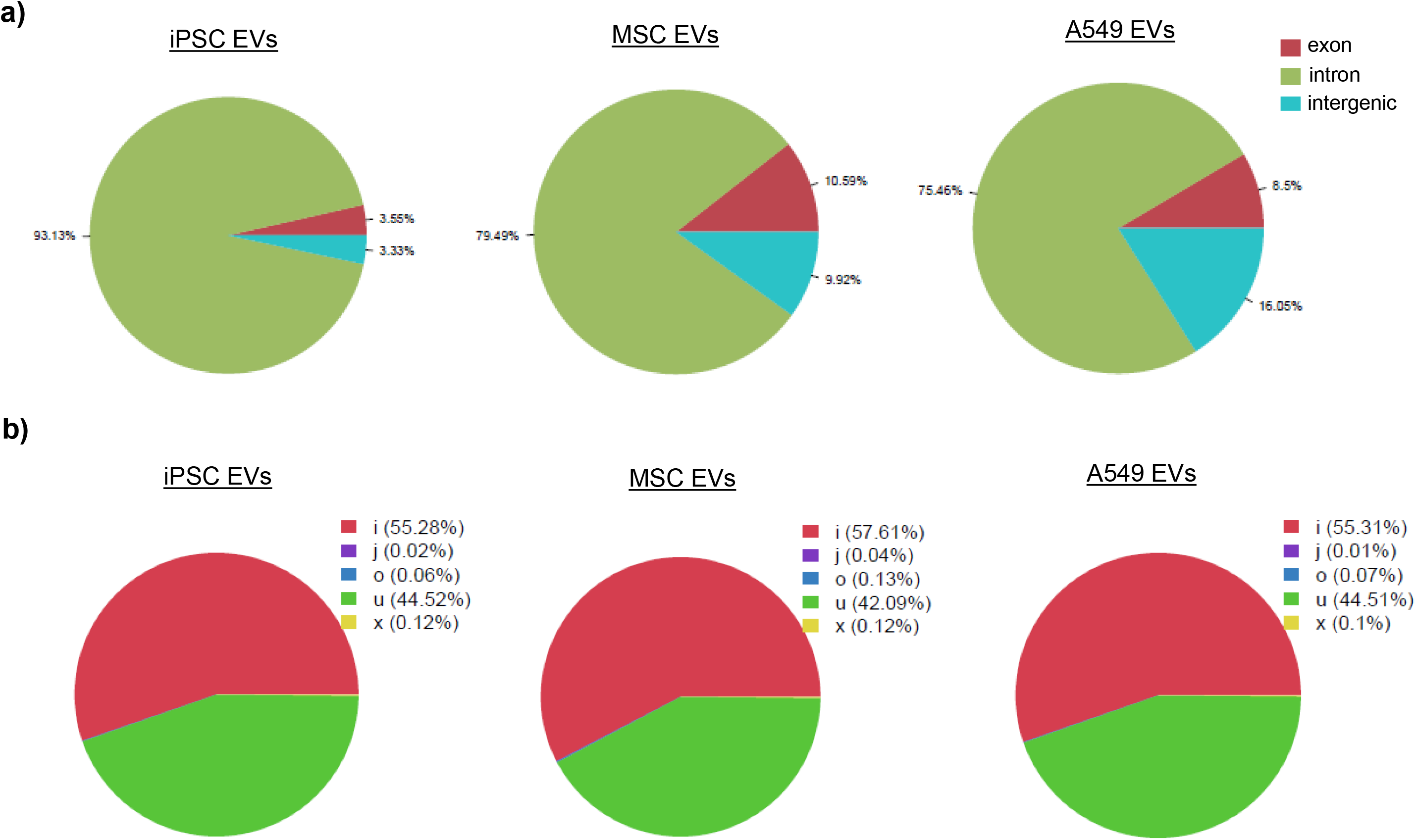
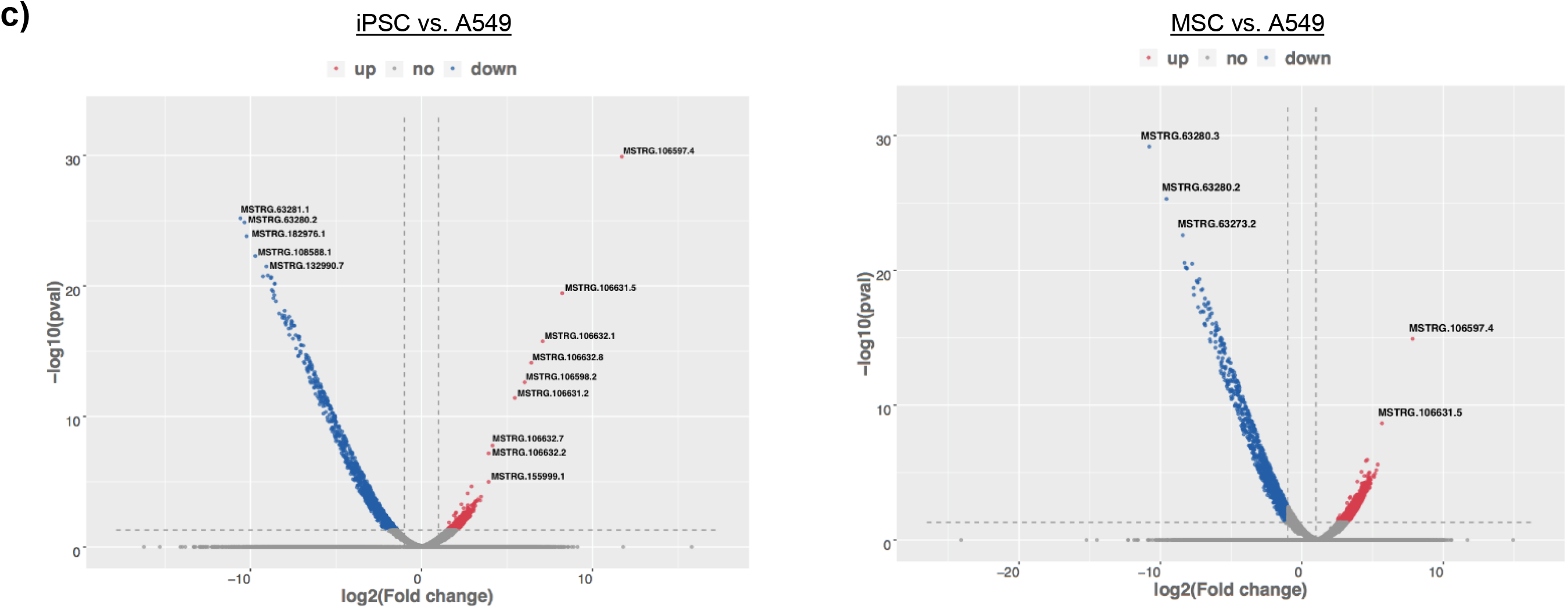
RNA analysis of EVs. RNA sequencing of EV-associated lncRNAs was performed. **(a)** Percentages of transcripts mapping to either exons, introns, or intergenic regions are shown. **(b)** Percentages of non-annotated transcripts. Transcripts not annotated in genome databases were designated as novel transcripts according to the following class codes: i = transfrag falling within a reference intron; j = potentially novel isoform with at least one splice junction shared with a reference transcript; o = generic exonic overlap with a reference transcript; u = unknown intergenic transcript; x = exonic overlap with reference on the opposite strand. **(c)** Volcano plots displaying the differentially expressed transcripts between iPSC and A549 EVs (left panel) and MSC and A549 EVs (right panel). Significantly regulated transcripts are annotated with their corresponding gene ID.

Transcripts that were not annotated in genome annotation databases were subsequently designated as novel transcripts according to the following class codes: i = transfrag falling within a reference intron; j = potentially novel isoform with at least one splice junction shared with a reference transcript; o = generic exonic overlap with a reference transcript; u = unknown intergenic transcript; x = exonic overlap with reference on the opposite strand. Data in Fig. 6b shows that within each EV prep, the majority of transcripts either fell within a reference intron (i) (> 50%) or were classified as unknown intergenic transcripts (u) (< 40%) and, moreover, that ≤ 0.1% of transcripts among each sample were classified as either “j”, “o”, or “x”.

Volcano plots in Fig. 6c provide a visual representation of the differentially expressed transcripts between iPSC and A549 EVs (Fig. 6c, left panel) and between MSC and A549 EVs (Fig. 6c, right panel). Transcripts displaying statistically significant regulation (either up or down) are annotated with their associated gene ID/transcript name. Overall, these plots reveal that a majority of transcripts associated with stem cell EVs are downregulated relative to A549 EVs. Additionally, iPSC EVs displayed a greater amount of significantly up-regulated transcripts relative to MSC EVs when compared to A549 EVs. Notably, transcript MSTRG.106597 was among the most significantly upregulated transcript associated with both iPSC EVs and MSC EVs and its approximate fold change relative to A549 EVs was 3392 for iPSC EVs and 229 for MSC EVs. For reference, this is designated as a novel transcript which corresponds to gene FP671120.1. Heat maps generated from all EV-associated lncRNA transcripts are included in Supplementary Fig. 4a and show the comparisons between iPSC and A549 EVs (left panel) and between MSC and A549 EVs (right panel). Annotated transcripts were further analyzed by performing Gene ontology (GO) (Supplementary Fig. 4b-c). Relative to A549 EVs, iPSC EVs had significantly enriched GO terms which included RNA and protein binding, positive regulation of GTPase activity and cell adhesion. Relative to A549 EVs, MSC EVs had significantly enriched GO terms which included RNA binding, protein binding, and T cell receptor signaling pathway. Chromosome maps of EV-associated lncRNA transcripts revealed striking differences in the density distribution when comparing stem cell EVs and A549 EVs. Specifically, transcripts from A549 EVs appeared more evenly distributed and spanned all chromosomes while transcripts from iPSC and MSC EVs only mapped to certain chromosomal regions (Supplementary Fig. 4d).

The heatmap comparing the expression of all lncRNA transcripts between iPSC and MSC EVs is shown in Supplementary Fig. 5a and the corresponding volcano plot is show in Supplementary Fig. 5b. Of note, this graph shows a relatively even distribution of up- and down-regulated transcripts. Collectively, this data reveals distinct differences in the expression of lncRNA transcripts associated with EVs derived from stem cells versus cancer (A549) cells. Future studies will include a more thorough assessment of the differentially expressed transcripts among iPSC and MSC EVs and will focus on their potential functional implications.

### Detection of RNA Binding Proteins Using RNA Pull-down Assay

In an effort to define the role of EV-associated lncRNAs and to better understand their potential functional significance in recipient cells, we recently identified at least one lncRNA transcript associated with iPSC EVs (AC120498.9) which possessed a unique secondary structure [76]. Specifically, this transcript contained two continuous stretches of double-stranded RNA that were greater than 30 base pairs (bp) in length, a property that has previously been associated with the activation of PKR [105, 106]. This prompted us to further examine the secondary structures of other EV-associated lncRNAs, first by focusing on those that were more highly associated with iPSC EVs. We subsequently generated secondary structures from over 240 iPSC EV-associated lncRNAs that had FPKM values of ≥ 2 relative to MSC and A549 EVs. From this analysis we identified another transcript (ADIRF-AS1) which also contained a continuous stretch of double-stranded RNA greater than 30 bp. For reference, the secondary structures of ADIRF-AS1 and AC120498.9 are shown in Fig. 7a. RT-qPCR confirmed that both of the relevant sequences shown in Fig. 7a (i.e. double-stranded RNA greater than 30 base pairs) were detected in EVs and that their expression was significantly increased in iPSC EVs relative to MSC EVs (Fig. 7b).

**Figure 7.**
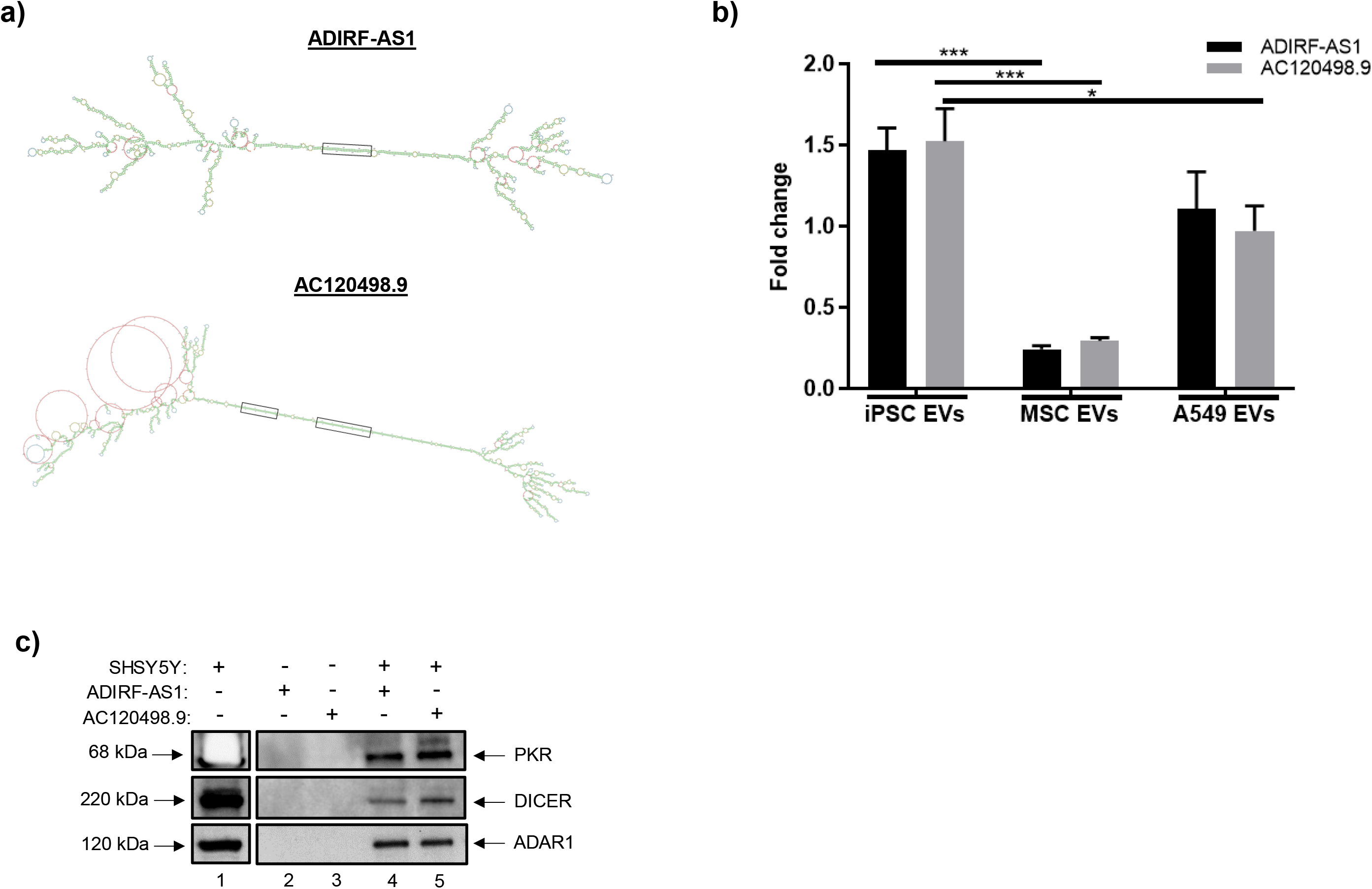
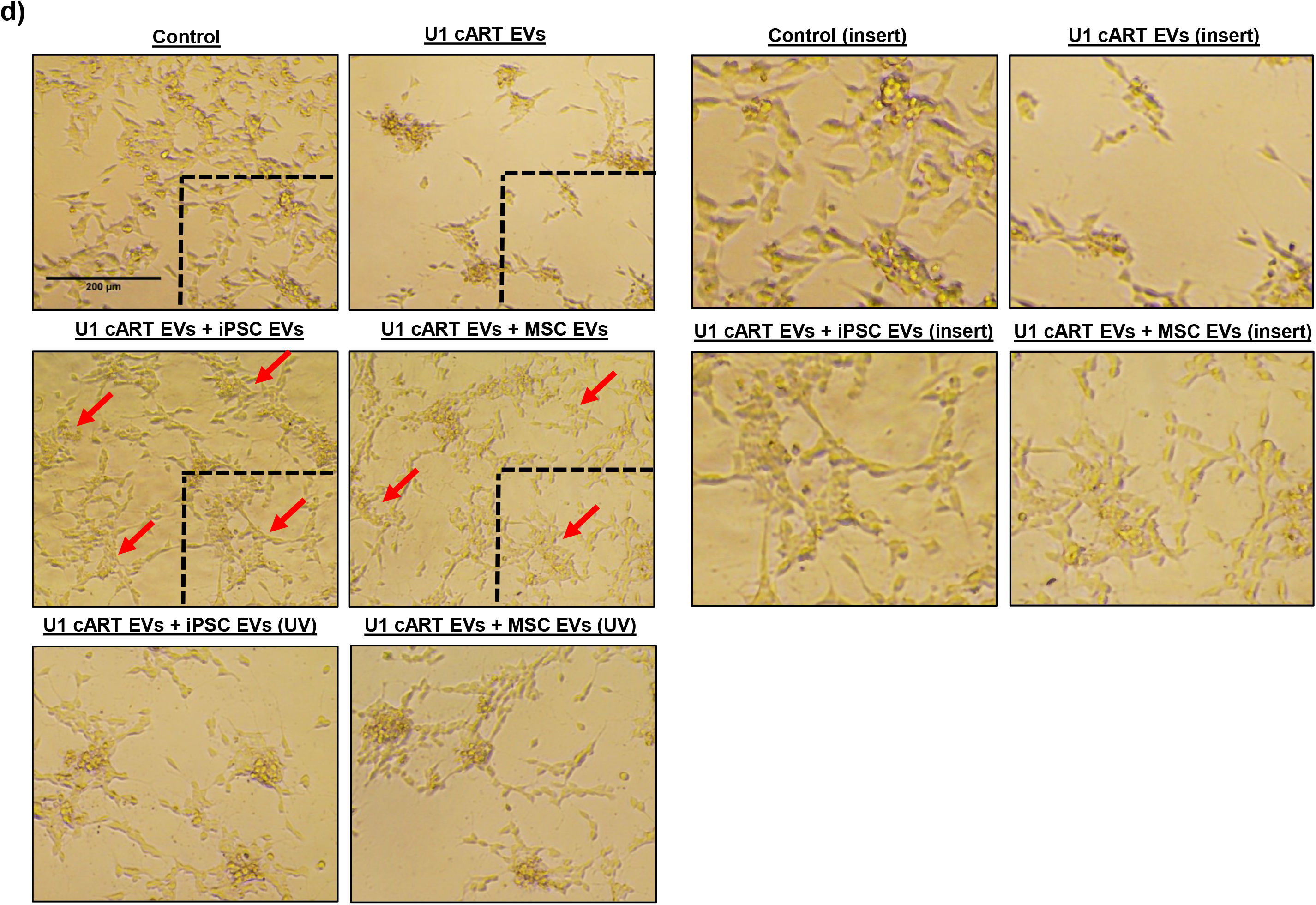
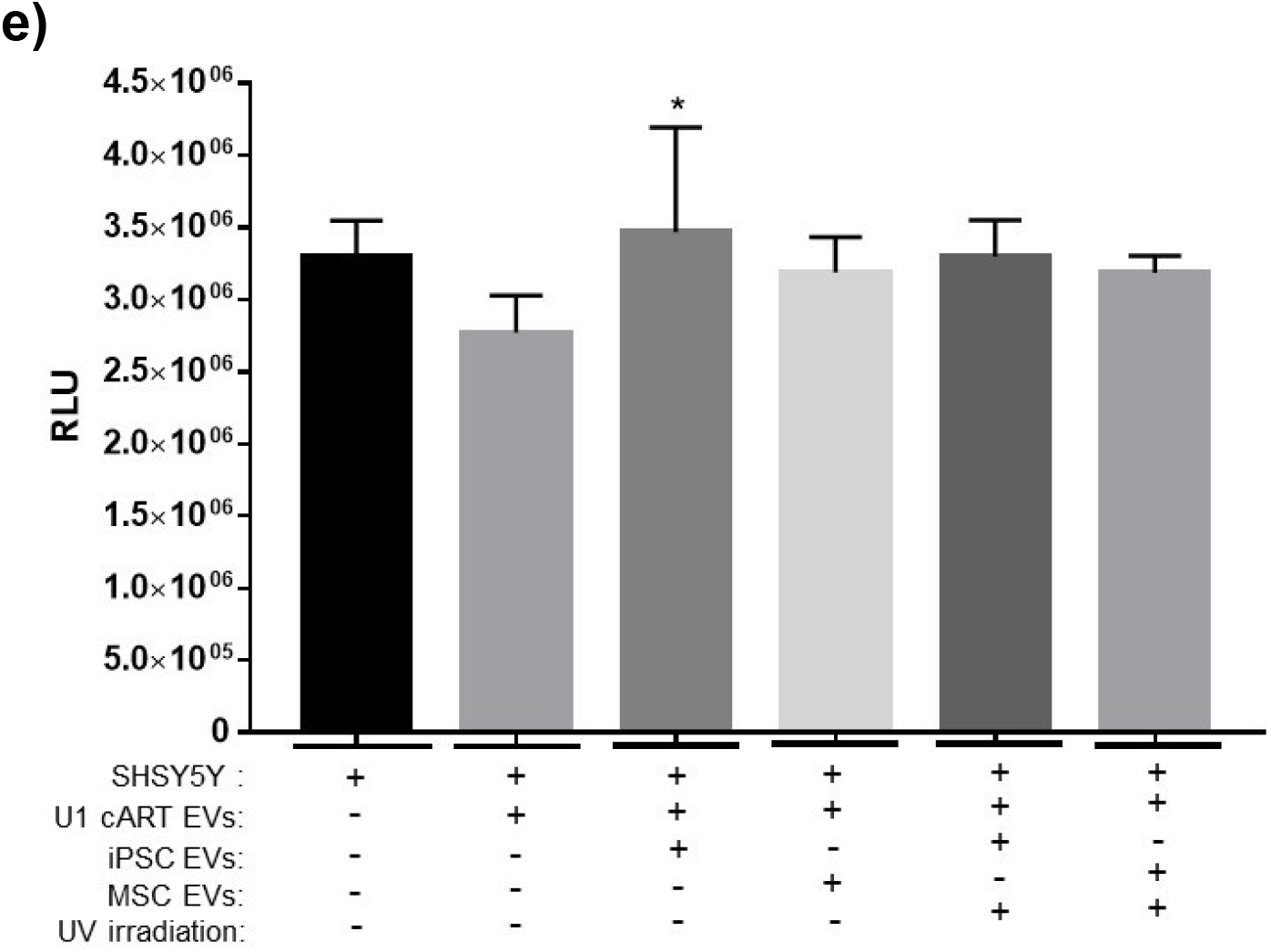
RNA pulldown assay. **(a)** The secondary structures of lncRNAs ADIRF-AS1 and AC120498.9 are shown. These lncRNAs contained continuous stretches of dsRNA ≥ 30 base pairs in length; for reference, the corresponding regions are enclosed in black boxes. **(b)** Total RNA was isolated from EVs and RT-qPCR was carried out using primers that were specific to the boxed sequences shown in panel a. Data were normalized to GAPDH and the relative expression levels were evaluated using the 2^−ΔΔCT^ method. n = 3. * p < 0.05, *** p < 0.0001. **(c)** Biotin-conjugated synthetic RNA sequences corresponding to the sequences of interest were incubated with SHSY5Ywhole cell extract. Western blot was performed to assess the expression of different RNA binding proteins (PKR, Dicer, ADAR1). **(d)** SHSY5Y cells were exposed to EVs from cART-treated, HIV-1 infected cells (U1) with or without treatment of stem cell EVs at an approximate ratio of 1:1000 (recipient cell to EV ratio). Stem cell EVs were exposed to UV(C) to inactivate EV-associated RNAs. Representative images show the appearance and morphology of cells after 8 days in culture. Inserts are included for higher resolution. Scale bar = 200 µm. n = 3. **(e)** Cell viability was quantified via CellTiter-Glo. n=3. * p < 0.05 relative to U1 cART EVs.

To assess whether the identified sequences were capable of binding to PKR the biotin-conjugated synthetic RNA sequences were incubated with whole cell lysate from the neuronal cell line SHSY5Y and then purified using Streptavidin-Sepharose beads. Western blot was performed to assess the expression of PKR, as well as other critical RNA binding proteins. The results in Fig. 7c confirm that the relevant sequences from both ADIRF-AS1 and AC120498.9 were capable of binding not only to PKR, but also to other double stranded RNA (dsRNA) binding proteins including DICER and ADAR1. We additionally performed similar experiments using whole cell lysate from either astrocytes or MDMs (Supplementary Fig. 6). Again, western blot analysis confirmed expression of PKR, DICER, and RIG-I, thereby highlighting the potential of these RNAs to bind to multiple different RNA binding proteins in CNS relevant cell types. As a negative control for these experiments, we did not detect expression of p65 (data not shown). Collectively, we suspect that these innate immune molecules may bind to EV-associated lncRNAs to either activate their enzymatic activities or, alternatively, to suppress their functions in recipient cells. Future studies will better focus on the potential significance of this binding.

### Stem Cell EVs Rescue Cellular Viability in HIV-1 Damaged Neurons

Based on the above data showing that NPC-derived neurospheres are susceptible to HIV-1 infection and that stem cell EVs may exert neuroprotective effects, we next assessed the potential functional effects of stem cell EVs on damaged neurons *in vitro*. Here, we utilized two-dimensional cultures of the neuronal cell line SHSY5Y; to induce neuronal damage cells were exposed to EVs derived from U1 (HIV-1 infected) cART-treated cells (“damaging EVs”). Our rationale was based on results from previous studies showing that EVs from HIV-1 infected cells contain viral by-products (e.g. TAR, Tat, Nef, *env*) that can be transferred to recipient cells [107, 108, 109, 110]. Thus, we hypothesized that exposure of neurons to these damaging EVs would be characterized by a loss in cellular viability and, moreover, that stem cell EVs may have the potential to reverse this effect.

For this assay SHSY5Y cells were treated with EVs from either U1 cART-treated cells or a combination of U1 cART EVs with stem cell EVs. Representative images in Fig. 7d show that after approximately eight days in culture the untreated cells appeared healthy, well-attached, and exhibited a spindle-like morphology. Conversely, neurons exposed to U1 cART EVs displayed an altered appearance that was characterized by a reduction in attachment and clumping/ aggregation of the remaining attached cells. However, treatment with either iPSC or MSC EVs at an approximate ratio of 1:1000 (recipient cell: EV) promoted the re-attachment and spreading of cells that more closely resembled the untreated control culture. To ascertain whether these effects may be mediated by EV-associated RNA, stem cell EVs were exposed to UV(C) irradiation to deactivate RNA. Treatment of neurons with irradiated EVs appeared to abrogate the morphological effects associated with non-irradiated stem cell EVs. Cellular viability was next assessed using a luminescence-based assay. As shown in Fig. 7e, treatment of neurons with U1 cART EVs resulted in a slight decrease in viability relative to the control. While the treatment with stem cell EVs did improve cellular viability, only iPSC EVs were associated with a significant increase. Furthermore, there were no significant effects of UV(C) irradiated EVs on cellular viability.

In a separate experiment, we have also utilized EVs isolated from cells infected with the human coronavirus OC43 to induce neuronal damage (Supplementary Fig. 7). Similar to what was observed with U1 cART EVs, neurons exposed to OC43 EVs detached from the substrate and appeared round in morphology. Again, treatment with both iPSC and MSC EVs appeared to permit the re-attachment and spreading of cells. Exposure to OC43 EVs also resulted in a strong reduction of neuronal viability and treatment with both iPSC and MSC EVs significantly abrogated this effect. Moreover, exposure of stem cell EVs to UV(C) irradiation substantially inhibited their ability to restore cellular viability. Collectively, these results suggest that stem cell EVs may at least partially reverse the damaged phenotypes associated with infection. Future studies will focus on optimizing these experiments to better characterize the mechanism(s) associated with virally induced damage and stem cell EV-mediated repair.

## Discussion

Throughout the last decade, iPSCs and their derivatives have revolutionized the fields of regenerative medicine, drug development, and disease modeling [11]. With respect to the CNS, iPSC-derived NPCs and NSCs have been used to successfully generate 3D neuronal cultures, or neurospheres. Relative to traditional 2D cultures, 3D models are expected to serve as a more relevant platform for the study of biological processes. For example, neurospheres have the potential to provide enhanced molecular insight into key cellular processes relating to migration, replication, apoptosis [27]. Additionally, their ability to be cultured in serum-free and chemically defined conditions renders them particularly important for studies pertaining to neural development and neurogenesis [32]. Thereby, neurospheres may represent an ideal platform for preliminary *in vitro* studies surrounding the pathogenesis of various neurological diseases and disorders. Previous works by others have utilized 3D neuronal cultures to successfully model Zika and HSV-1 infections [33, 34, 35, 49]. More recently, Dos Reis et al. reported results demonstrating that HIV-1 infected microglia could be incorporated into human brain organoids [36]. With respect to HIV-1 infection, it is also important to note that a recent study by Rai et al. demonstrated that iPSC-derived microglia displayed significant expression of both CD4 and CCR5 receptors and were readily infected by HIV-1 [111]. Importantly, we also have data demonstrating that our NPC-derived neurospheres, which were generated from human iPSCs, express both CD4 and CCR5. Moreover, this expression remained relatively consistent after treatment of infected samples with cART and EVs; this data further supports our claims regarding the susceptibility of neurospheres to HIV-1 infection (Supplementary Fig. 8). To the best of our knowledge, this is the first study utilizing human NPC-derived neurospheres for retroviral infection.

The use of well-characterized and qualified material, as well as the establishment of robust and reproducible protocols is essential for ensuring the integrity of scientific data. The protocols employed here permitted the efficient formation of neurospheres, resulting in well-defined cultures that displayed uniformity in size and appearance throughout the duration of the study; thereby overcoming some of the challenges that have historically been associated with use of 3D neuronal cultures [52]. The NPCs used for these experiments also underwent extensive characterization to authenticate not only their identity, but also their differentiation potential. Our data suggests that NPC-derived neurospheres maintain their ability to differentiate into both mature neurons (e.g. dopaminergic, GABAergic, glutamatergic) and glial cells (i.e. astrocytes and potentially microglia) in 3D cultures. It is worth noting that properly distinguishing between microglia and macrophages has historically been challenging as these cell types are often associated with many of the same surface markers. Due to the difficulties associated with their phenotypic identification, here we use the term “microglia-like cells” to correlate with positive expression of either CD163, CD11b, CD45, and IBA-1 as these markers are associated with either resting or activated microglia and macrophages [112, 113, 114, 115, 116]. While the potential of neurospheres to differentiate into microglia remains a topic of debate, it is important to consider that the expression of cell-specific markers may greatly vary in response to different biological and biochemical parameters such as cell maturation, cell culture conditions, and the inherent phenotypic heterogeneity within a given cell type [117]. Along these lines, D’Aiuto et al. recently utilized human iPSC-derived NPCs to generate scaffold-free adherent 3D neuronal cultures which expressed not only neuronal (glutamatergic, GABAergic, dopaminergic) markers but also GFAP and IBA-1 [35]. In addition, we suspect that exposure of neurospheres to a stimulating agent (e.g. LPS) could further promote the expression of a microglial-like phenotype.

Despite being regarded as “immune privileged,” the CNS contains at least three types of immune cells, namely CD4+ T cells, macrophages, and microglia, that can be infected by HIV-1 and subsequently serve as viral reservoirs [59]. In the CNS, HIV-1 can activate numerous inflammatory mediators and enzymes that disrupt downstream signaling pathways in both neuronal and glial cells [118]. HIV-1 infection is also associated with a range of neurological symptoms collectively known as HAND, and the release of toxic viral proteins by infected cells is expected to contribute to HAND pathogenesis [5,6,7,8]. However, the underlying mechanisms remain poorly understood; consequently, there is a need to develop advanced assays to better characterize the molecular mechanisms that contribute to HAND and other HIV-1 associated disease phenotypes. In these studies we sought to assess the feasibility of using NPC-derived neurospheres for future studies modeling HIV-1 infection.

Dual-tropic HIV-1 89.6 utilizes the CXCR4 and CCR5 coreceptors; both of which are expressed on resident CNS cells including macrophages, microglia, and astrocytes [118]. Whether or not astrocytes can harbor productive infection remains a matter of debate; although integrated HIV-1 has been detected in astrocytes isolated from infected patients and it has been reported that infected astrocytes can produce viral proteins (e.g. Tat, Rev, Nef) that contribute to neuroinflammation and neuronal damage [2, 59, 62, 119, 120, 121, 122]. Results from our experiments confirmed viral replication in neurospheres as evidenced by the expression of p24 capsid protein, uncleaved Gag polyprotein Pr55, and accessory protein Nef, all of which were detected after fourteen days in culture. As a measure of validation, we also detected p24 in neurospheres exposed to two other lab-adapted strains (JR-CSF and CHO40). IP assays further revealed that the highest copies of HIV-1 TAR DNA were associated with microglia-like cells. The JR-CSF infected sample also displayed relatively high copy numbers of TAR DNA associated with astrocytes; a finding that is not entirely surprising given that this viral isolate originated from the cerebrospinal fluid of an infected patient.

Perhaps more importantly, our data suggests that treatment with IL-34, M-CSF, and TGFβ-1 for a period of one week had the potential to induce a microglia-like phenotype in neurospheres. Notably, we also found that cytokine-treated neurospheres harbored more efficient viral replication. To highlight this point, we observed undetectable expression of Pr55 Gag polyprotein that corresponded with a robust expression of cleaved p24 capsid protein. Moreover, the copy numbers of TAR and *env* RNA were relatively similar in cytokine-treated samples, thus indicating an increased level of HIV-1 transcription in these cultures. When considering the mixed cellular composition of differentiated neurospheres, which clearly retained a distinct proportion of NPCs (Fig 1.d), we speculate that addition of IL-34, M-CSF, and TGFβ-1 induced these progenitor cells to differentiate in a myeloid/microglia phenotype that was more permissible to HIV-1 infection.

Overall, this data is in agreement with HIV-1 tropism towards glial-like cells. However, whether HIV-1 DNA represents a mixture of both non-integrated DNA as well as integrated proviral DNA in neurospheres has not been quantitatively analyzed. Along these lines we have recently performed preliminary integration experiments (Supplementary Fig. 9) adapted from previously published *Alu* PCR protocols [123, 124, 125, 126]. While the data generated from these experiments suggests a mixture of both integrated (> 50%) and unintegrated (< 50%) HIV-1 DNA, it does not specifically address factors relating to site of integration and does not distinguish between cell types infected (i.e. astrocytes vs. microglia). For example, in the cART samples, although integrated copies (Alu-*gag*) are increased (i.e. lower Ct value) relative to unintegrated copies (*Gag*-only), this does not address whether the integrated copies are from astrocytes or microglia or whether unintegrated DNA are preferentially increased in one cell type over another. Therefore, our future experiments will have to carefully address a number of issues including the location of integration in each cell type prior to cART and post-cART, and whether there is a difference in integration depending on the viral strain used (e.g. dual tropic vs. neurotropic). These complex sets of questions can, however, be addressed in “3D” neurosphere models, especially if we are able to maintain long-term cultures over a period of three to six months prior to genome analysis.

Interestingly, we observed unique expression profiles of cell-specific markers among uninfected and infected neurospheres (Fig. 3a). First, these results are in general agreement with our previous assays (ICC and IP) which relied on the use of antibodies associated with astrocytes (GFAP), microglia-like cells (CD11b, CD163), neurons (FOXA2, GAD65), and NSCs (SOX2) as varying levels of expression for each of these markers was detected. GFAP is an intermediate filament protein that is most commonly associated with astrocytes and increased GFAP expression is a hallmark of astrocyte activation (e.g. astrogliosis or astrocytosis) [127, 128]. Our observation of increased GFAP expression in HIV-1 infected neurospheres is consistent with previous literature that has characterized HIV-1 infection by astrocytosis [129, 130, 131]. As indicated previously, CD11b is an integrin protein that can be used to identify either resting or activated microglia-like cells. Our observation of relatively consistent CD11b expression may be related to the specificity of the antibody used. While the apparent increase in the expression levels of neuronal markers (TH^+^, GAD65^+^/67^+^, BNPI^+^) in HIV-1 infected neurospheres were unexpected, further studies are warranted to see if this phenotype reproduces. Without having an accurate quantification of the cellular composition of neurospheres, it cannot be ruled out that some samples had inherently higher levels of certain cell types to begin with. In addition, the spatial location of certain cell types within each sphere needs to be considered. Without knowing the kinetics of HIV-1 infection in this 3D platform, it is also possible that the virus was not able to efficiently penetrate the neurospheres; along these lines, an incubation period longer than fourteen days may be required to better assess the long-term effects of HIV-1 infection on certain cell types. Nonetheless, we believe that these novel results provide a justifiable proof-of-concept for continued studies that can further fine tune these experimental details.

The primary focus of this manuscript surrounds retroviral infections, namely HIV-1 and HTLV-1. HTLV-1 is the etiological agent of both adult T cell leukemia (ATL) and the progressive neurological disorder known as HTLV-1-associated myelopathy/tropical spastic paraparesis (HAM/TSP) [66, 67, 68, 69]. As in the case of HIV-1, the underlying mechanisms contributing to HTLV-1 associated pathologies remain poorly understood. Thus, there is an imminent need for the identification of suitable *in vitro* platforms to drive the field of HTLV-1 research. Exposure to IR is known to initiate cell cycle arrest; moreover, we have recently published data using IR to activate transcription of HTLV-1 in HUT 102 cells [70, 71, 72, 132]. Therefore, we rationalized that co-culturing of irradiated HUT 102 cells with neurospheres would provide a reliable platform to assay for infection. To this end, we performed western blot to assess the relative expression of Tax and p19, two indicators of HTLV-1 viral replication. When accounting for background levels in the non-irradiated HUT 102 control sample, there were clear increases in the expression of both Tax and p19. RT-qPCR also confirmed increased expression of *tax, env*, and *hbz* in the co-cultures. Treatment with IFN/AZT was also associated with reductions in the corresponding protein/gene expression levels. Therefore, this preliminary data suggests that neurospheres may be reliable models for studying the mechanisms surrounding HTLV-1 associated neuropathogenesis. Future experiments will utilize other HTLV-1 and HIV-1 infected cells (e.g. U1 cells) that have been irradiated to potentially increase viral spread.

In recent years our laboratory has also focused extensively on EVs and considerable efforts have been made in the characterization of EVs from both infected and non-infected cells [72, 76, 108, 109, 110, 133, 134, 135]. Here specifically, we were interested in examining the functional effects of stem cell EVs on HIV-1 infected neurospheres. Our rationale was based on the results from numerous studies which have highlighted the protective and reparative properties of stem cell EVs among various CNS pathologies, as reviewed elsewhere [43, 48, 48, 136]. For these studies we utilized well-characterized preparations of iPSC and MSC EVs that have previously demonstrated reparative effects in various *in vitro* assays (e.g. cellular migration, cellular viability, angiogenesis) [76].

The results from our uptake assays first confirmed that EVs were capable of penetrating neurospheres. Moreover, fluorescently labeled EVs remained detectable for at least eight days post-treatment. These results were consistent among two independent experiments that used different fluorescent probes to stain either EV lipid membranes or EV-associated RNA. Interestingly, gelatin zymography indicated that iPSC and A549 EVs had relatively high levels of the active forms of the gelatinases MMP9 and MMP2 as compared to MSC EVs. While we observed distinct differences in the morphological appearance of iPSC EV and A549 EV-treated neurospheres, it is imprudent to attribute this effect to any specific EV cargo at this time. More importantly, these results demonstrated feasibility and provided the justification for subsequent experiments focused on the functional effects of stem cell EVs in neurosphere cultures.

As previously mentioned, over amplification of inflammatory pathways, coupled with the release of toxic viral proteins from infected cells, is believed to contribute to the neuronal damage associated with HIV-1 [5,6,7,8]. Additionally, activated microglia release neurotoxic factors including cytokines and chemokines (e.g. IL6, IL8, IL1β, TNFα, IFNα) that can further exacerbate the inflammatory response [61, 118]. In recent years, the neuroprotective and anti-inflammatory properties of stem cell EVs have been well-documented; in the context of the CNS, there is accumulating evidence that stem cell EVs may be able to reverse neuronal apoptosis and modulate the polarization of both astrocytes and microglia from pro-inflammatory to anti-inflammatory phenotypes [85, 86, 87, 88, 89, 90, 91, 92, 93, 94, 95, 96]. To the best of our knowledge, the functional effects of stem cell EVs on 3D cultures has not previously been reported.

As expected, we observed increased expression of pro-inflammatory cytokines TNFα, IL-8, and IL-1β in HIV-1 infected neurospheres. Additionally, we also observed increased expression of both cleaved PARP-1, Caspase-3, and BAD in response to HIV-1 infection. These findings are consistent with the activation of cellular stress response pathways and the initiation of apoptotic signaling cascades [137, 138, 139]. The addition of stem cell EVs to HIV-1, cART-treated neurospheres abrogated each of these effects; thereby pointing towards a potential EV-mediated reversal of inflammatory and apoptotic signaling in these cultures. Since Caspase 3 is classically regarded as an “executioner” caspase [140, 141] this data also suggests that damaged cells may have been in the early stages of apoptosis, thus making them more receptive to reparative intervention by EVs. While our data suggests that A549 (cancer) EVs may also exert anti-apoptotic and anti-inflammatory properties, further interpretation and analysis of these results is outside the scope of the current manuscript. Unlike stem cell EVs, the use of cancer EVs for potential repair is not well-studied and the long-term functional outcomes associated with exposure to cancer EVs needs to be better characterized.

Although the results from these experiments aren’t cell-type specific, our future experiments will more thoroughly examine which cell types in neurospheres are associated with increased levels of apoptosis. Additionally, our future experiments will focus on examining the expression levels of other critical mediators of apoptosis, including p53 and its kinases (e.g. ATM, ATR), as well as a broader range of proteins involved in the early, middle, and late stages of apoptosis to better define the reparative mechanisms of stem cell EVs.

EVs are associated with a rich diversity of cargo which includes various non-coding RNAs (i.e. small non-coding and long non-coding RNAs), as described elsewhere [142, 143, 144, 145, 146, 147]. Mechanistically speaking, this cargo is expected to drive the functional effects that are attributed to EVs. While many studies have evaluated the reparative potential of various EV-associated miRNAs, the characterization of EV-associated lncRNAs remains largely unexplored. The data from our lncRNA analysis highlighted distinct differences among A549 (cancer) EVs and stem cell EVs, particularly with respect to chromosome localization and patterns of differentially expressed transcripts. Although lncRNAs from both iPSC and MSC EVs mapped to some chromosomes as compared to A549 EVs, it is difficult to decipher the functional significance of this data in the absence of mutant reagents. For instance, iPSC and MSC EVs contain lncRNAs that map to certain chromosomes; however, it is not clear if the same EVs that contain these RNAs also contain cytokines and reparative proteins. Furthermore, the actual copy number of these RNAs in various subpopulations of EVs (e.g. those ranging from ∼30 to 220 nm in size) requires further purification, followed by functional repair assays and then proteomics and RNA seq. This will better define which EV subtype contains cytokines, proteins, and RNAs that map to a certain chromosome. Therefore, future experiments will better define the type of EVs needed to repair, as well as their protein and RNA cargo that may functionally regulate either the host genome, splicing in the nucleus, translation inhibition, or mRNA degradation in the cytoplasm.

The identification of at least two iPSC EV-associated lncRNAs (i.e. AC120498.9, ADIRF-AS1) whose secondary structures contained sequences of double-stranded RNA ≥ 30 bp in length led us to examine their interactions with different RNA binding proteins *in vitro*, as this property has previously been linked to the activation of PKR [105, 106]. In an attempt to decipher their function, our data indicates that these EV-associated lncRNAs may regulate PKR, RIG-I, Dicer, and potentially eIF2α phosphorylation. However, it is not clear at this point whether the binding of innate immune molecules to these RNAs results in their increased stability or change of enzymatic function. Additionally, when mapping the secondary structures of iPSC EV-associated lncRNAs, we noted that many of these each contained multiple circular loops that resemble the structure of circular RNAs (circRNAs). For reference, we have included a panel of images of these structures in Supplementary Fig. 10. While circRNAs represent a relatively novel class of non-coding RNAs, there is emerging evidence to suggest that they may predominantly function as miRNA sponges to prevent binding of miRNAs to their target genes [148, 149].

Therefore, we speculate that these structures could potentially serve as RNA sponges for cellular miRNA that regulate pathways such as apoptosis, cell cycle, and cytokine secretion. Future experiments will better define how these stem and loop RNA structures are capable of binding to dsRNA proteins and how they may also regulate miRNAs that may be essential for cellular repair in damaged cells.

It has been shown that EVs released from HIV-1 infected cells contain viral by products including Nef, Tat *env*, and TAR [107, 108, 110, 150]. The effects of these damaging EVs include rendering recipient cells more susceptible to infection, enhancing the spread of infection, and induction of cellular death, as recently reviewed [151]. Since we were unable to directly study their effects in HIV-1 infected neurospheres, we utilized 2D cultures of neurons to determine if exposure to EVs from HIV-1 infected cells could induce neuronal death and, subsequently, to assess whether stem cell EVs could exert any functional effects on damaged cells. Our data confirmed a decline in cell healthy and slight reduction in viability upon treatment with damaging EVs, while stem cell EVs appeared to reverse these processes. Even though UV(C) treatment did not have a significant impact on cellular viability, the observed morphological differences in neurons treated with irradiated stem cell EVs relative to non-irradiated EVs suggests that UV(C) may have partially impaired the functional effects of both iPSC and MSC EVs. On the other hand, performance of the same assay using OC43 EVs to induce neuronal damage produced considerable outcomes. Here, treatment of damaged cells with stem cell EVs yielded a significant increase in viability whereas UV(C) irradiation abrogated their effects. Thus, these experiments highlight the existence of different mechanisms surrounding both EV-induced damage and EV-mediated repair in recipient cells.

Overall, the results from this study have demonstrated that NPC-derived neurospheres are susceptible to retroviral infection and, therefore, may serve as an alternative platform for studying the molecular mechanisms associated with HIV-1. Moreover, we also demonstrated the functional relevance of stem cell EVs in the context of mediating cellular repair induced by viral damage. Importantly, this data is in consensus with numerous other studies which have described the neuroprotective and anti-inflammatory properties of stem cell EVs. We acknowledge that further experiments are necessary to more carefully define and characterize the mechanisms contributing to both viral-induced damage and EV-mediated repair. However, based upon the current data, we hypothesize that the functional effects of stem cell EVs in both uninfected and virally infected cells may be partially mediated through the interactions between EV-associated lncRNAs and nucleic acid sensors (e.g. PKR, ADAR1, RIG-I) in recipient cells. For example, in uninfected cells this binding could trigger the activation of intracellular signal transduction pathways, thereby resulting in increased production of interferon and a potent antiviral response that may ultimately protect the cell from viral infection. Conversely, infected cells would have already mounted an immune response to undergo cell cycle arrest. In this scenario, EV-associated lncRNAs could prolong this effect by binding to, and thus inactivating, the host innate immune molecules to prevent further viral replication. In turn, this would also dampen the production of damaging EVs from infected cells. Over time, the EV-associated cytokines and growth factors will promote cellular replication of both infected and uninfected cells. However, we believe that the net outcome will confer an overall protective effect, since we suspect that uninfected cells represent a majority (≥ 90%) of the total cell population. In support of this, we have data showing that treatment of HIV-1 infected neurospheres with stem cell EVs results in increased expression of different cell populations including NPCs, astrocytes, and microglia-like cells (data not shown). Thus, we believe that this research sets an important precedent for the continued use of NPC-derived neurospheres for modeling biological processes relating to the CNS. We look forward to utilizing these cultures to further expand our knowledge in this area.

## Methods

### Cell Culture

#### NPCs and Neurospheres

Normal Human Neural Progenitor Cells (ATCC ACS-5003) were expanded per the manufacturer’s recommendations. Cells were cultured with STEMdiff Neural Progenitor Medium (STEMCELL technologies) in cell matrix-coated flasks. Media was changed every other day. Cells were subcultured using Accutase diluted 1:1 with PBS. To generate neurospheres, NPCs were seeded at approximately 1.0 x 10^5^ cells per well in ultra-low attachment, U-shaped bottom 96-well plates in Neural Progenitor Medium. After approximately 48 to 72 hours, the medium was removed from the wells and replaced with DMEM:F12 (ATCC 30-2006) + Dopaminergic Neuron Differentiation Kit (ATCC ACS-3004) to induce differentiation. Neurospheres received a complete media replacement every two to three days for a period of two weeks. Phase contrast images were acquired using an inverted microscope and calibrated measurements were performed using Zen imaging software (Zeiss).

To further induce differentiation in neurospheres, the fully supplemented differentiation medium (listed above) was spike with the following cytokines: IL-34 (100 ng/mL), M-CSF (25 ng/mL), and TGFβ-1 (50 ng/mL). Neurospheres were cultured in this medium for an additional seven days, receiving fresh treatments every two to three days, prior to HIV-1 infection.

#### Cell Lines

The human neuroblastoma cell line SHSY5Y (CRL-2266), the human astrocytoma cell line CCF-STTG1 (CRL-1718), human monocytic leukemia cell line THP-1 (TIB-202), and the human lung carcinoma cell line A549 (CCL-185) were obtained from the American Type Culture Collection (ATCC). SHSY5Y cells were cultured in a 1:1 mixture of EMEM (ATCC 30–2003) and F-12K (ATCC 30–2004) supplemented with 10% fetal bovine serum (FBS) (ATCC 30–2020). CCF-STTG1 and THP-1 cells were cultured in RPMI-1640 (ATCC 30–2001) supplemented with 10% FBS. A549 cells were cultured in F-12K (ATCC 30–2004) supplemented with 10% FBS. Cells were maintained in culture following the manufacturer’s recommendations. HIV-1 infected U937 Cells (U1; NIH AIDS Reagent Program) were cultured in RPMI-1640 supplemented with 10% exosome-depleted FBS.

To generate MDMs, THP-1 cells and U1 cells were treated with phorbol 12-myristate 13-acetate (PMA; 100 nM) and incubated for four to five days. U1-MDMs then received two treatments of cART (emtricitabine, tenofovir disoproxil fumarate, ritonavir, darunavir; 2.5 µM each) over a five day incubation period.

#### Immunocytochemistry (ICC)

Cells were fixed in 4% paraformaldehyde. Neurospheres were embedded in OCT and sectioned at a thickness of 8 µm. Samples were mounted on charged slides. After permeabilization (0.2% Triton X-100+ 0.01% Tween 20) and blocking (5% goat serum), cells were incubated overnight with the following primary antibodies: α-SOX2 (Santa Cruz Biotechnology sc-365823; 1:50), α-Tyrosine Hydroxylase (TH; Millipore AB152; 1:1000), α-GFAP (Stemcell Technologies 60048; 1:100), α-IBA-1 (Santa Cruz Biotechnology sc-32725; 1:50), α-Tubulin β 3 (Tuj1; Biolegend MMS-435P; 1:200). The appropriate secondary antibody (Alexa Fluor 594 goat anti-rabbit, Alexa Fluor 594 goat anti-mouse, Alexa Fluor 488 goat anti-rabbit, or Alexa Fluor 488 goat anti-mouse) was then added for one hour. Nuclei were counter-stained with DAPI. As an additional quality control measure, H&E staining was also performed on sectioned neurospheres. Fluorescent and brightfield imaging was performed using the BioTek Cytation5 imager fitted with the appropriate filters.

### Infection of Neurospheres

#### HIV-1 Infection

Differentiated neurospheres were infected with either dual-tropic HIV-1 89.6 (MOI:10; ∼250 ng of p24), JR-CSF (MOI:10), or CHO40 (MOI:10) and InfectinTM (Virongy, LLC) (day 0). The total final volume of spheres + virus + Infectin was approximately 200 µL. After 48 hours of incubation (day 2), neurospheres were gently washed with PBS and received fresh differentiation media with or without cART cocktail (lamivudine (NRTI), tenofovir disoproxil fumarate (NRTI), emtricitabine (NRTI), indinavir (protease inhibitor); 2.5 µM each). HIV-1 + cART neurospheres also received treatment with stem cell EVs at an approximate ratio of 1:250 (uninfected recipient cell to EV). HIV-1 89.6 neurospheres received additional cART and EV treatments every two to three days. After a period of seven to fourteen day, neurospheres and supernatants were harvested for downstream analysis. To enrich cytokines from neurosphere supernatants, Nanotrap particles (NT082/NT080; Ceres Nanosciences, Inc.) were used following protocols recently published by our lab [72, 133, 152].

#### HTLV-1 co-culture

HUT 102 cells (log phase) were cultured in RPMI-1640 supplemented with 10% heat-inactivated, exosome-depleted FBS for 5 days. Cells were then seeded at 1.0 x 10^6^ viable cells/mL and exposed to IR using a RS-2000 X-Ray Irradiator (Rad Source Technologies) at a dose of 10 Gy. Irradiated cells were co-cultured with differentiated neurospheres (day 0) at an approximate ratio of 1:100 (infected cells: uninfected cells in neurospheres) with or without combination treatment of interferon-α (10 K unit) and zidovudine (20 µM), hereafter referred to as IFN/AZT. Treatments were added every two to three days and after fourteen days the neurosphere co-cultures were harvested for downstream analysis.

#### Western Blot

Protein lysates were analyzed via Bradford assay following the manufacturer’s recommendations (Bio-Rad). For western blot, equal amounts of protein were mixed with Laemmli buffer, heated at 95°C, and separated on a 4-20% Tris-glycine SDS gel (Invitrogen). Proteins were transferred to an Immobilon PVDF membrane (Millipore) overnight at 50 milliamps. After blocking the membrane with 5% milk in PBS with 0.1% Tween-20, the appropriate primary antibody was added for overnight incubation at 4 °C. Antibodies used for these experiments included α-p24 (NIH AIDS Reagent Program; 1:1000), α-Nef (NIH AIDS Reagent Program; 1:1000), α-Tax (kindly provided by Dr. Scott Gitlin, University of Michigan; 1:1000), α-p19 (Santa Cruz Biotechnology sc-1665; 1:200), α-Integrin αM/CD11b (Santa Cruz Biotechnology sc-515923; 1:200), α-CD163 (Santa Cruz Biotechnology sc-20066; 1:200), α-glutamine synthetase (Santa Cruz Biotechnology sc-74430; 1:200), α-GFAP (Santa Cruz Biotechnology sc-166481; 1:200), α-Tyrosine Hydroxylase (Millipore Sigma AB152; 1:200), α-FOXA2 (Santa Cruz Biotechnology sc-101060; 1:200), α-GAD65 (Santa Cruz Biotechnology sc-377145; 1:200), α-GAD67 (Santa Cruz Biotechnology sc-28376; 1:200), α-BNPI (Santa Cruz Biotechnology sc-377425; 1:200), α-SOX2 (Santa Cruz Biotechnology sc-365823; 1:200), α-PARP-1 (Santa Cruz Biotechnology sc-7150; 1:200), α-Caspase 3 (Santa Cruz Biotechnology sc-7148; 1:200), α-BAD (Santa Cruz Biotechnology sc-8044; 1:200), microglia marker sampler panel (α-CD11b, α-CD45, α-IBA-1; Abcam ab226482; 1:1000 each), α-IBA-1 (Santa Cruz Biotechnology sc-32725; 1:200), α-TNFα (Santa Cruz Biotechnology sc-52749; 1:200), α-IL-8 (Santa Cruz Biotechnology sc-376750; 1:200), α-IL-1β (Santa Cruz Biotechnology sc-32294; 1:200), α-PKR (Santa Cruz Biotechnology sc-136352; 1:200), α-DICER (Cell Signaling Technology 3363S; 1:1000), α-ADAR1 (Santa Cruz Biotechnology sc-271854; 1:200), α-RIG-I (Santa Cruz Biotechnology sc-376845; 1:200), α-MMP9 (Cell Signaling Technology 2270S; 1:200), α-CD4 (R&D Systems MAB3791; 1:1000), α-CCR5 (R&D Systems MAB1802; 1:000) and α-Actin (Abcam ab-49900; 1:5000). Membranes were subsequently washed, incubated with the appropriate secondary antibody at 4 °C, and then immediately developed using either Clarity or Clarity Max Western ECL Substrate (Bio-Rad). Chemiluminescence imaging was performed with the ChemiDoc Touch Imaging System (Bio-Rad). For reference, all full-length blots are presented in Supplementary Fig. 11.

#### Reverse Transcription and Quantitative PCR

Total RNA from neurospheres and EVs was extracted using TRIzol reagent (Invitrogen) following the manufacturer’s recommendations. Reverse transcription was carried out with the GoScript Reverse Transcription System (Promega) following the manufacturer’s recommended protocols. The following specific primers were used: HIV-1 trans-activation response (TAR) reverse primer (5′-CAACAGACGGGCACACACTAC-3′), HIV-1 envelope reverse primer: (5′-TGGGATAAGGGTCTGAAACG-3′), HTLV-1 Tax reverse primer (5′-AACACGTAGACTGGGTATCC-3′), HTLV-1 bZIP factor (HBZ) reverse primer (5′-TGACACAGGCAAGCA TCG-3′), HTLV-1 envelope reverse primer (5′-CCATCGTTAGCGCTT CCAGCCCC-3′) [68, 129], AC120498.9 reverse primer (5′-TGAAGTCTCGTTCTGTTGCC-3′), ADIRF-AS1 reverse primer (5′-CGGAGTCTTACTCTGTC ATCCA-3′), and GAPDH reverse primer (5′-CAGAGTTAAAAGCAGCCCTGGT-3′).

qPCR for HIV-1 TAR and envelope was carried out with IQ Supermix (Bio-Rad). The following primers/probes were used: TAR forward primer (5′-GGTCTCTCTGGTTAGACCAGATCTG-3′), TAR reverse primer (5′-CAACAGACGGGCACACACTAC-3′), TAR Probe ( 5′-56-FAM-AGCCTCAAT AAAGCTTGCCTTG AGTGCTTC-36-TAMSp-3′). Serially diluted DNA from HIV-1 8E5 cells (NIH AIDS Reagent Program) were included as standards [129]. The reaction conditions were 50°C for 2 minutes (1 cycle), 95 °C for 3 minutes (1 cycle), 95 °C for 15 seconds and 60°C for 40 seconds (41 cycles). Quantification was determined from the average cycle threshold (Ct) values of each sample relative to the standard curve. All reactions were run triplicate on the CFX96 Real-Time PCR Detection System (Bio-Rad).

qPCR of HTLV-1 Tax, HBZ, and envelope was carried out with SYBR Green Master Mix (Bio-Rad) using the following primers: Tax forward primer (5′-ATCCCGTGGAGACTCCTCAA-3′), Tax reverse primer (5′-AACACGTAGACTGGGTATCC-3′), HBZ forward primer (5′-AGAACGCGA CTCAACCGG-3′), HBZ reverse primer (5′-TGACACAGGCAAGCATCG-3′, Tm), envelope forward primer (5′-CGGGATCCTAGCGTGGGAA CAGGT-3′), envelope reverse primer (5′-CCATCGTTAGCGCTTCCAGCCCC-3′). Serially diluted DNA from HTLV-1 infected cells (HUT 102) served as standards [68]. The reaction conditions were as follows: Tax: 50°C for 2 minutes (1 cycle), 95°C for 2 minutes (1 cycle), 95°C for 15 seconds, 51°C for 20 seconds, and 72°C for 40 seconds (40 cycles), 65°C for 5 seconds (1 cycle), 95°C for 5 seconds (1 cycle); HBZ: 50°C for 2 minutes (1 cycle), 95°C for 2 minutes (1 cycle), 95°C for 15 seconds, 65.7°C for 20 seconds, and 72°C for 40 seconds (40 cycles), 65°C for 5 seconds (1 cycle) and 95°C for 5 seconds (1 cycle); envelope: 50°C for 2 minutes (1 cycle), 95°C for 2.5 minutes (1 cycle), 95°C for 15 seconds and 64°C for 40 seconds (40 cycles), 65°C for 5 seconds (1 cycle), and 95°C for 5 seconds (1 cycle) [68]. Sample quantification was determined from the average cycle threshold (Ct) values of each sample relative to the standard curve. All reactions were run in triplicate (technical replicates) on the CFX96 Real-Time PCR Detection System (Bio-Rad).

qPCR of EV-associated lncRNAs was carried out with SYBR Green Master Mix (Bio-Rad) using the following primers: AC120498.9 forward primer (5′-GGTCCTGGGAATCTGCTTT-3′), AC120498.9 reverse primer (5′-GCCTAGCCAACATGGTGAAA-3′), ADIRF-AS1 forward primer (5′-TGCAAT GGCGGGATCTT-3′) and ADIRF-AS1 reverse primer (5′-GTGGCTCATGCCTGTAATCT-3′). qPCR of EV-associated GAPDH was carried out with IQ Supermix (Bio-Rad) using the following primers and probe: GAPDH forward primer (5′-GAAGGTGA AGGTCGGAGTCAA C-3′), GAPDH reverse primer (5′-CAGAGTTAAAAGCAGCCCTGGT-3′), GAPDH probe (5′-/56-FAM/TTTGGT CGTATTGGGCGCCT/36-TAMSp/-3’). The reaction conditions were as follows: 95°C for 10 minutes (1 cycle), followed by 41 cycles of 95°C for 15 seconds, 54.8 °C/54.7°C (depending on primers) for 20 seconds, and 72°C for 10 seconds.

All reactions were run in triplicate on the CFX96 Real-Time PCR Detection System (Bio-Rad). Data were normalized to GAPDH and the relative expression levels were evaluated using the 2^−ΔΔCT^ method [153].

#### HIV-1 Integration Assay

DNA was extracted from neurosphere samples and 8E5 (positive control) cells using the Qiagen DNeasy Blood & Tissue Kit following the manufacturer’s protocol. DNA was quantified with the NanoDrop and 100 ng of DNA from each sample was used for the Alu-*Gag* PCR assay. The following primers/probe were used: 1^st^ integration (targeting Alu-*Gag*): Alu forward primer (5′-GCCTCCCAAAGTGCTGGGATTACAG-3′), HIV-*Gag* reverse primer (5′-GTTCCTGCTATGTCACTTCC-3′) [125]; 2^nd^ integration (targeting HIV-1 TAR): TAR forward primer (5′-GGTCTCTCTGGTTAGACCAGATCTG-3′), TAR reverse primer (5′-CAACAGACGGG CACACACTAC-3′), TAR probe (5′-56-FAM-AGCCTCAAT AAAGCTTGCCTTGAGTGCTTC-36-TAMSp-3′) [129]. The reaction conditions for the 1st integration PCR were as follows: 95°C for 2 minutes (1 cycle), followed by 39 cycles of 95°C for 15 seconds, 50°C for 15 seconds, and 72°C for 3.5 minutes. The reaction conditions for the 2nd integration (quantitative) PCR were as follows: 95°C for 3 minutes (1 cycle), followed by 41 cycles of 95°C for 15 seconds and 60°C for 40 seconds. Samples were run in triplicate on the CFX96 Real-Time PCR Detection System (Bio-Rad) and Ct values were exported for data analysis.

### Isolation and Characterization of EVs

#### Stem Cell EVs and A549 EVs

EV isolation was performed as previously described [72]. Briefly, human bone marrow MSCs (BM-MSCs) (ATCC PCS-500-012) and human lung carcinoma cell line A549 (ATCC CCL-185) were cultured in the presence of exosome-depleted FBS (Thermo Fisher Scientific) prior to collection of conditioned medium. Human iPSCs (ATCC ACS-1019), which were cultured in serum-free/xeno-free medium, received full media changes every day and the conditioned medium was collected and stored at 4°C throughout expansion. Bulk conditioned medium was first spun at 2,000 x g and then filtered with a 0.22 µm filter to remove cellular debris and larger vesicles. Filtered medium then underwent concentration and buffer exchange via TFF (Spectrum) to isolate EVs. Samples were subsequently aliquoted and frozen at -20°C for downstream analysis. The size distribution and concentration of each EV sample was assessed via nanoparticle tracking analysis (NTA) using the NanoSight NS300 (Malvern Panalytical). Prior to analysis the equipment was calibrated per the manufacturer’s protocols. EV samples were diluted in sterile PBS immediately before measurement. The acquired data was analyzed by the instrument’s built-in software.

#### HIV-1 and Human Coronavirus OC43 EVs

U1 cART EVs were isolated via ultracentrifugation. First, supernatants were centrifuged at 1,200 rpm for five minutes to pellet any cellular debris. The resulting supernatant was then centrifuged at 100,000 x g for 90 minutes using the 70Ti rotor (Beckman). The pellet was washed with PBS followed by another centrifugation at 100,000 × g for 90 minutes. EV pellets were resuspended in PBS and then frozen at -20°C for downstream analysis.

A549 cells were infected with 5.0 x 10^5^ TCID50 of human coronavirus OC43 for 96 hours. Supernatants were collected and centrifuged at 2,000 rpm for five minutes, filtered through a 0.22 µm sterile filter, and then centrifuged at 100,000 x g for 90 minutes using the 70Ti rotor (Beckman). The pellet was resuspended in sterile PBS and treated with 45 mJ/cm^3^ of UV(C) LED irradiation. Samples of UV(C)-treated EVs (1, 5, 10 µL) were added to Vero cells for two passages and no viral outgrowth was detected (data not shown). Remaining EVs were frozen at -20°C for downstream analysis.

U1 cART and OC43 EVs were analyzed via NTA using the ZetaView Z-NTA (Particle Metrix). Prior to analysis the equipment was calibrated per the manufacturer’s recommendation and each EV sample was diluted in deionized water prior to analysis. The acquired data was analyzed by the instrument’s built-in software.

#### Transmission Electron Microscopy (negative staining method)

To prepare EV samples for electron microscopy, different dilutions of each sample were prepared and 5 µL was adsorbed to the formvar side of a glow-discharged, 200 mesh copper grid with formvar and carbon coating. Excess sample was removed from the grids with filter paper. Samples were fixed with 4% glutaraldehyde in 0.12 M sodium cacodylate buffer (pH 7.2) for five minutes, washed four times with distilled, deionized water, and stained with 1% aqueous uranyl acetate for 60 seconds. The uranyl acetate was removed with filter paper and allowed to air dry. Imaging was performed using an FEI Talos F200X transmission electron microscope at 80KV (Thermo Fisher Scientific).

#### EV Uptake

EVs were labeled with either SYTO RNASelect Green Fluorescent Stain (Thermo Fisher) or BODIPY 493/503 (Thermo Fisher) following the manufacturer’s recommended protocol. Excess dye was removed via centrifugation using a Sephadex G-10 (Sigma-Aldrich) column with a glass wool bottom packed with 0.75 mL beads. Labeled EVs were added to differentiated neurospheres and incubated at 37°C (day 0) at an approximate ratio of 1:250 (uninfected recipient cell to EV ratio). On day 1, neurospheres received a complete media replacement. Neurospheres also received fresh media on days four and seven. Fluorescent imaging was performed using the BioTek Cytation 5 and phase contrast images were taken with a Zeiss Axio Observer.Z1 microscope.

#### RNA Sequencing and Analysis

Total RNA was extracted from EVs using TRIzol reagent (Invitrogen) following the manufacturer’s recommendations. RNA sequencing and sequence analysis was performed by LC Sciences, LLC. Briefly, the 2100 Bioanalyzer System (Agilent) was used to analyze total RNA and ribosomal RNA was removed per the Ribo-Zero Gold Kit (Illumina). The remaining RNA fragments were used to construct a strand-specific cDNA library using the dUTP method (average library insert size was 300 ± 50 bp) and pair-end sequencing was performed using the Hiseq 4000 (Illumina) following the manufacturer’s recommended protocol. Reads containing adaptor contamination and low quality bases were removed with Cutadapt and in-house perl scripts [154]. FastQC was used to verify transcript quality, Tophat2 [155] and Bowtie 2 [156] were used for mapping reads to the genome, and in-house Perl scripts were used for GO and KEGG enrichment analysis. StringTie [157] was used for transcript assembly and Ballgown [158] was used to perform differential expression analysis. Significant terms for GO analysis were determined by a hypergeometric equation and GO terms with a p value <0.05 were considered significant [159]. All data generated from the RNAseq analysis represents data from one biological replicate each of A549, MSC, and iPSC EVs.

The secondary structures of selected long non-coding RNAs were predicted using the RNAfold Server of the ViennaRNA Websuite [160]. RNA sequences were first retrieved from NCBI and the FASTA sequence was submitted to the server. This server predicted the minimum free energy (mfe) secondary structure of single sequences using a programming algorithm originally developed by Zuker and Stiegler [161]. The secondary structures were predicted using a temperature setting of 37 °C and keeping all other parameters to default [162].

#### Immunoprecipitation for direct PCR

Neurospheres were washed with PBS and treated with 0.25% Trypsin/EDTA (diluted 1:1 with PBS) for three to five minutes at 37°C. Trypsin was neutralized by the addition of RPMI-1640 with 10% FBS. The appropriate antibodies (α-CD11b (Santa Cruz Biotechnology sc-515923; 1:200); α-GFAP (Santa Cruz Biotechnology sc-166481; 1:200), αGAD-65 (Santa Cruz Biotechnology sc-377145; 1:200), α-SOX2 (Santa Cruz Biotechnology sc-365823; 1:200)) were added (5 µg per 10E5 cells) and incubated at 4°C for four hours on a rotator. Protein A+G washed beads (30 µL; 30% slurry in PBS with 0.01% BSA) were added to the cell/antibody mixture and incubated for 1.5 hours at 4°C. Samples were spun at 1,000g for five minutes, washed once with PBS/BSA buffer, and pellets (protein A/G beads + cells + antibody in ∼25µl packed beads) were resuspended in PBS (20 µL). Approximately 1/5 of the solution was used for direct PCR. The first cycle of PCR (heat for strand separation) is sufficient to crack open viruses and eukaryotic cells. An initial titration of an astrocyte cell line with antibody and beads was used to titrate the amount of beads/cell/antibody needed for direct PCR using GAPDH primers (data not shown).

#### RNA Pull-down

The following RNA sequences were synthesized with 5’ Biotin conjugates: UAGCUGGGAUUACAG GCGUGCACCACCACACCCGGCUAAUUUUUGUAUUUUU (AC120498.9) and CUCACUGCAACCUCU GCCUUCCAGGUUCAA (ADIRF-AS1). Whole cell lysate (SHSY5Y, CCF-STTG-1, or MDM) (approximately 75 µg) was incubated with approximately 10 µg of each biotinylated RNA overnight at 4°C. The next day, Streptavidin-Sepharose beads (BioVision) were added and again incubated at 4°C for an additional two hours. The samples were spun at 14000 x g and resuspended in TNE50 + 0.1% NP40. Samples were mixed with Laemmli buffer for subsequent SDS/PAGE and western blot was performed for PKR, DICER, ADAR1, and RIG-I (as described above).

#### Cell Viability Assay

SHYSY5Y cells were seeded in a 96-well plate (day 0). On day one, cells were treated with either U1 cART EVs (1:50,000 approximate recipient cell to EV ratio) or OC43 EVs (1:6,500 approximate recipient cell to EV ratio) to induce damage. Stem cell EVs were treated with 45 mJ/cm3 of UV(C) LED irradiation to inactivate EV-associated RNA. Stem cell EVs were subsequently added at an approximate ratio of 1:1000 (recipient cell to EV ratio). Untreated cells received PBS. Each sample was performed in triplicate. Cells were observed daily. On day eight representative photographs were taken using an EVOS XL microscope and viability was assayed using the CellTiter-Glo Luminescent Cell Viability kit (Promega) per the manufacturer’s protocol. Luminescence was measured with the GloMax Multi Detection System (Promega).

#### Zymography

EVs were added to a 30% slurry of NT80/82 Nanotrap particles (Ceres Biosciences) and PBS was added to bring the final volume to 500 µL. Samples were rotated at 4°C for two hours, spun, and washed with PBS. Laemmli buffer was added to each sample, heated for ten minutes at 37°C, and then spun to pellet the Nanotraps. The resulting supernatants were loaded on a Novex 10% Zymogram Plus (Gelatin) Protein Gel (ThermoFisher Scientific). The gel was run at 120V for approximately 90 minutes, renatured for 30 minutes at room temperature on a rocker, and incubated at 37°C in developing buffer for a period of 24 to 36 hours. After development, the gel was gently stained for an additional 24 to 48 hours with Coomassie blue R-250. Imaging was performed with the ChemiDoc Touch Imaging System (Bio-Rad).

#### Statistical Analysis

Analysis of quantitative data was performed by using Graph Pad Prism. The *P* values were determined via analysis of variance (ANOVA) using multiple comparisons. To justify the use of ANOVA, experimental data was assumed to approximate a normal distribution. *P* values were defined as follows: of greatest statistical significance (< 0.0001), of greater significance (< 0.005), and significant (< 0.05).

## Acknowledgements

We would like to thank all members of the Kashanchi lab for their contributions, especially Gwen Cox and interns Gabrielle Heller and Sahan Raghavan. We also would like express thanks to ATCC management, especially Drs. Mindy Goldsborough and James Kramer for supporting this work. This work was further supported by National Institutes of Health (NIH) Grants AI078859, AI074410, AI127351 -01, AI043894, and NS099029 (to FK), R21DA050176 and R01NS099029 (to FK and LAL), and R33 CA206937 and R01AR068436 (to LAL). The content is solely the responsibility of the authors and does not necessarily represent the official views of the National Institutes of Health. Targeted Pharmaceuticals, LLC partially funded experiments related to 3D cultures and HIV-1 infections.

## Author Contributions

H.B. wrote and edited the manuscript. D.Y. and S.J. contributed to development, characterization, and production of iPSCs and NPCs. H.B. contributed to the production of EVs and C.A.B. contributed to EV analysis. H.B., P.K., S.A.S., M.C., Y.K., J.E. carried out experiments and contributed to data analysis. L.A.L. and N.E. contributed to the experimental designs. F.K. contributed to the overall direction and coordination of the study as well as contributions to experimental design and data analysis.

## Competing Interests

H.B, D.Y., and S.J. are employed by ATCC and L.A.L. is affiliated with Ceres Nanosciences, Inc. All other authors declare no potential conflicts of interest.

