## Supplementary Material for "Retroviral Infection of Human Neurospheres and Use of Stem Cell EVs to Repair Cellular Damage"

**Supplementary Table 1.** Valid data and associated quality scores corresponding to lncRNA from each EV preparation.

| **Sample** | **Raw Data** | | **Valid Data** | | **Valid Ratio(reads)** | **Q20%** | **Q30%** | **GC content%** |
|---|---|---|---|---|---|---|---|---|
| | **Read** | **Base** | **Read** | **Base** | | | | |
| **A549** | 3E+07 | 5.14G | 3E+07 | 3.90G | 75.97 | 96.93 | 85.78 | 41.50 |
| **MSC** | 3E+07 | 4.96G | 2E+07 | 2.86G | 57.74 | 97.06 | 87.98 | 56 |
| **iPSC** | 1E+08 | 17.79G | 7E+07 | 9.77G | 54.89 | 98.15 | 89.47 | 69 |

**Branscome et al. Supplementary Table 1**

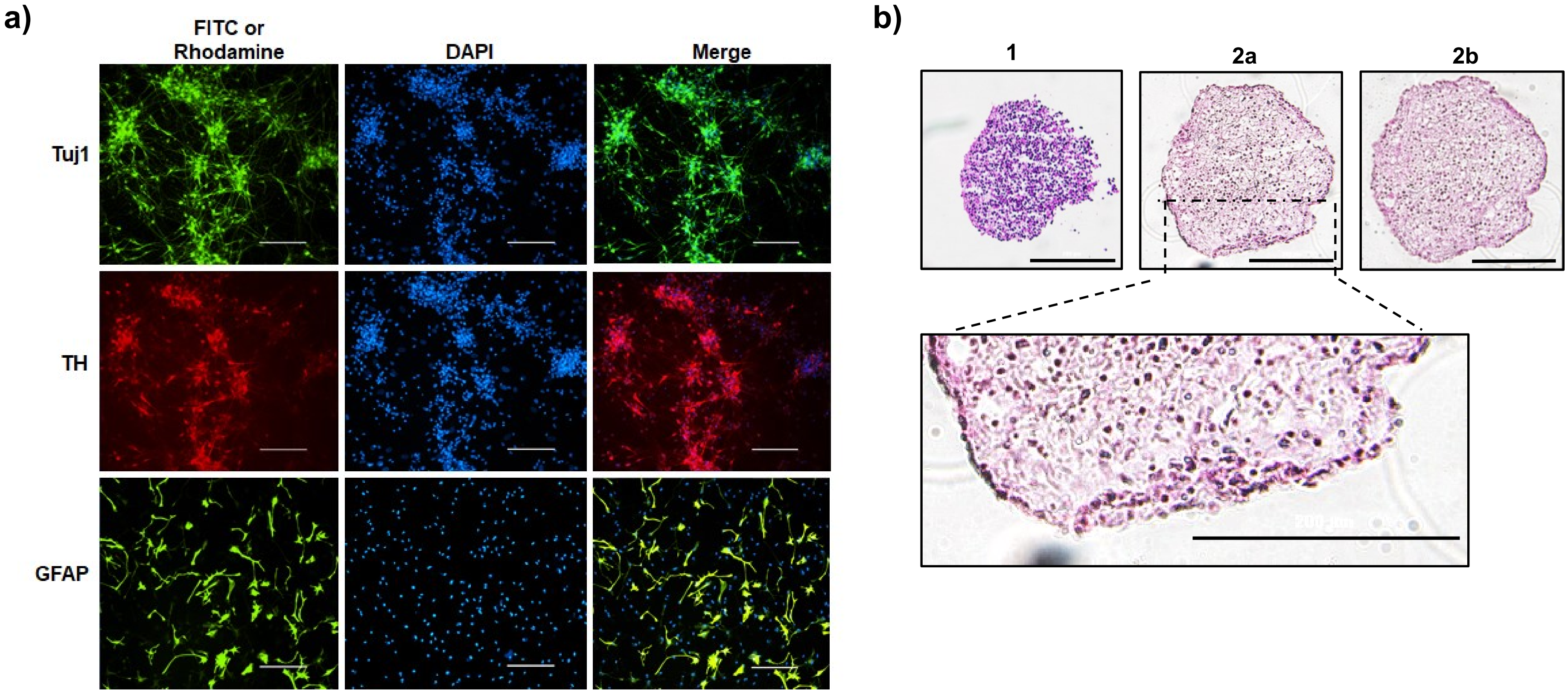


**Supplementary Figure 1. (a)** ICC staining of 2D cultures of differentiated NPCs. Differentiated NPCs were incubated with antibodies against immature neurons (Tuj1), dopaminergic neurons (TH), and astrocytes (GFAP). Fluorescent images show the relative expression of Tuj1, TH, and GFAP. Nuclei were counterstained with DAPI. Scale bar = 400 µm. **(b)** H&E staining was performed on cross-sectioned, differentiated NPC-derived neurospheres. Representative images from two different neurospheres are shown. Images 2a and 2b represent different cross-sections generated from the same neurosphere.

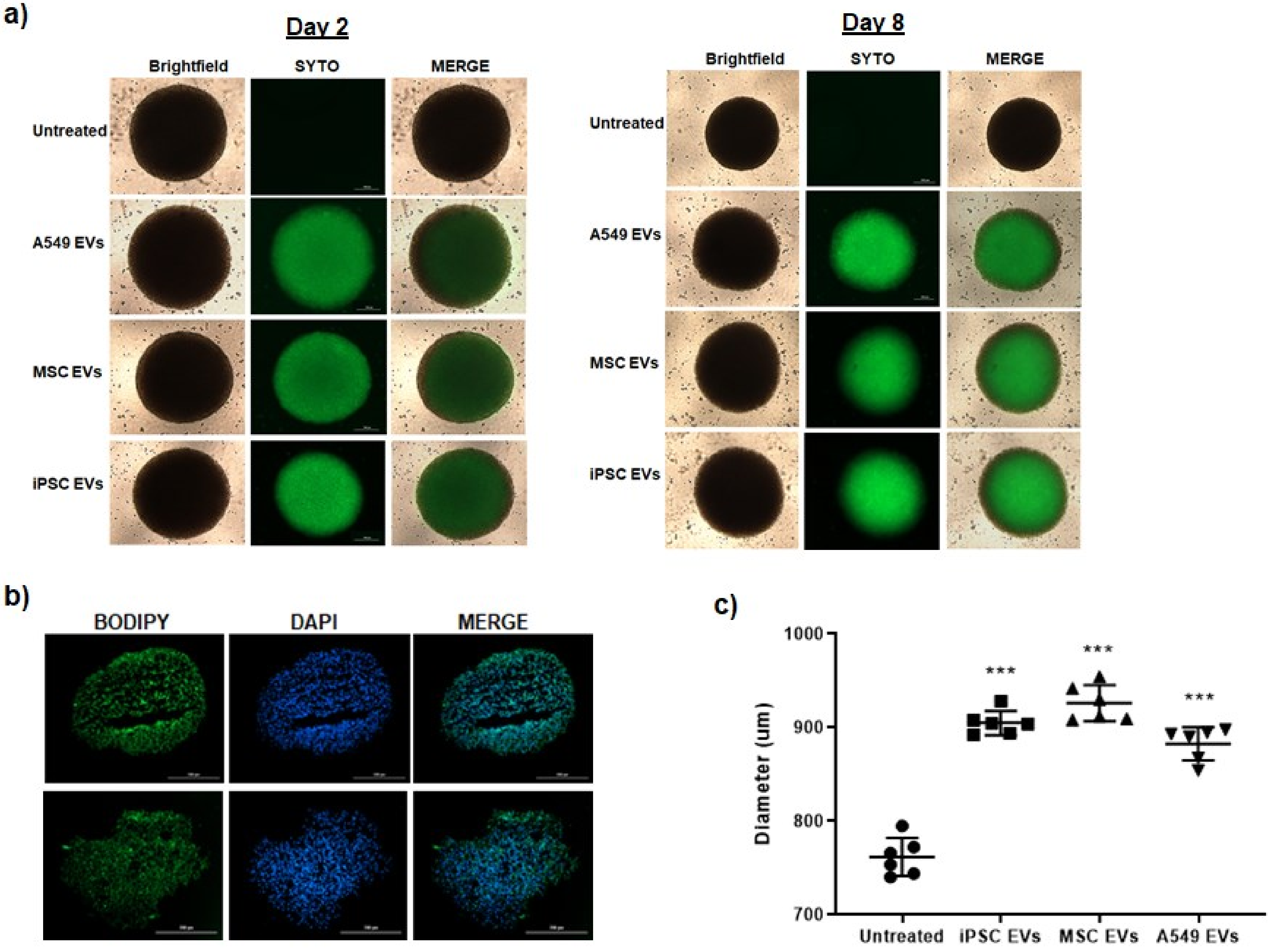


**Branscome et al. Supplementary Figure 2a-c**

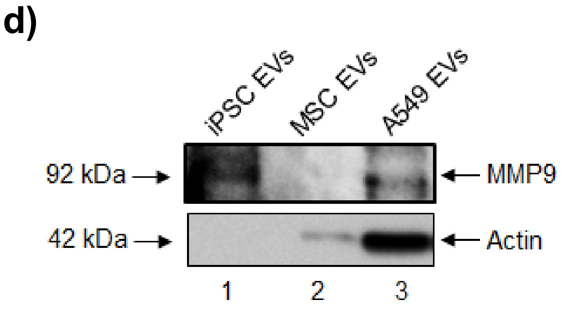


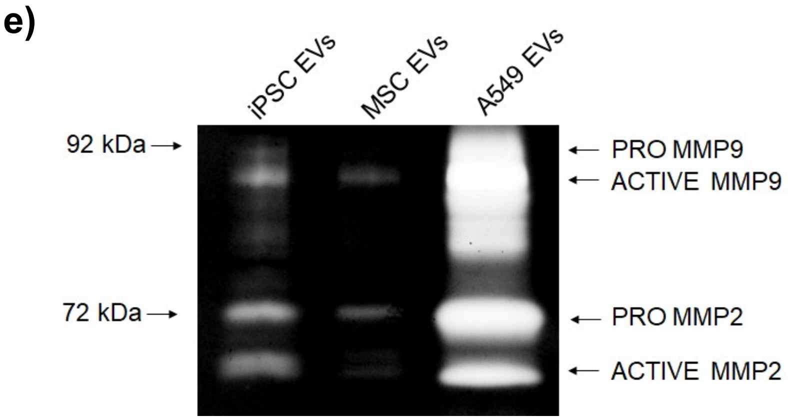


**Supplementary Figure 2. (a)** EVs were fluorescently labeled with SYTO RNASelect Green Fluorescent Stain. After removal of excess dye, EVs were added to differentiated neurospheres (day 0) at an approximate ratio of 1:250 (recipient cell to EV ratio). After 24 hours media was completely replaced. Representative brightfield and fluorescent images show the relative uptake of EVs on days 2 and 8. Scale bar = 200 µm. n = 3. **(b)** Fluorescent images of cross-sectioned, iPSC EV-treated neurospheres show the relative distribution of fluorescently labeled (BODIPY) EVs. Scale bar = 200 µm. **(c)** The average diameter (two measurements per sphere) of EV-treated neurospheres was measured using calibrated imaging software. n = 3. *** $p < 0.0001$ relative to untreated. **(d)** Western blot was performed to evaluate the relative expression of MMP9 in each EV preparation. **(e)** Gelatin zymography was performed to examine the proteolytic activity of EVs. Imaging of the stained gel shows areas of gelatin degradation corresponding to both MMP9 and MMP2 activity.

**Branscome et al. Supplementary Figure 2d-e**

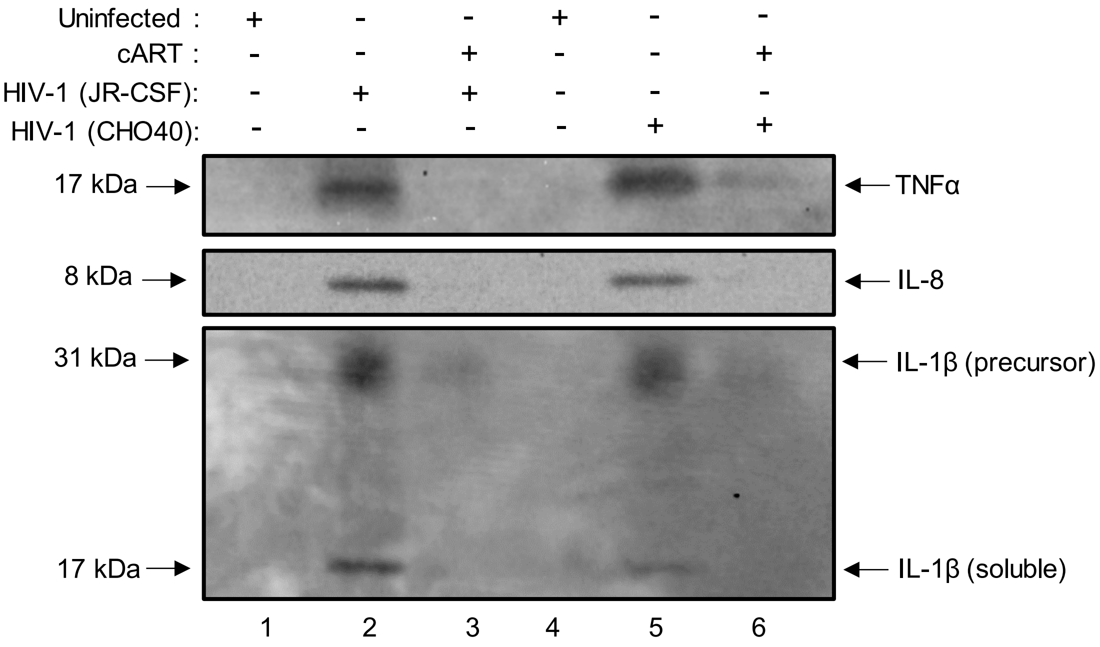


**Supplementary Figure 3.** Differentiated neurospheres were exposed to HIV-1 (JR-CSF and CHO40; MOI:10) with or without cART (lamivudine, tenofovir disoproxil fumarate, emtricitabine, indinavir) for a period of seven days. Western blot was performed on neurosphere supernatants to assess the relative expression of TNFα, IL-8, and IL-1β.

### Branscome et al. Supplementary Figure 3

a)

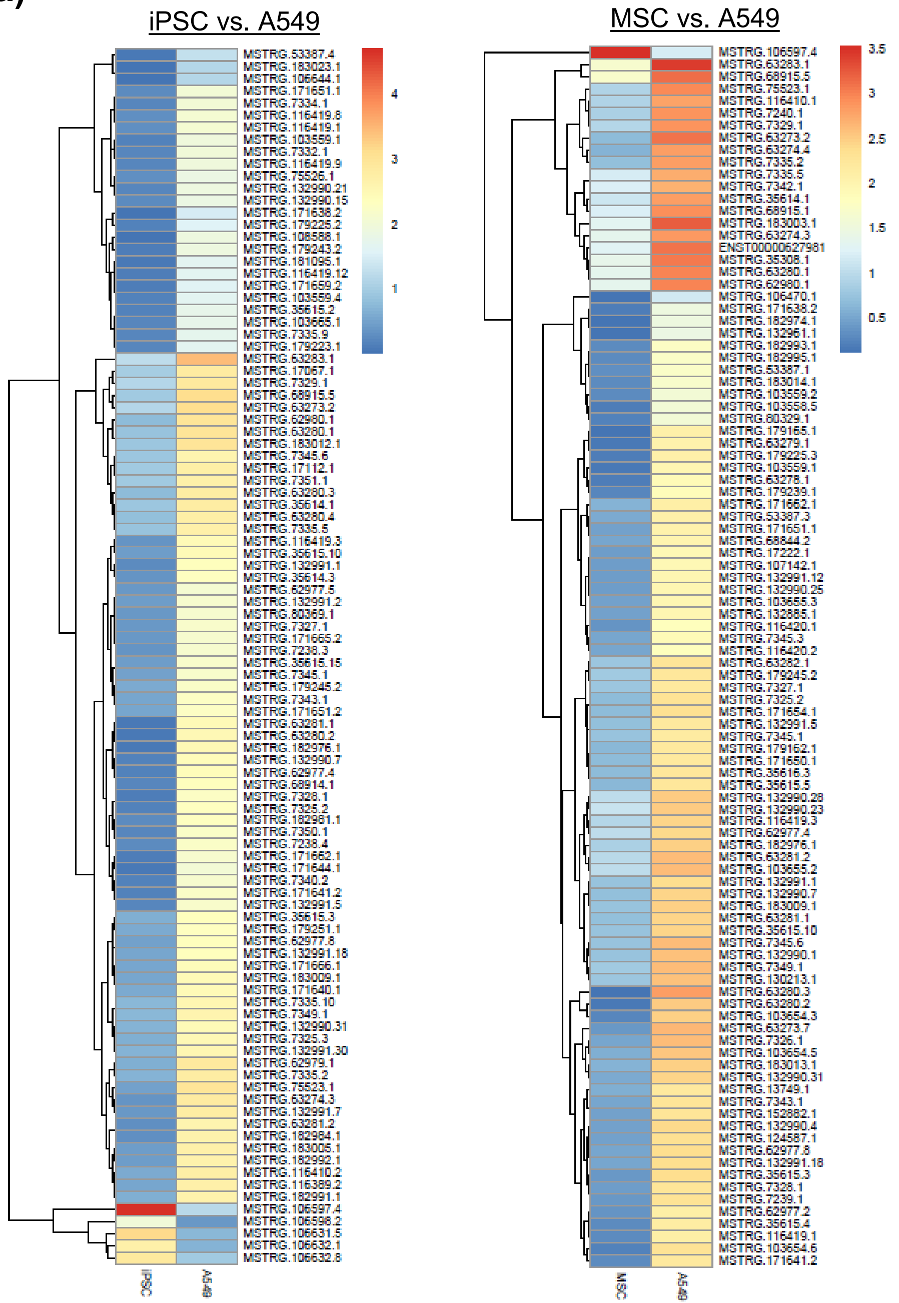


**Branscome et al. Supplementary Figure 4a**

b) iPSC vs. A549

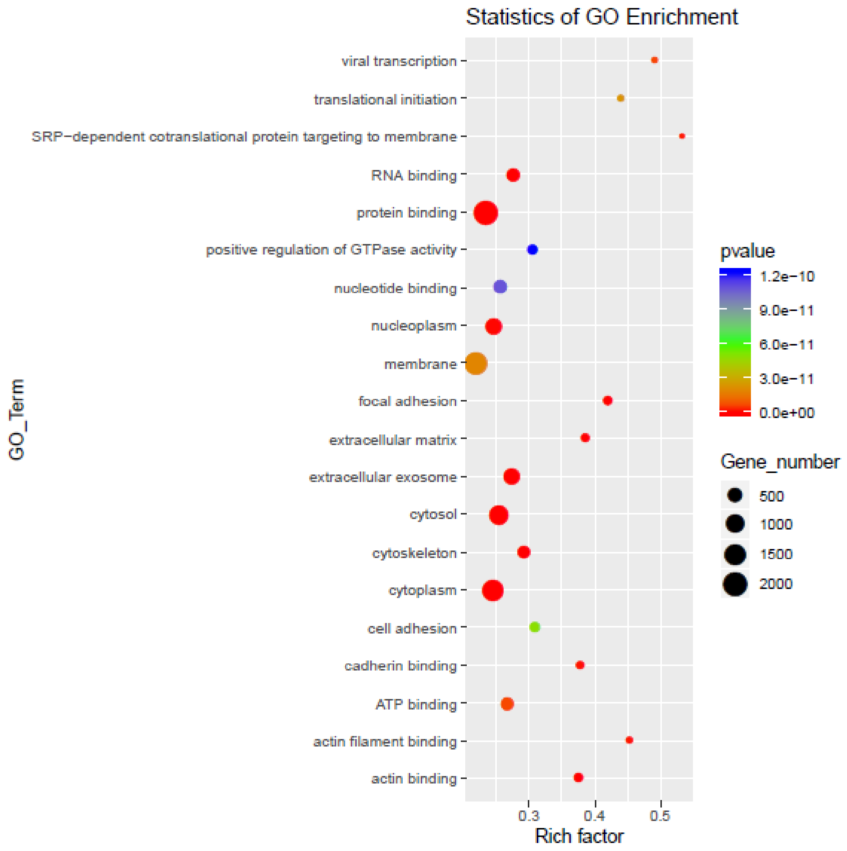


c) MSC vs. A549

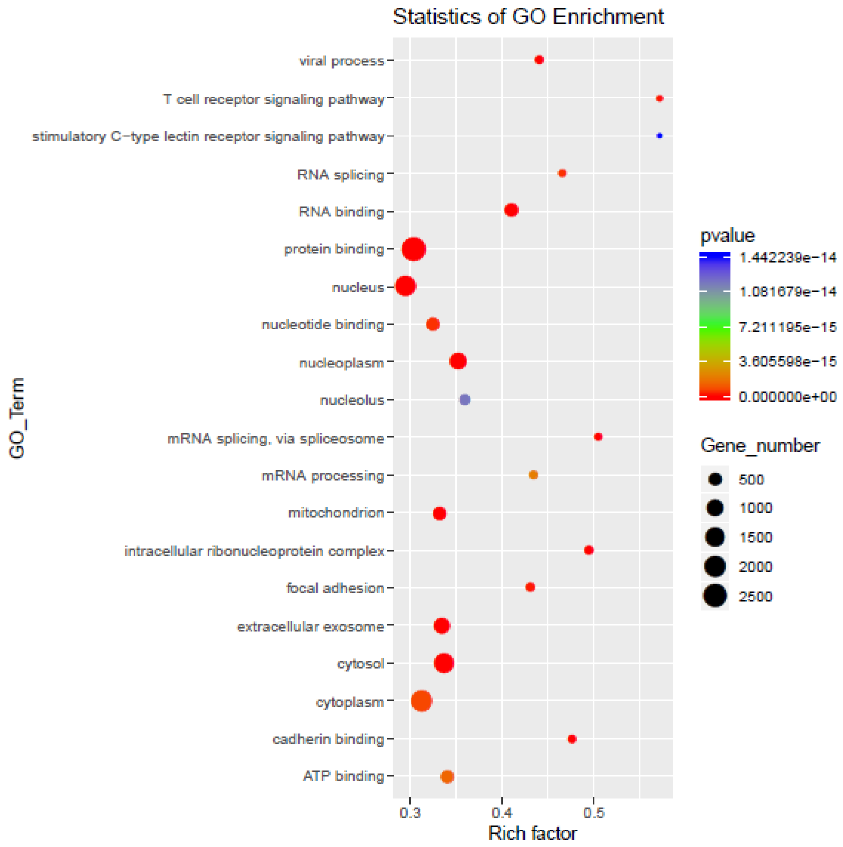


**Branscome et al. Supplementary Figure 4b-c**

d)

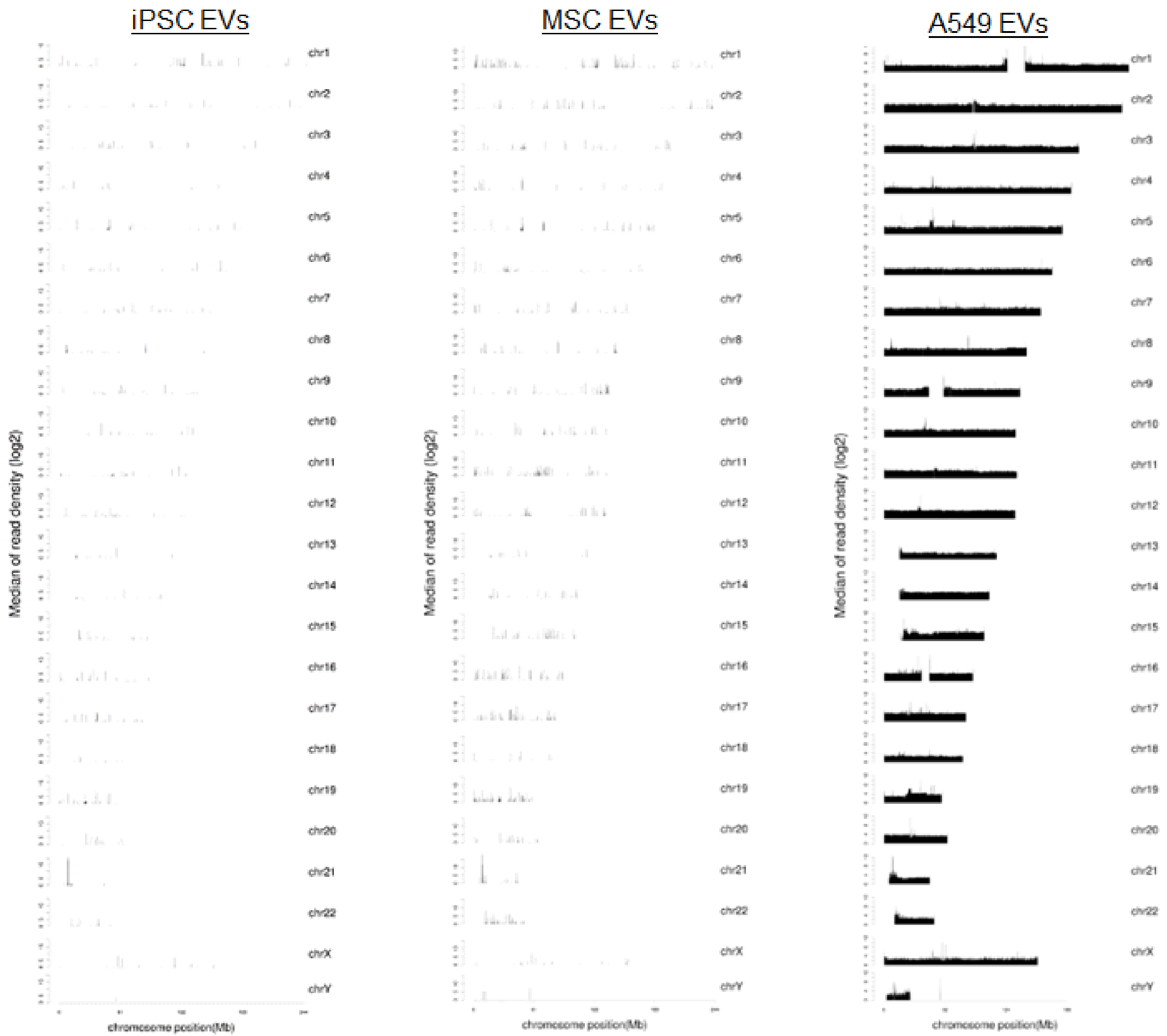


**Supplementary Figure 4**. **(a)** Heat maps comparing lncRNA transcripts between iPSC and A549 EVs (left panel) and MSC and A549 EVs (right panel). Statistics of lncRNA GO enrichment analysis for **(b)** iPSC vs. A549 EVs and **(c)** MSC vs. A549 EVs. Plots depict the p value/gene number for the corresponding GO term enrichment. **(d)** Chromosome maps show the relative density distribution of EV-associated transcripts.

**Branscome et al. Supplementary Figure 4d**

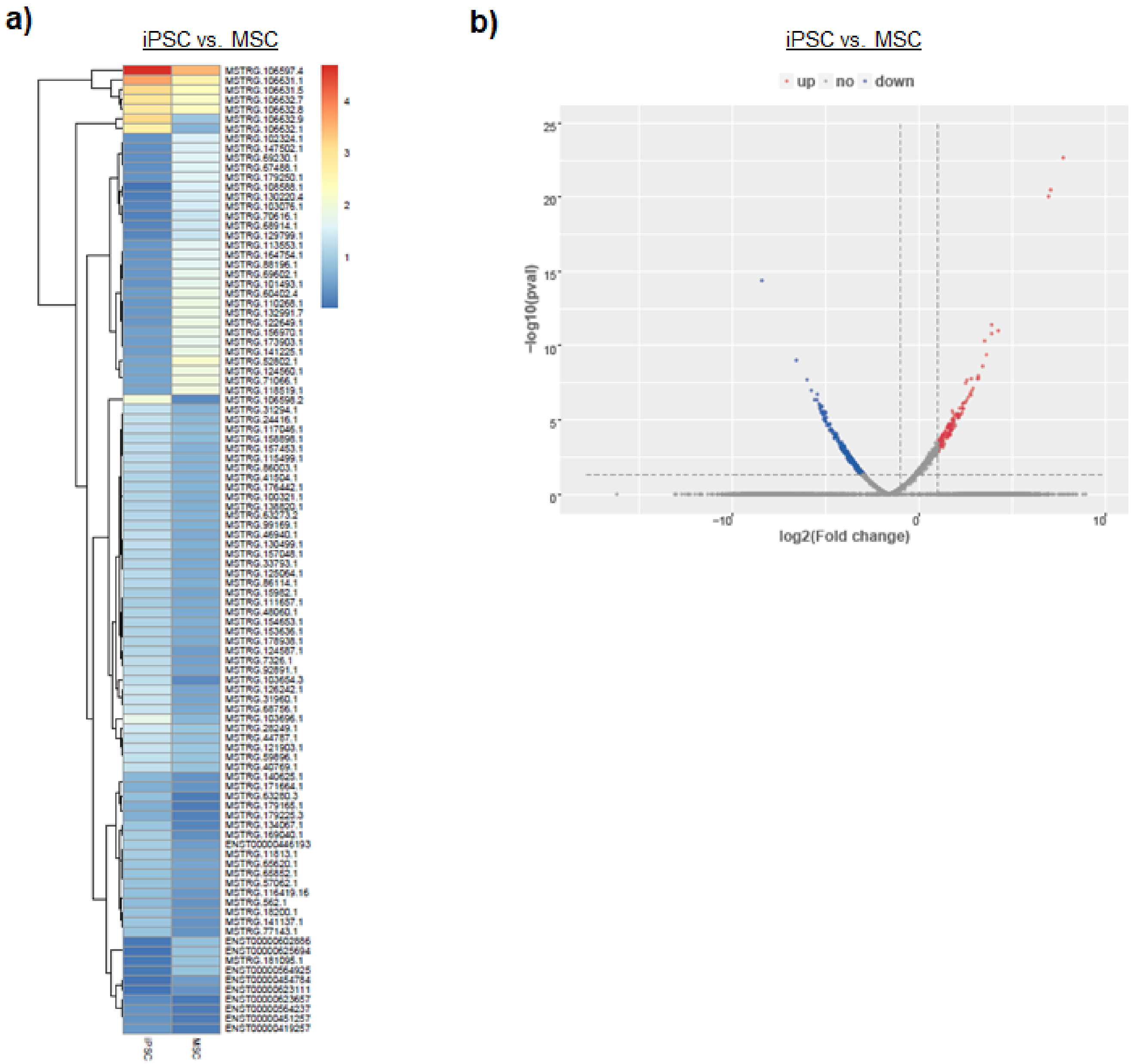


**Supplementary Figure 5. (a)** Heat maps comparing lncRNA transcripts between iPSC and MSC EVs. **(b)** Volcano plots displaying the differentially expressed transcripts between iPSC and MSC EVs.

### Branscome et al. Supplementary Figure 5

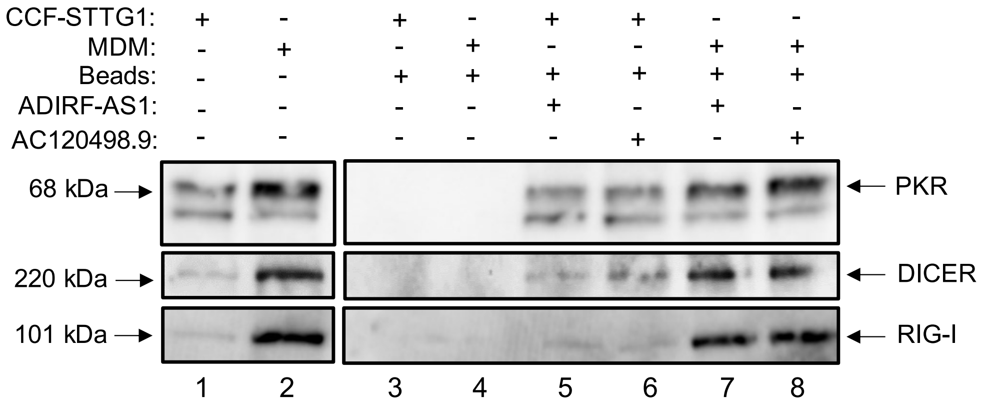


**Supplementary Figure 6.** RNA pulldown assay. Biotin-conjugated synthetic RNA sequences corresponding to the sequences of interest from ADIRF-AS1 and AC120498.9 were incubated with either CCF-STTG1 or MDM extract. Western blot was performed to assess the expression of different RNA binding proteins (PKR, Dicer, RIG-I).

**Branscome et al. Supplementary Figure 6**

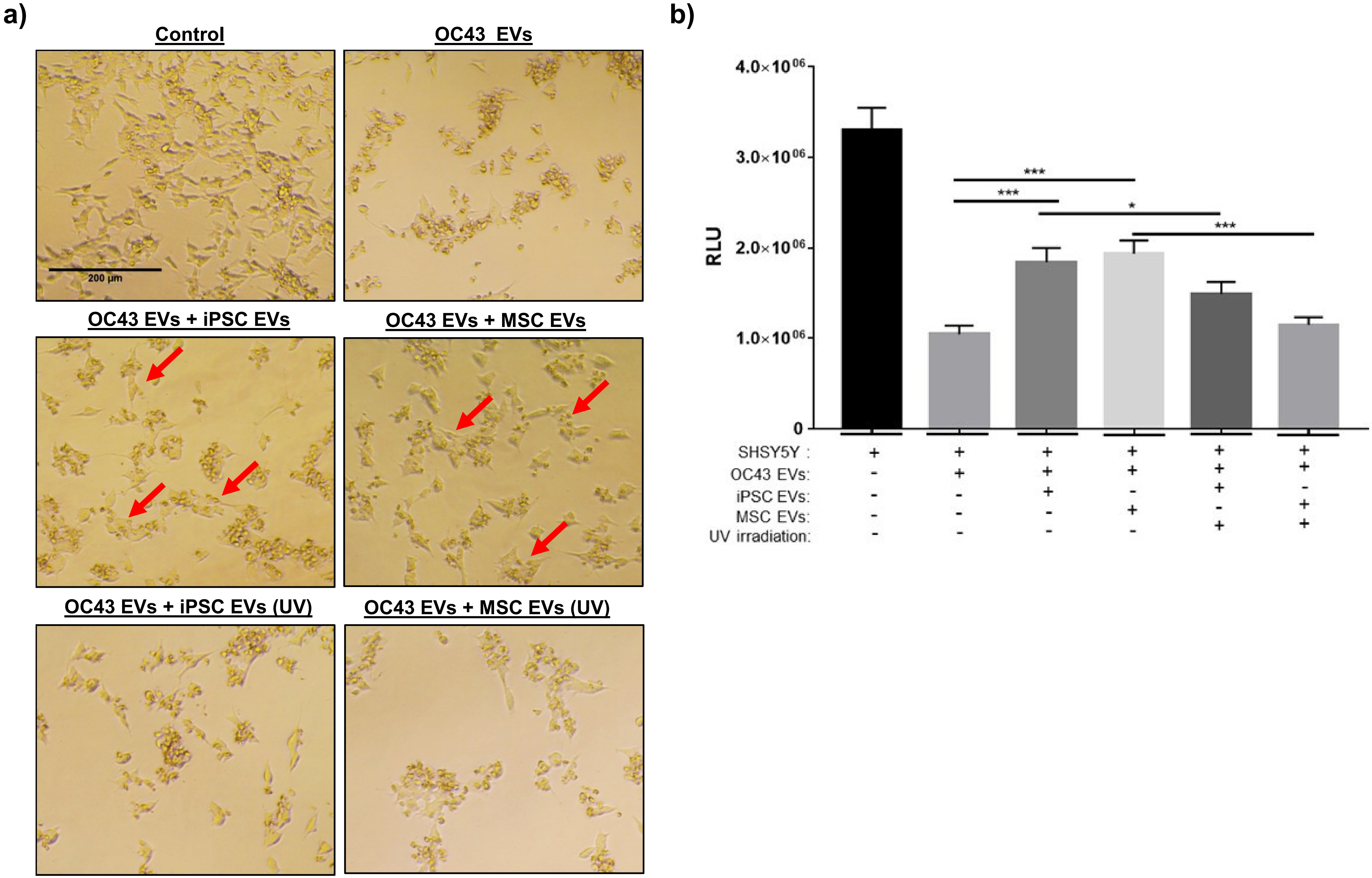


**Supplementary Figure 7.** Cellular viability assay. SHSY5Y cells were exposed to EVs from OC43 infected cells with or without treatment of stem cell EVs at an approximate ratio of 1:1000 (recipient cell to EV ratio). Stem cell EVs were exposed to UV(C) to inactivate EV-associated RNAs. **(a)** Representative images show the appearance and morphology of cells after 8 days in culture. Scale bar = 200 µm. n = 3. **(b)** Cell viability was quantified via CellTiter-Glo. n=3. * $p < 0.05$, *** $p < 0.0001$.

**Branscome et al. Supplementary Figure 7**

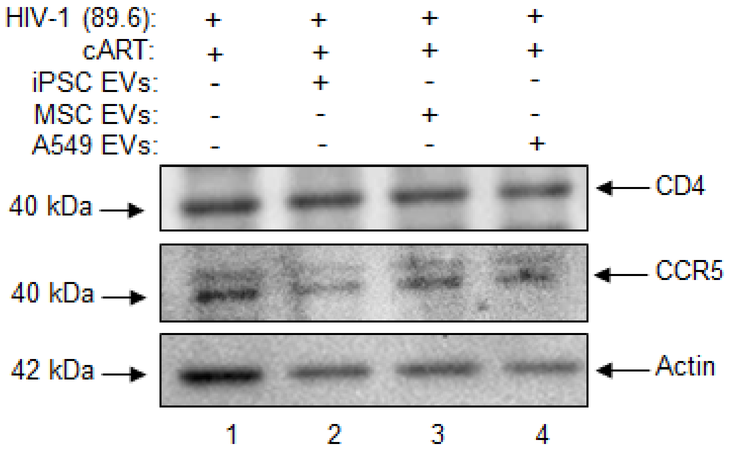


**Supplementary Figure 8**. Differentiated neurospheres were exposed to dual-tropic HIV-1 89.6 with or without cART (lamivudine, tenofovir disoproxil fumarate, emtricitabine, indinavir) for a period of fourteen days. Additionally, the HIV-1 + cART treated samples were treated with either iPSC, MSC, or A549 EVs. Western blot was performed on neurosphere lysates to evaluate the relative expression of CD4 and CCR5.

**Branscome et al. Supplementary Figure 8**

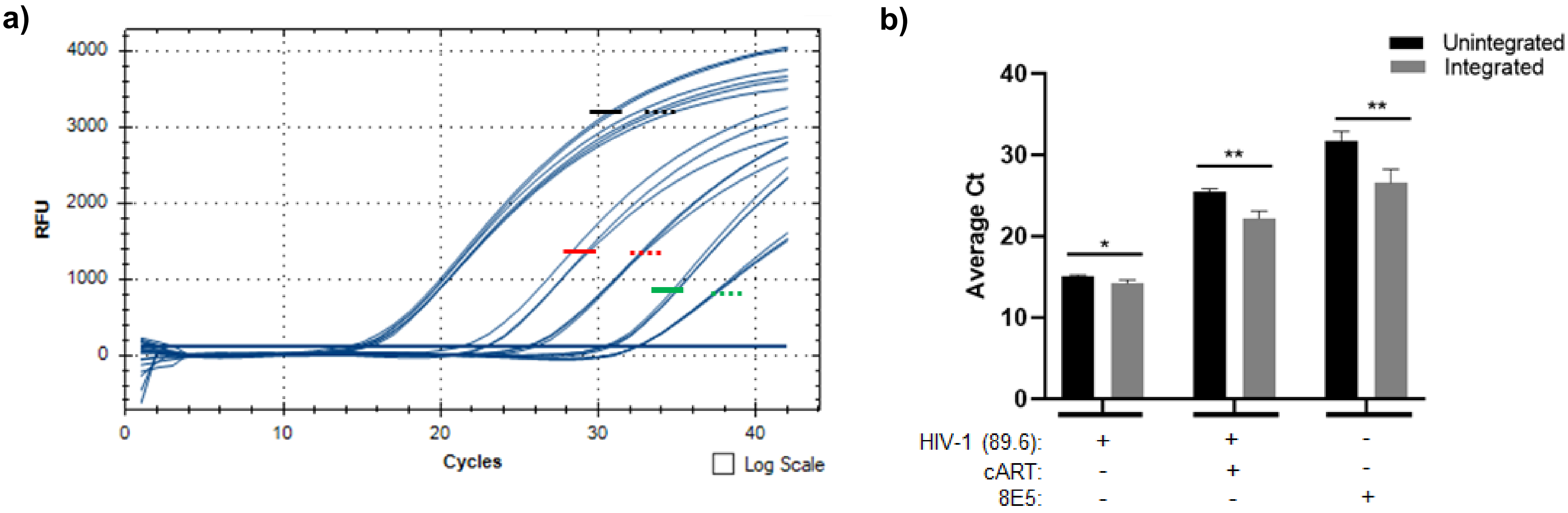


**Supplementary Figure 9.** Integration assay. Differentiated neurospheres were exposed to dual-tropic HIV-1 89.6 with or without cART (lamivudine, tenofovir disoproxil fumarate, emtricitabine, indinavir) for a period of fourteen days. **(a)** DNA from each sample (100 ng) was used to perform PCR using Alu-*gag* (integrated) or *gag*-only (unintegrated) primers, followed by a second quantitative PCR using nested primers and probes targeting the HIV-1 LTR region. DNA from the HIV-1 infected cell line 8E5 (100 ng) was used as a positive control. Ct values for the three replicates corresponding to each sample are shown. Ct values corresponding to integrated DNA are denoted by solid lines and Ct values corresponding to unintegrated DNA are denoted by dashed lines. Black lines represent data from HIV-1 infected neurospheres, red lines represent HIV-1 infected + cART neurospheres, and green lines represent 8E5 positive control cells. **(b)** The bar graph shows the average Ct values shown in panel a. A two-tailed Student's t-test was used to assess significance. n = 3. * $p < 0.05$, ** $p < 0.01$.

**Branscome et al. Supplementary Figure 9**

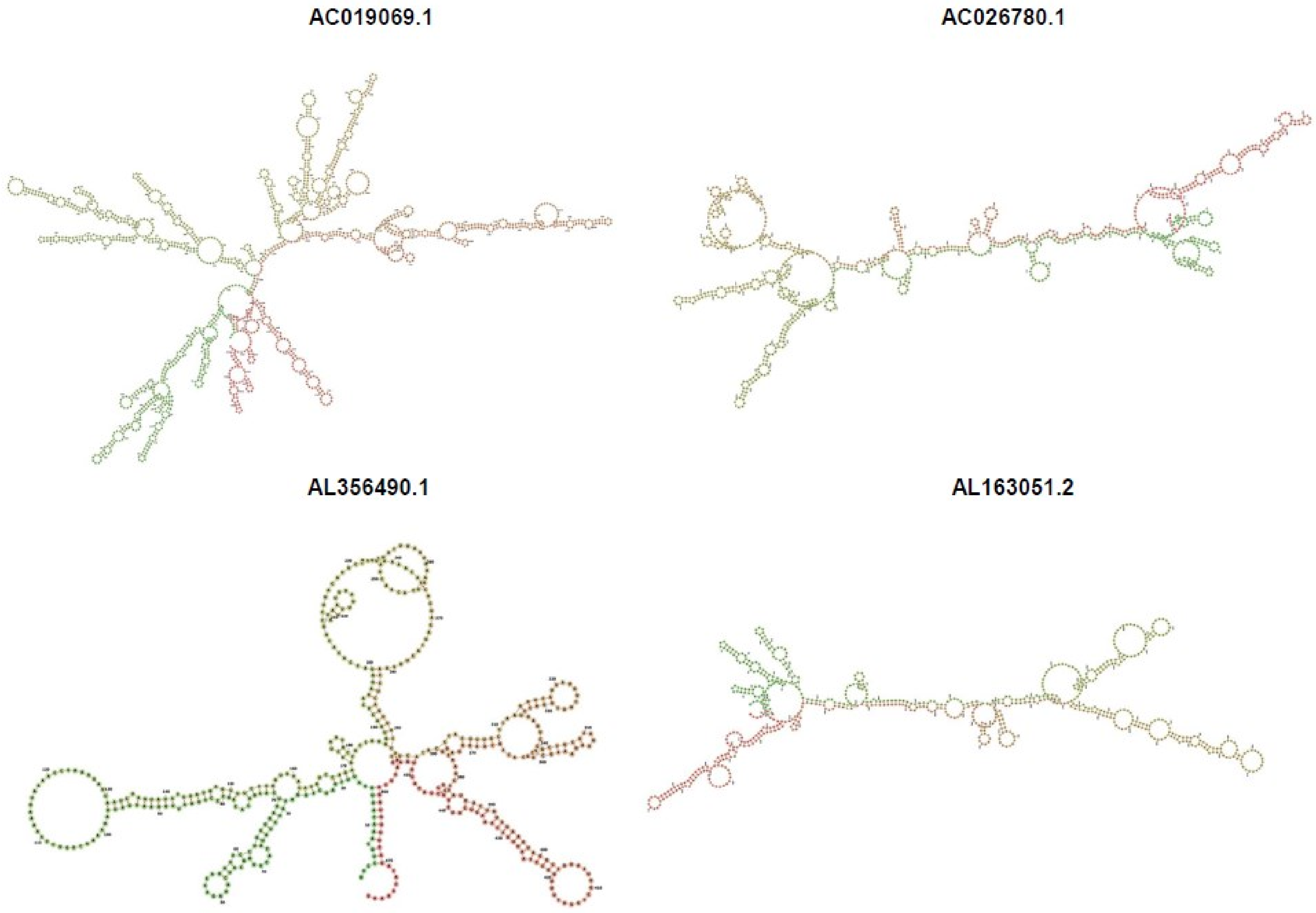


**Branscome et al. Supplementary Figure 10**

**LINC02283**

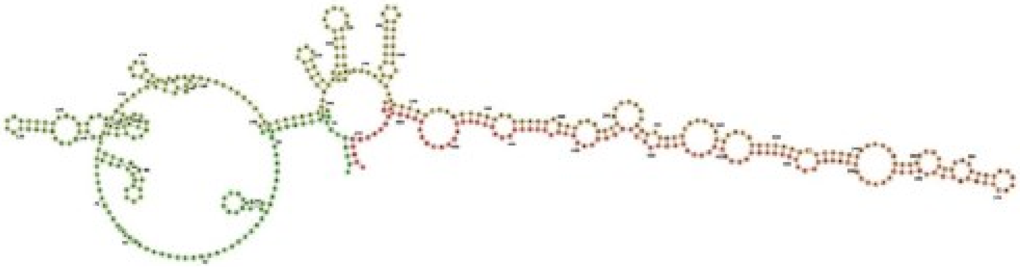

**AL358472.3**

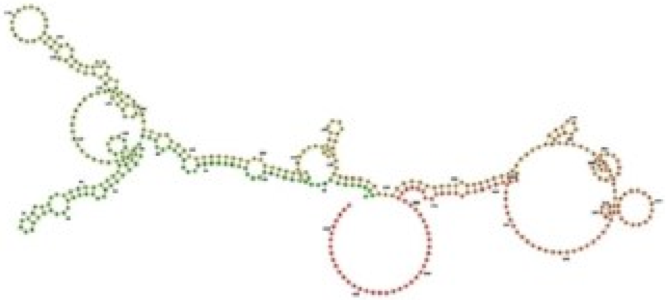

**AC010976.1**

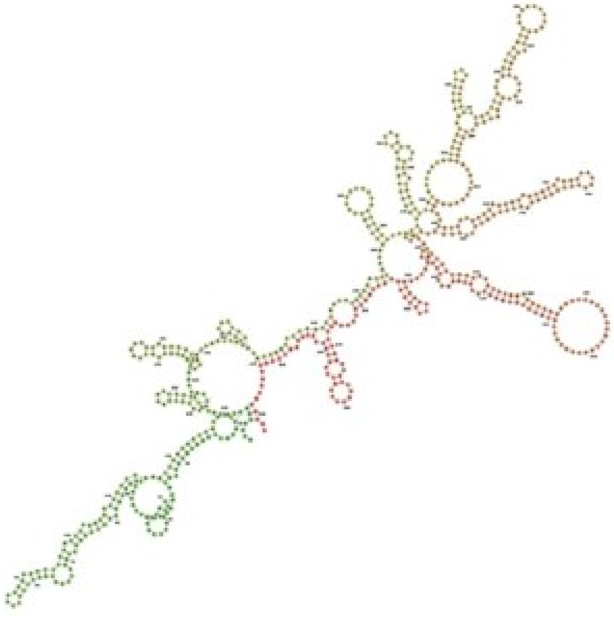

**AL512625.3**

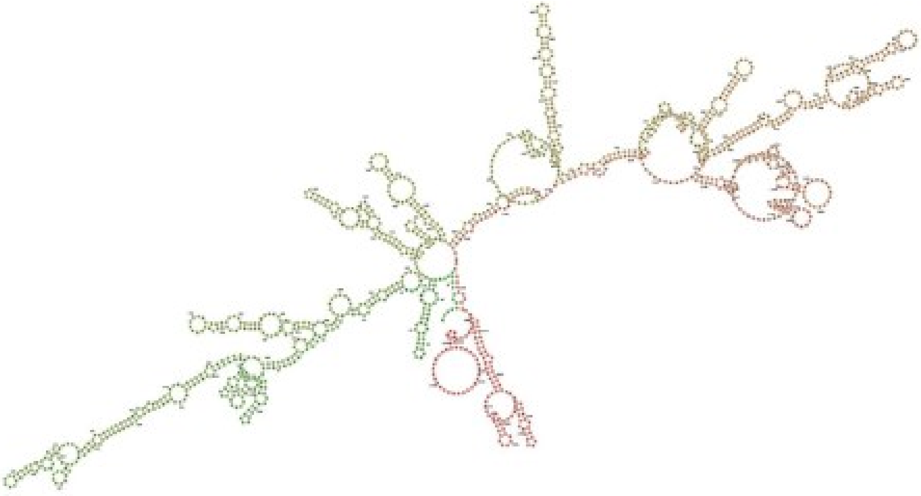

**Branscome et al. Supplementary Figure 10**

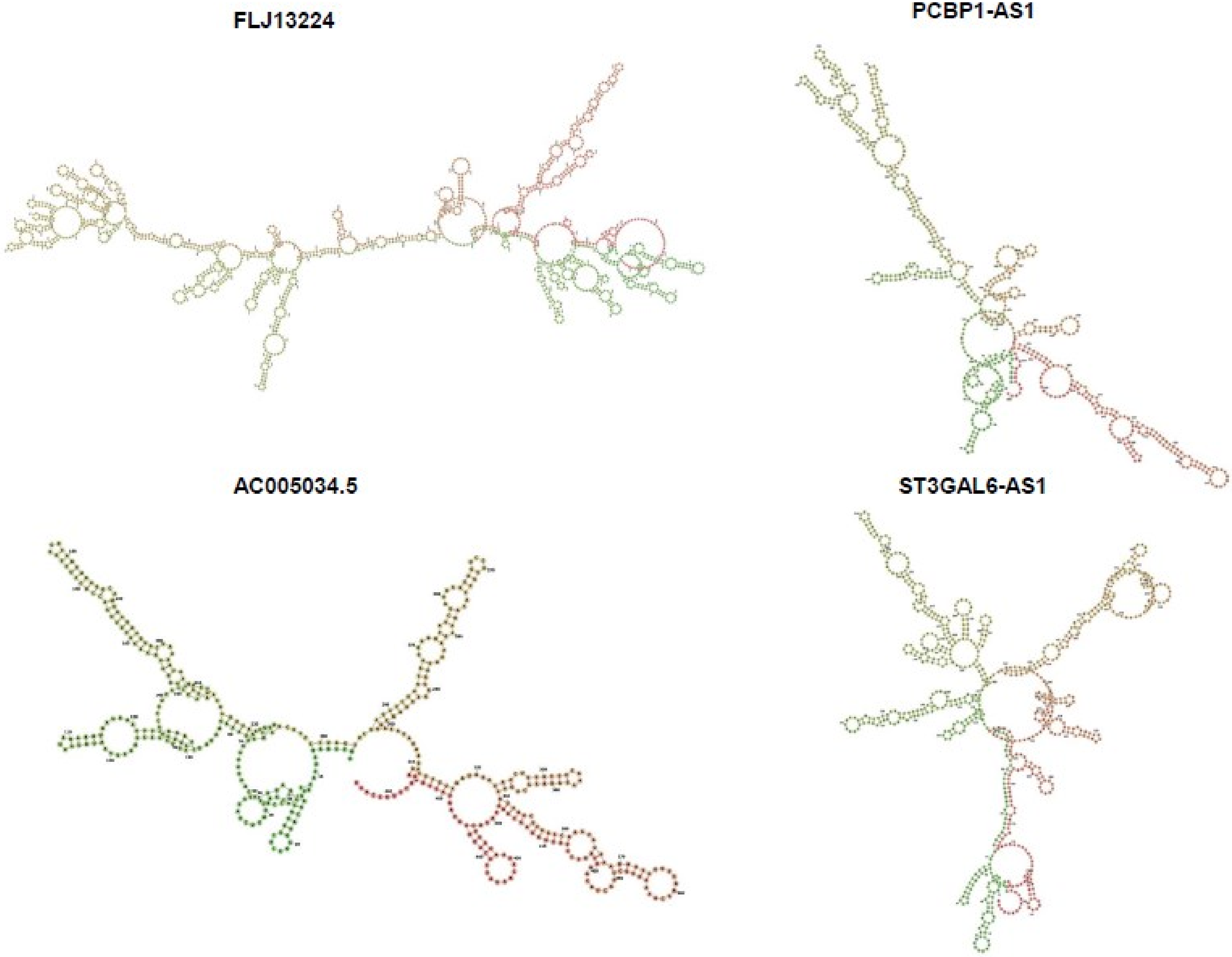


**Branscome et al. Supplementary Figure 10**

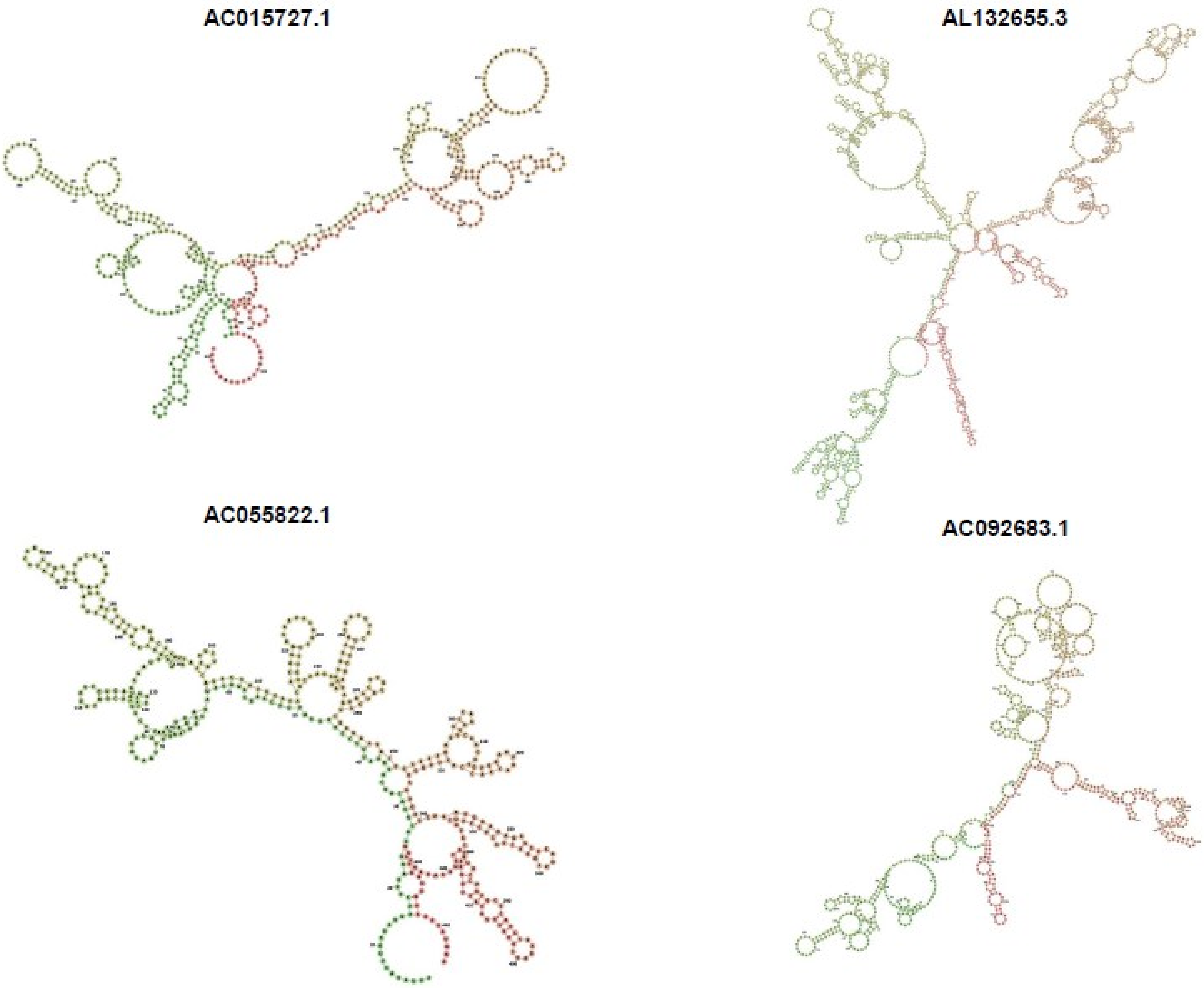


Branscome et al. Supplementary Figure 10

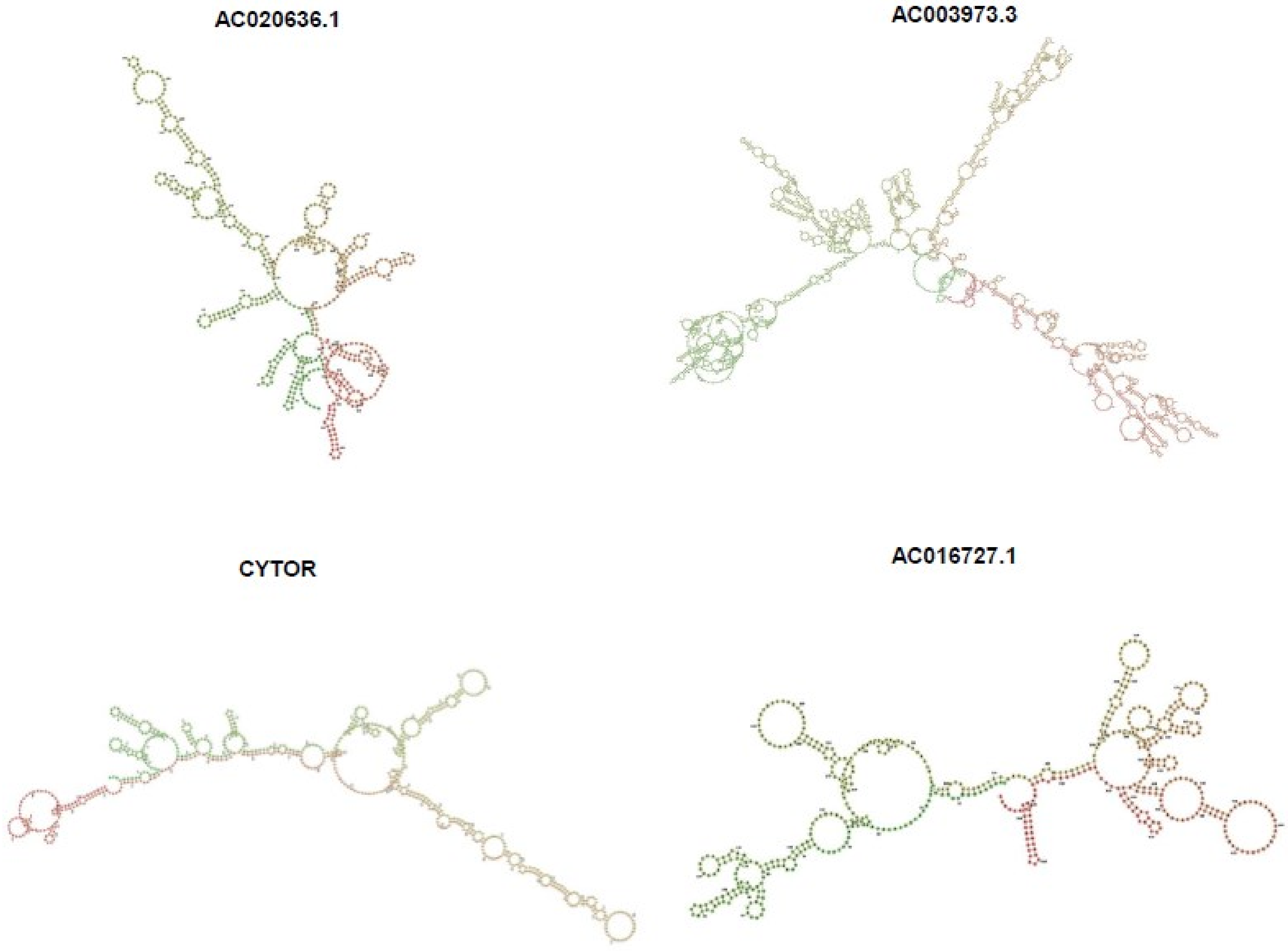


**Branscome et al. Supplementary Figure 10**

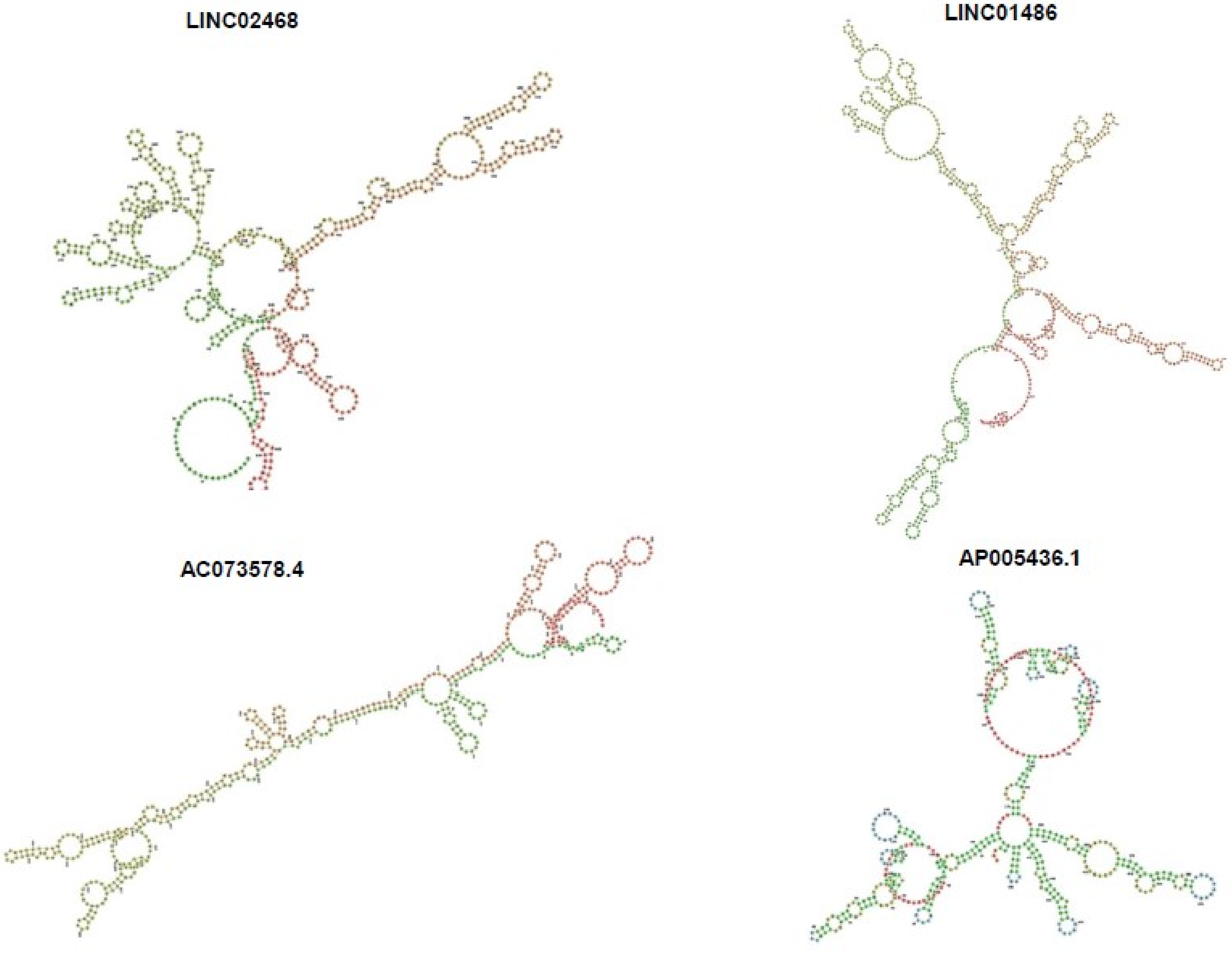


**Branscome et al. Supplementary Figure 10**

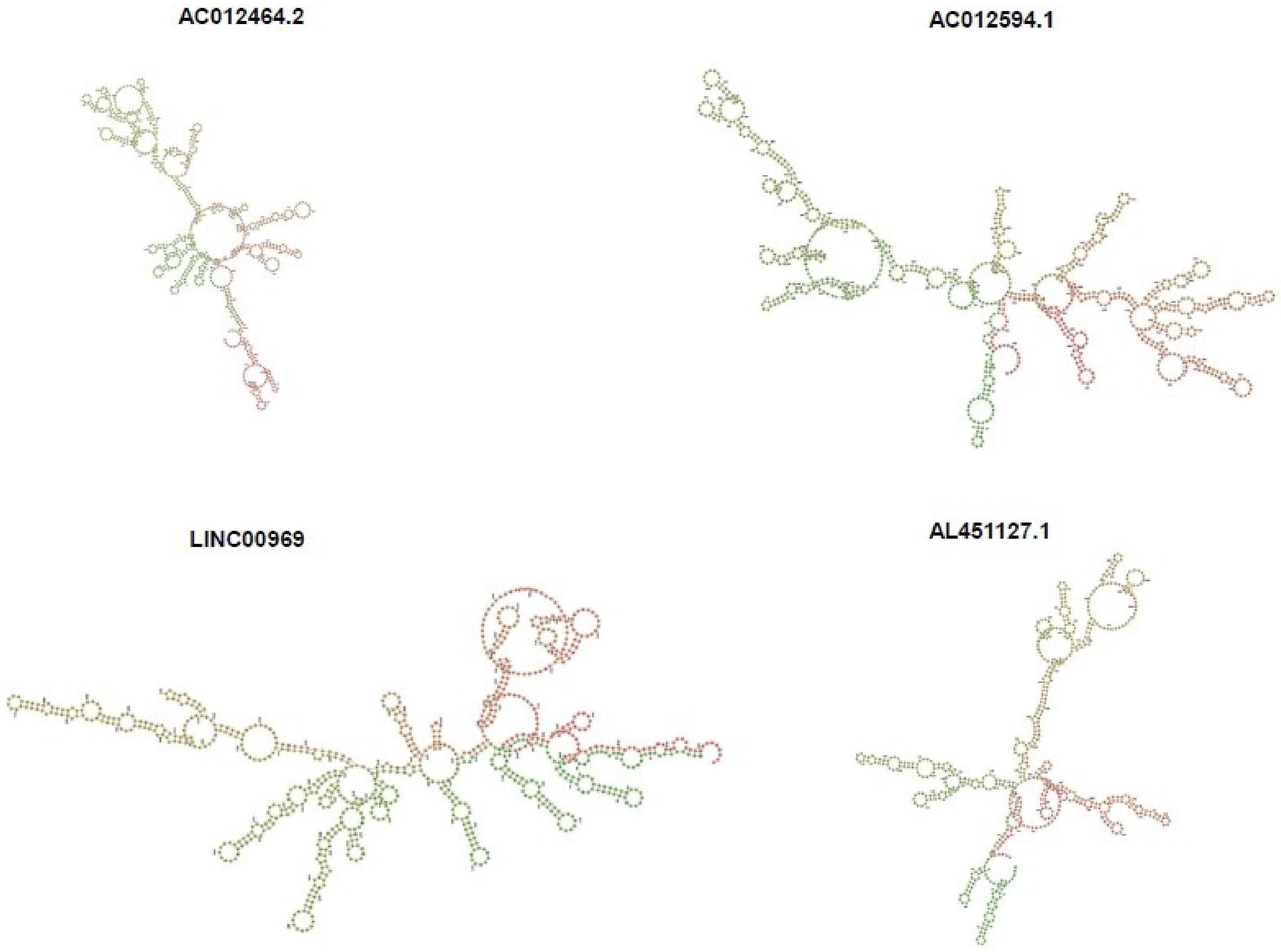


**Branscome et al. Supplementary Figure 10**

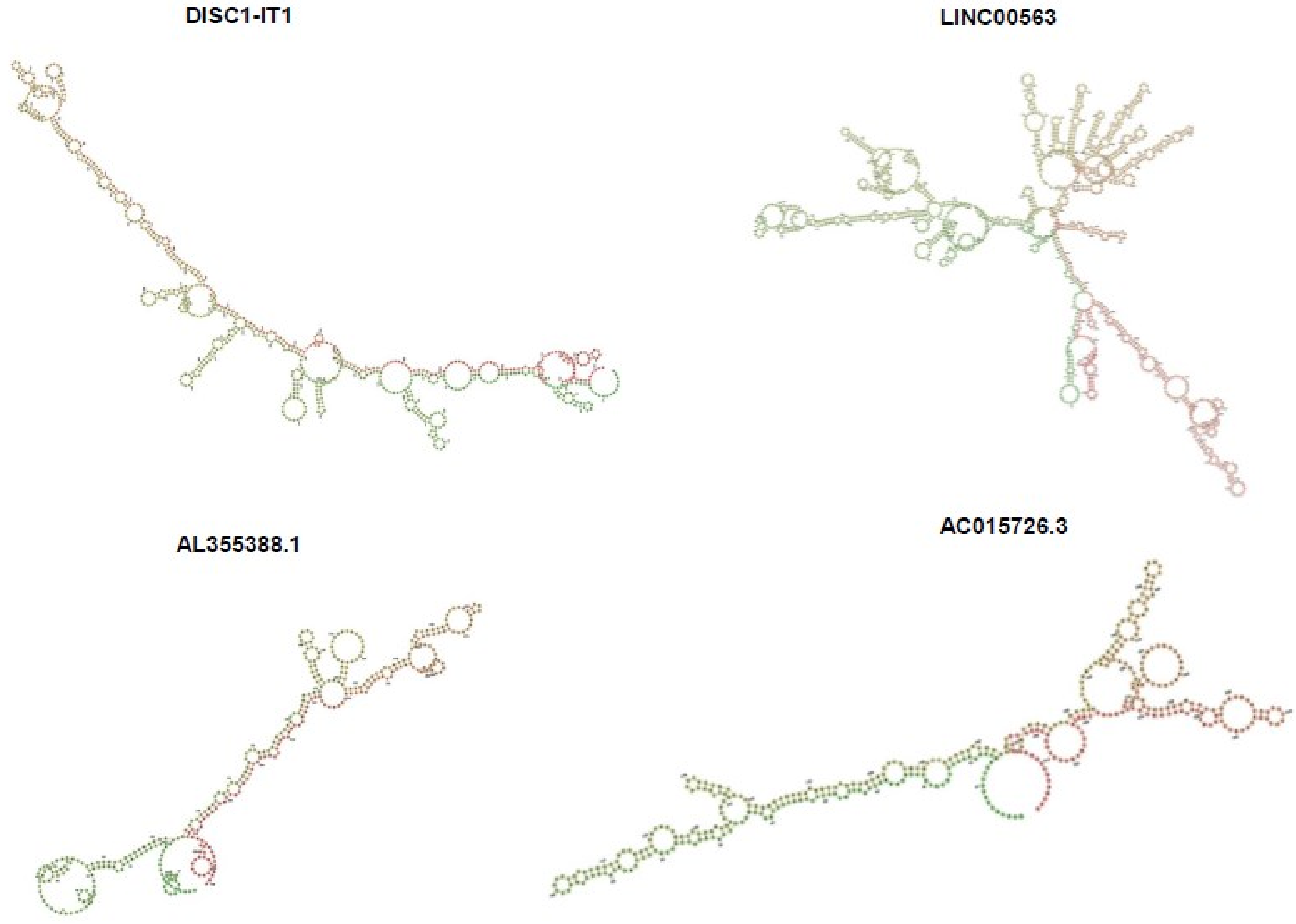


**Branscome et al. Supplementary Figure 10**

**AL353612.1**

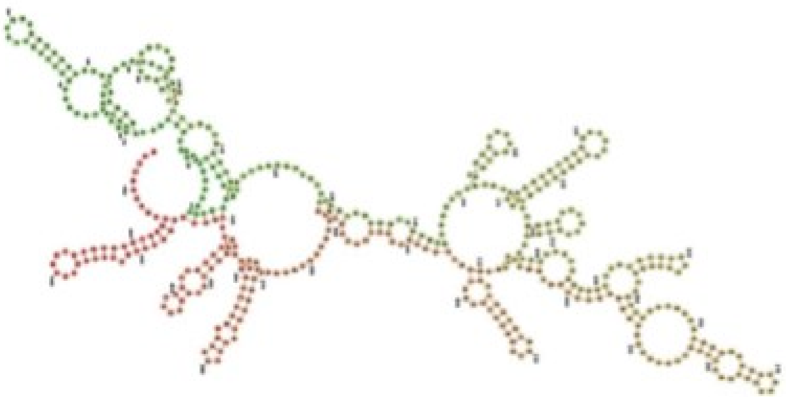

**AZIN1-AS1**

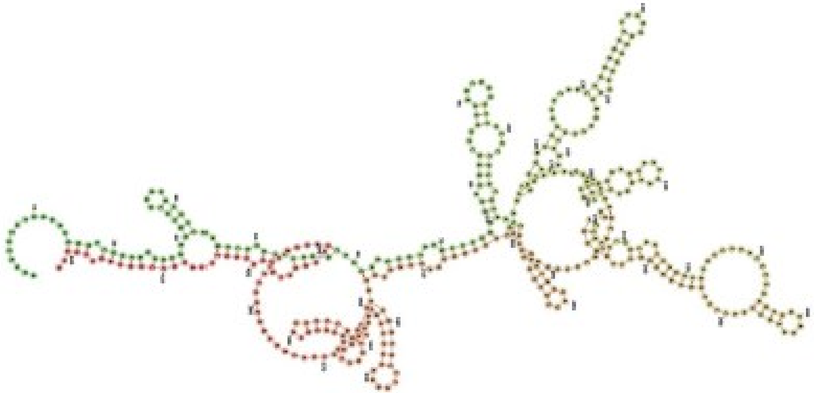

**AC004930.1**

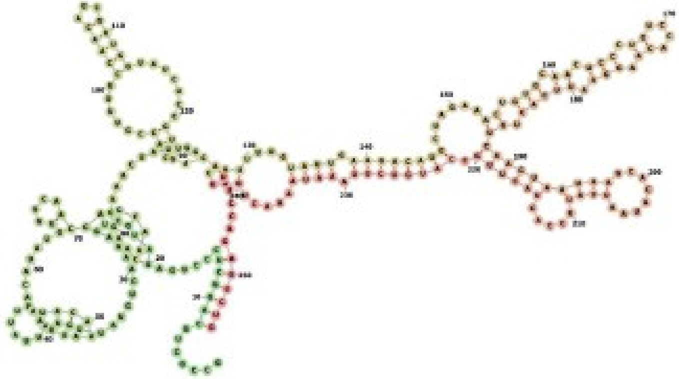

**AL358975.1**

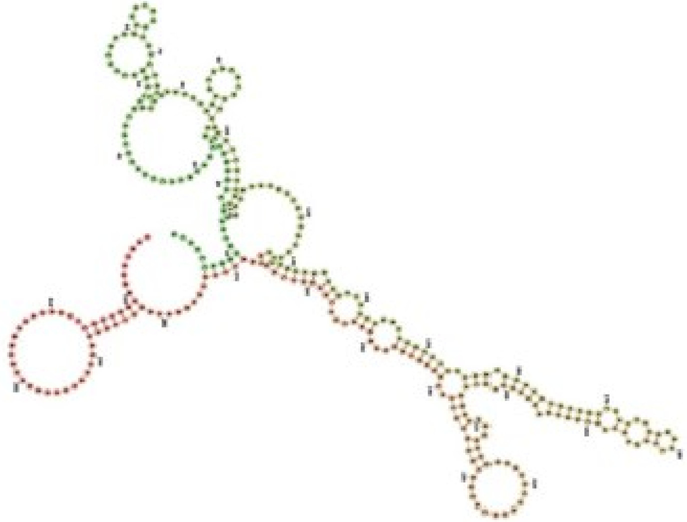

**Branscome et al. Supplementary Figure 10**

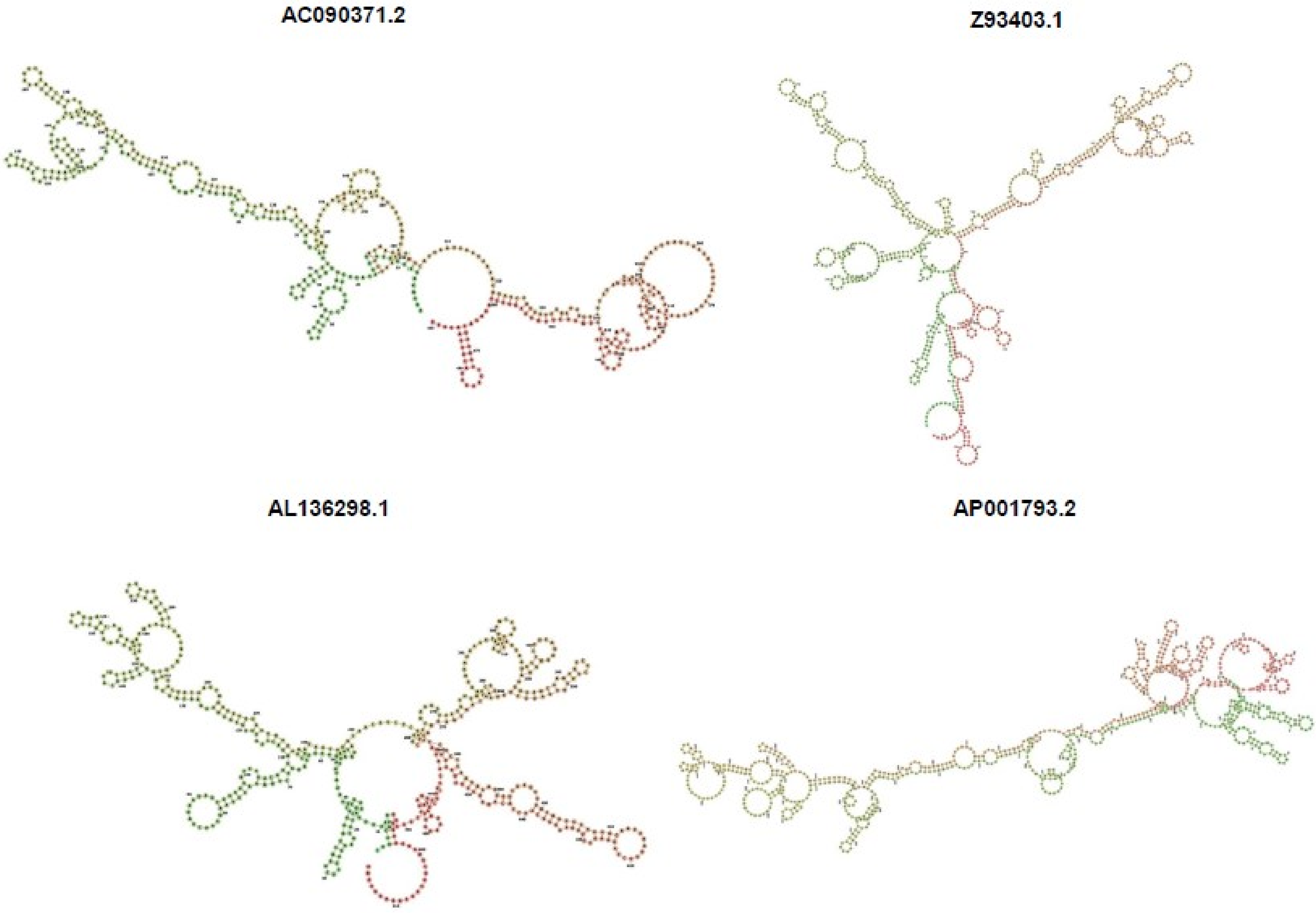


**Branscome et al. Supplementary Figure 10**

**AC120498.9**

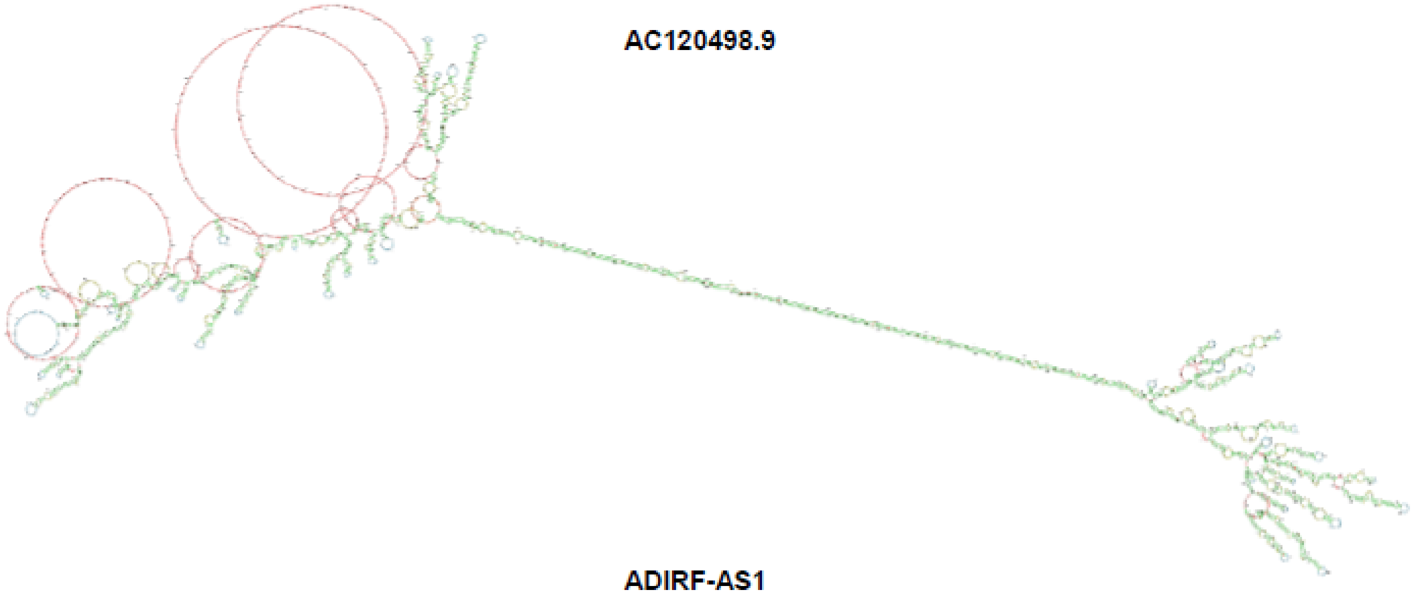

**ADIRF-AS1**

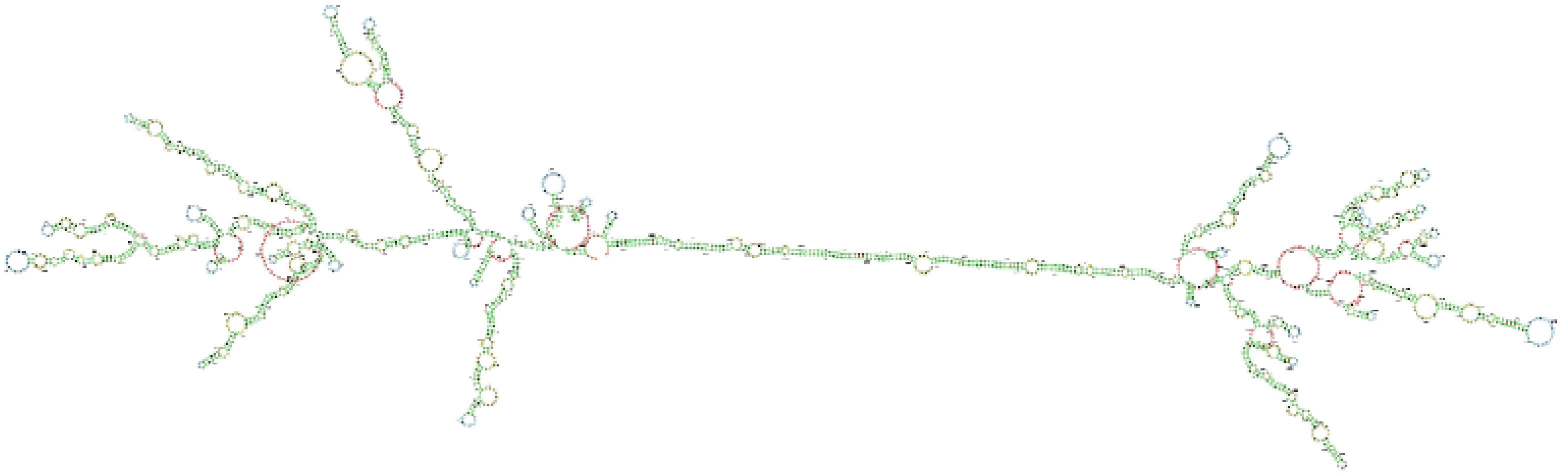

**Supplementary Figure 10.** iPSC EV-associated lncRNAs were sorted based on their corresponding FPKM values (FPKM ≥ 2 relative to MSC and A549 EV-associated lncRNAs). The secondary structures of these lncRNAs were predicted by the RNAfold Server of the ViennaRNA Websuite. Out of approximately 244 lncRNAs, 42 were found to contain multiple circular loops and rings. The secondary structures of those 42 lncRNAs are shown.

**Branscome et al. Supplementary Figure 10**

Pr55/p24 (Fig. 2a)

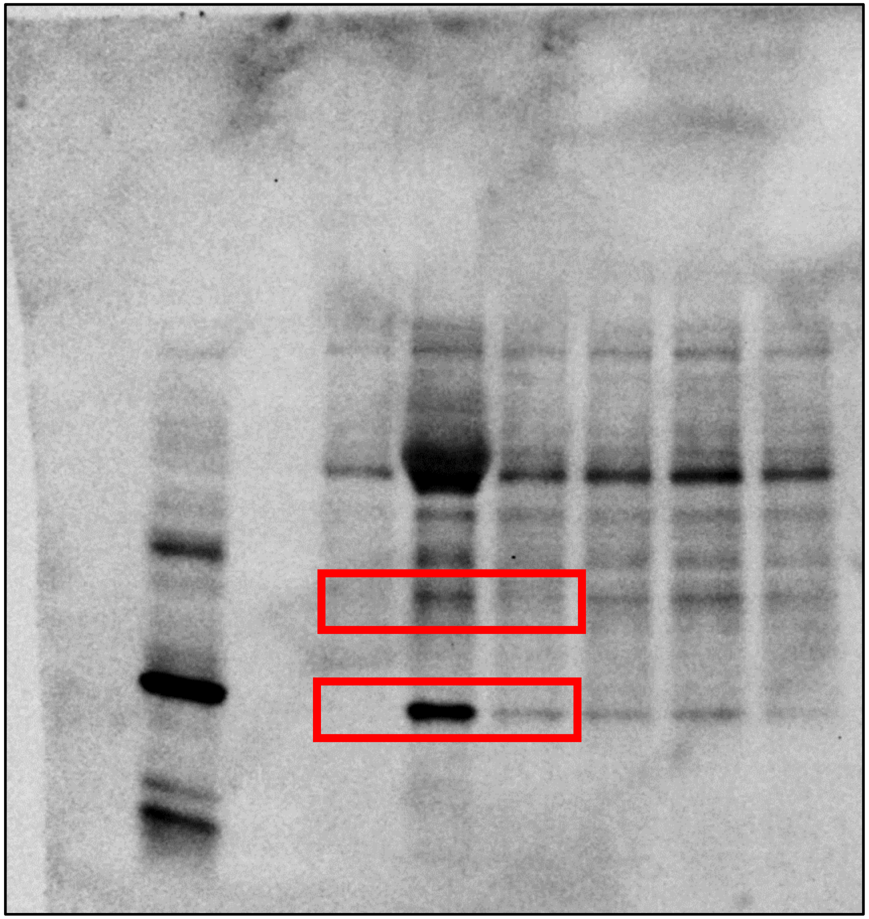

Nef (Fig. 2a)

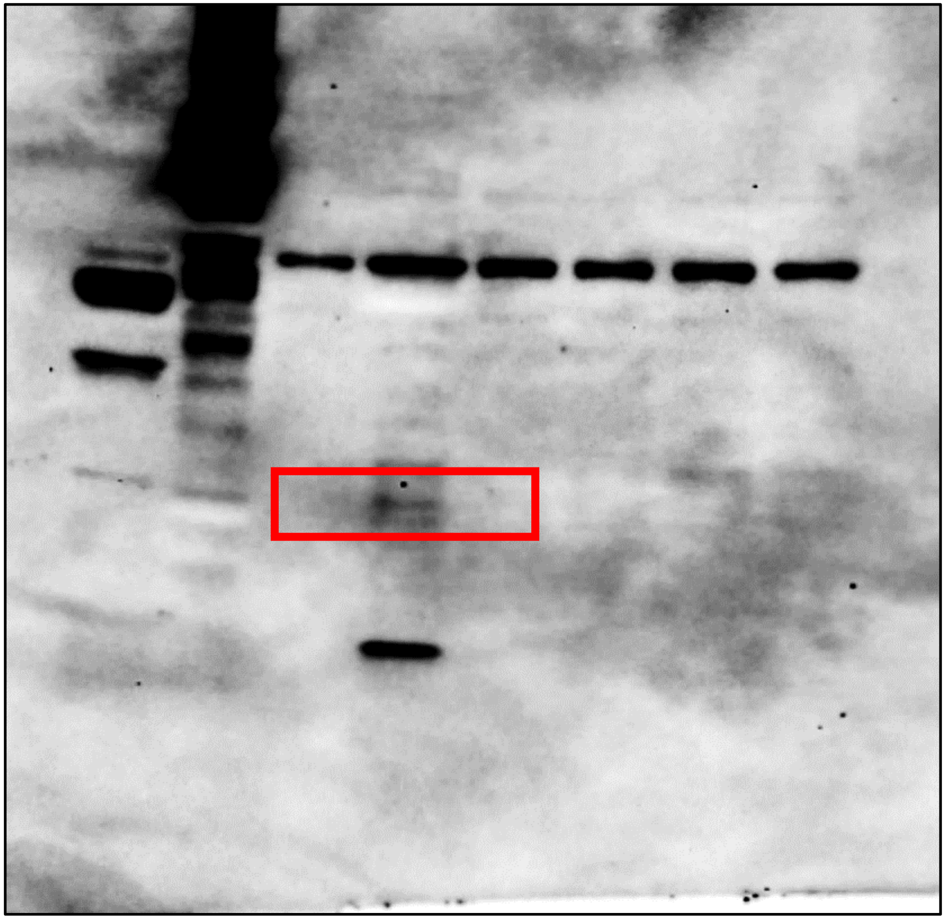

**Branscome et al. Supplementary Figure 11**

Actin (Fig. 2a)

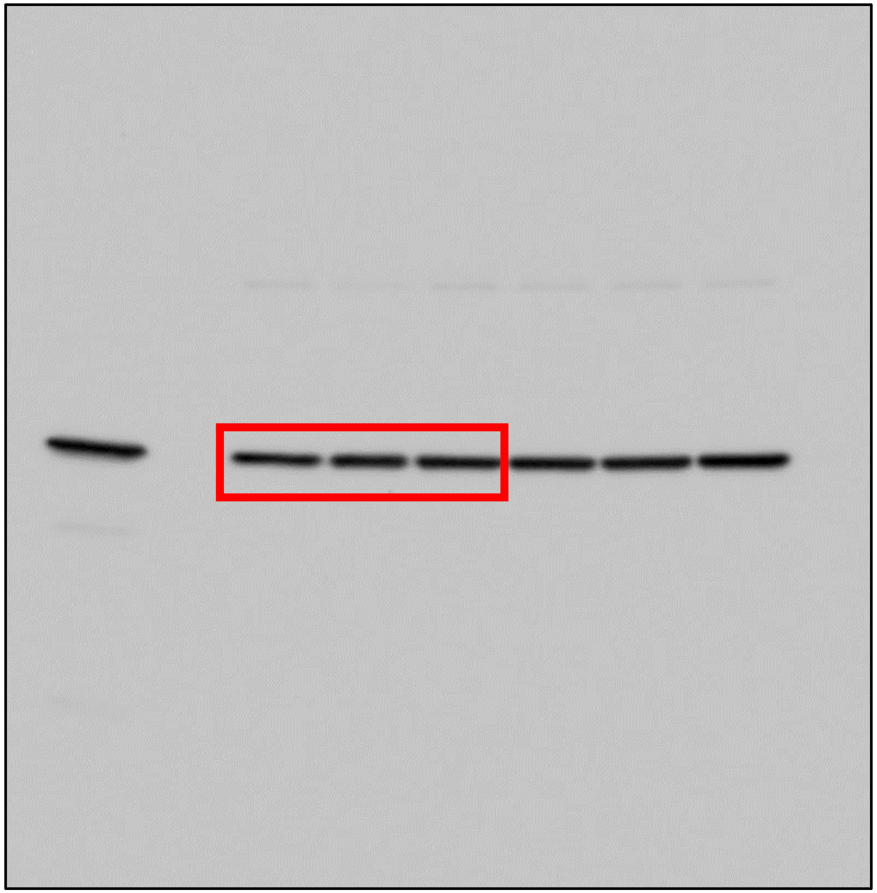

**Branscome et al. Supplementary Figure 11**

p24 (Fig. 2c)

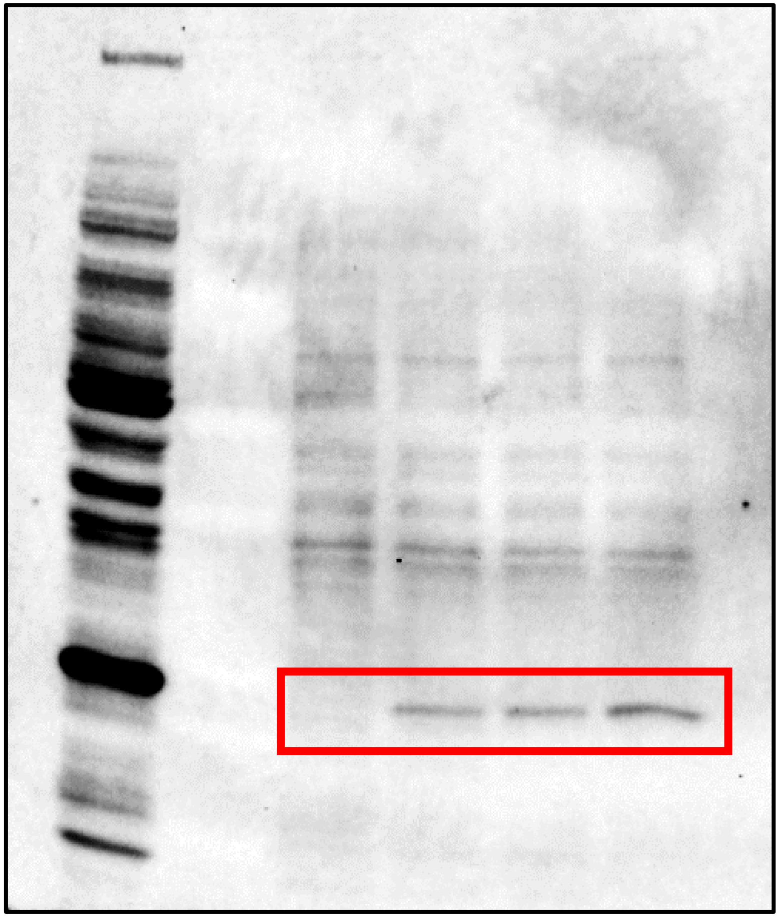

Actin (Fig. 2c)

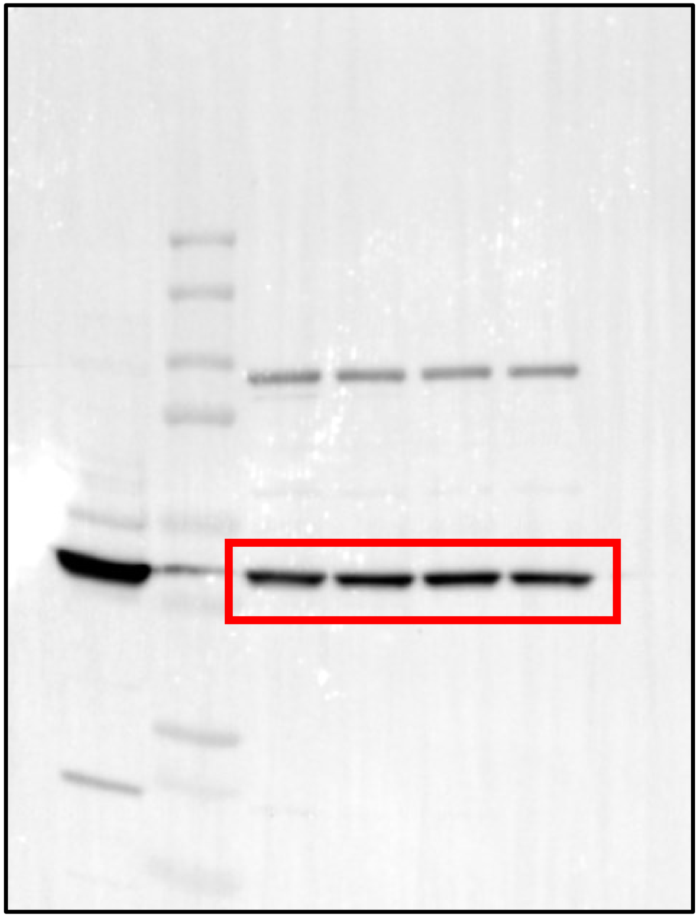

**Branscome et al. Supplementary Figure 11**

Pr55/p24 (Fig. 2e)

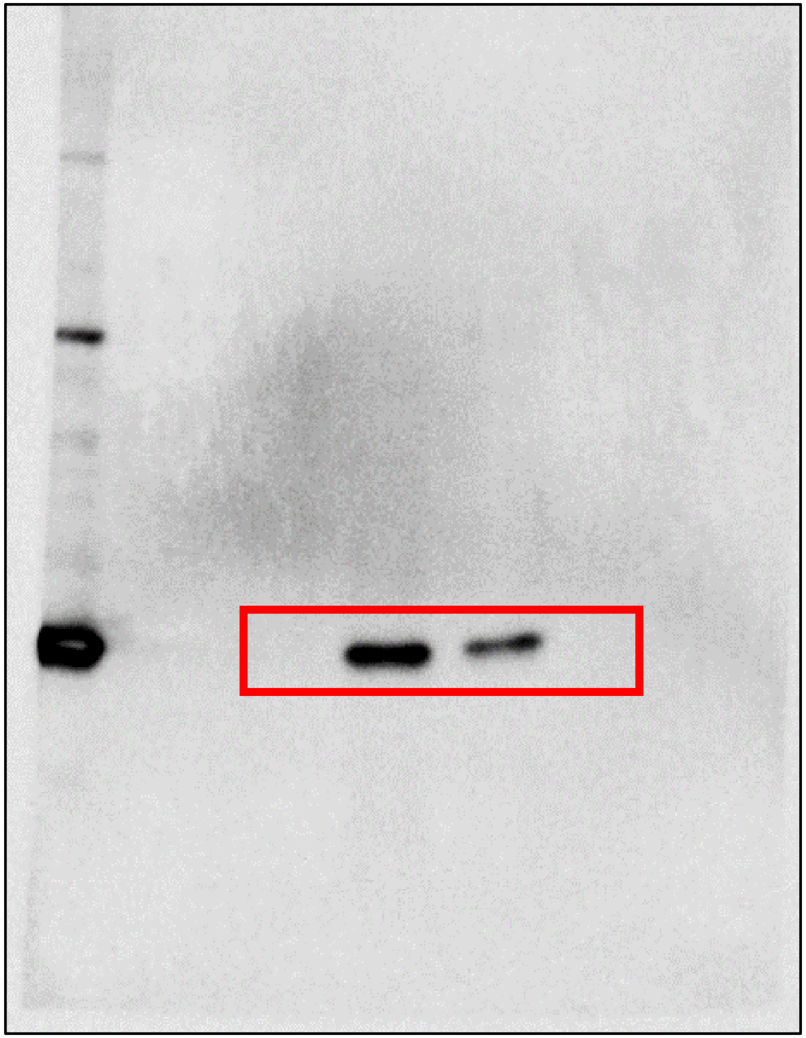

Nef (Fig. 2e)

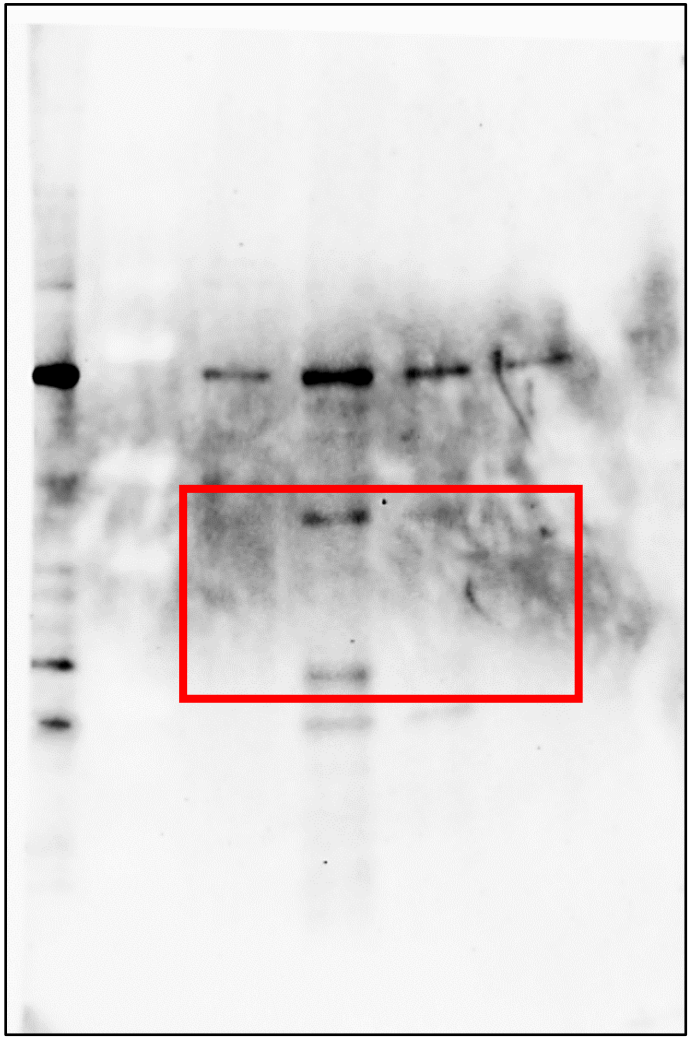

**Branscome et al. Supplementary Figure 11**

IBA-1 (Fig. 2e)

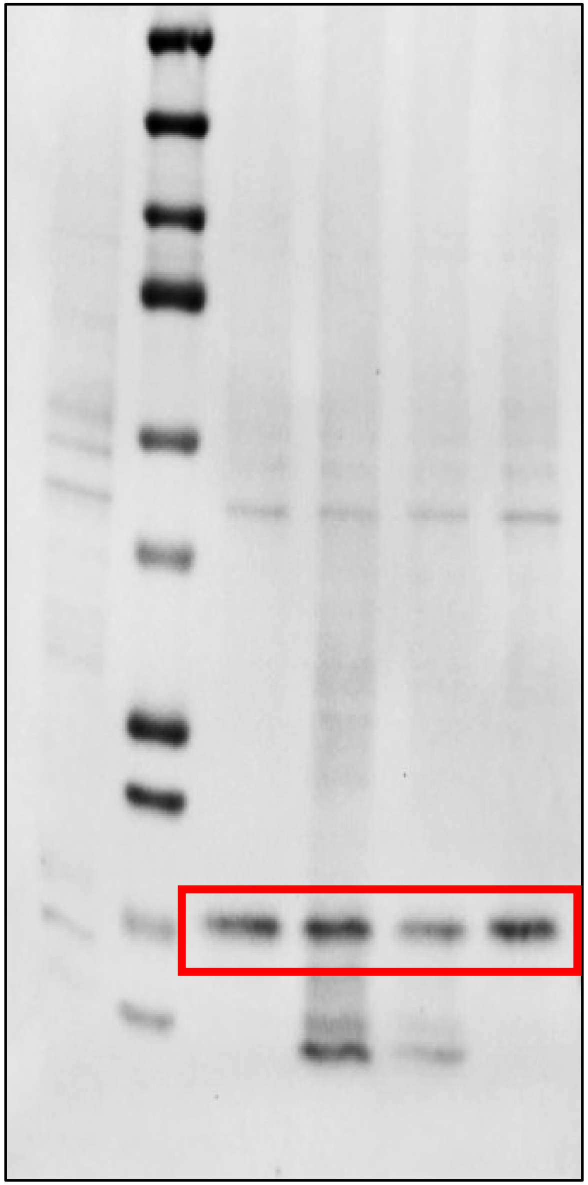

CD11b (Fig. 2e)

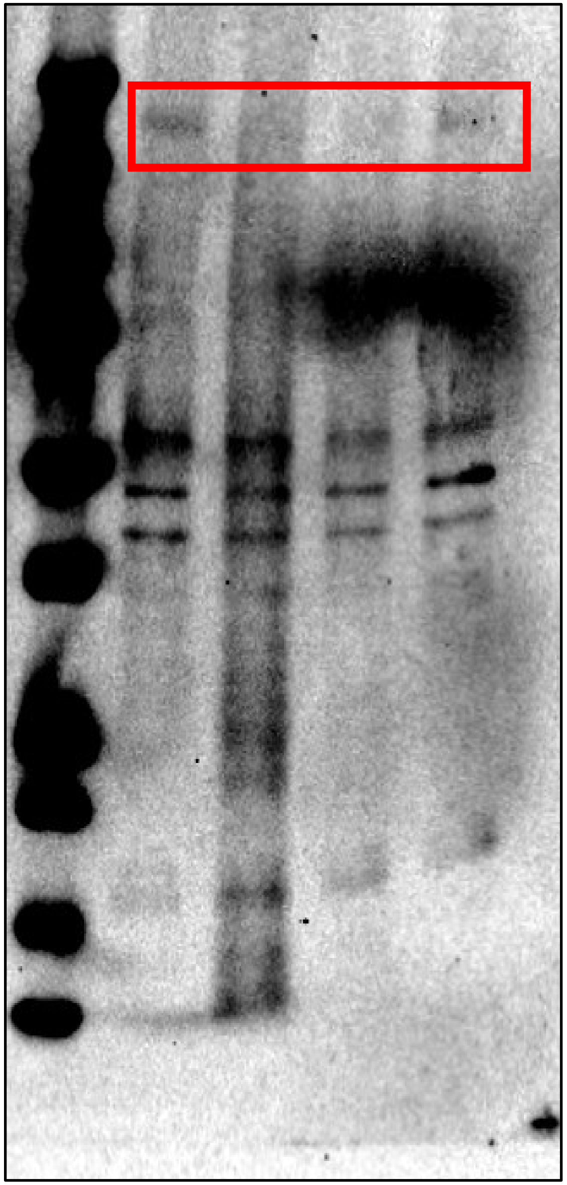

**Branscome et al. Supplementary Figure 11**

CD45 (Fig. 2e)

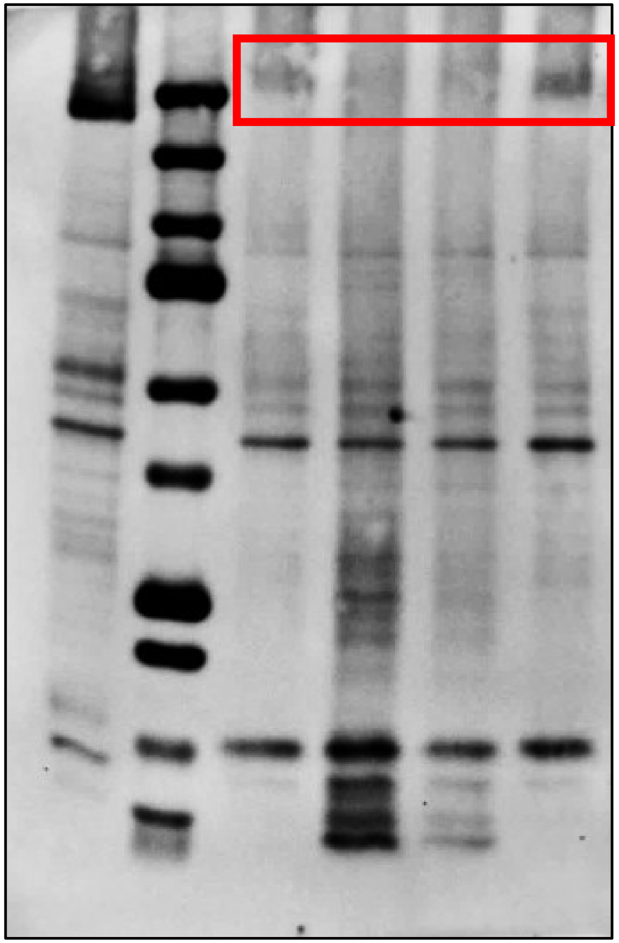

PARP-1 (Fig. 2e)

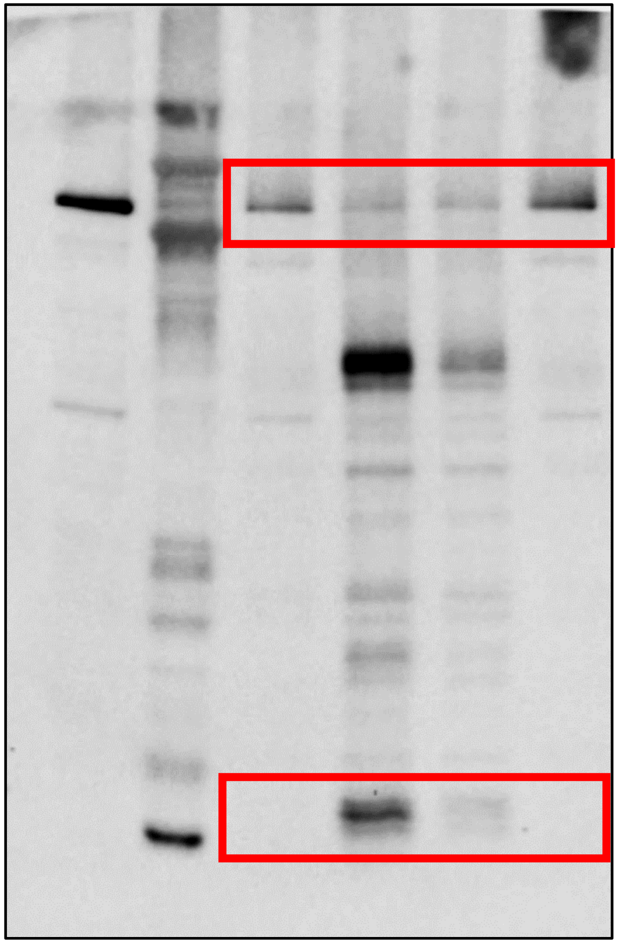

**Branscome et al. Supplementary Figure 11**

BAD (Fig. 2e)

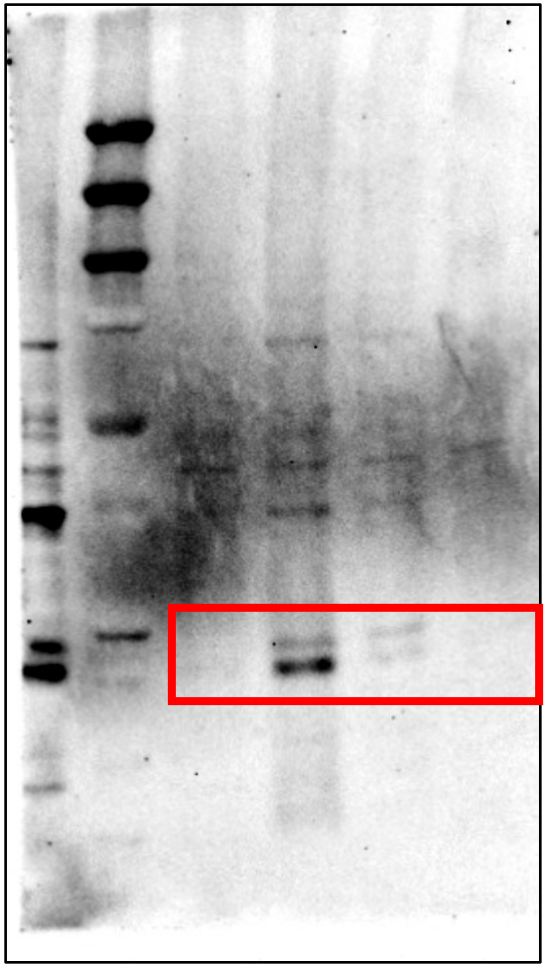

Actin (Fig. 2e)

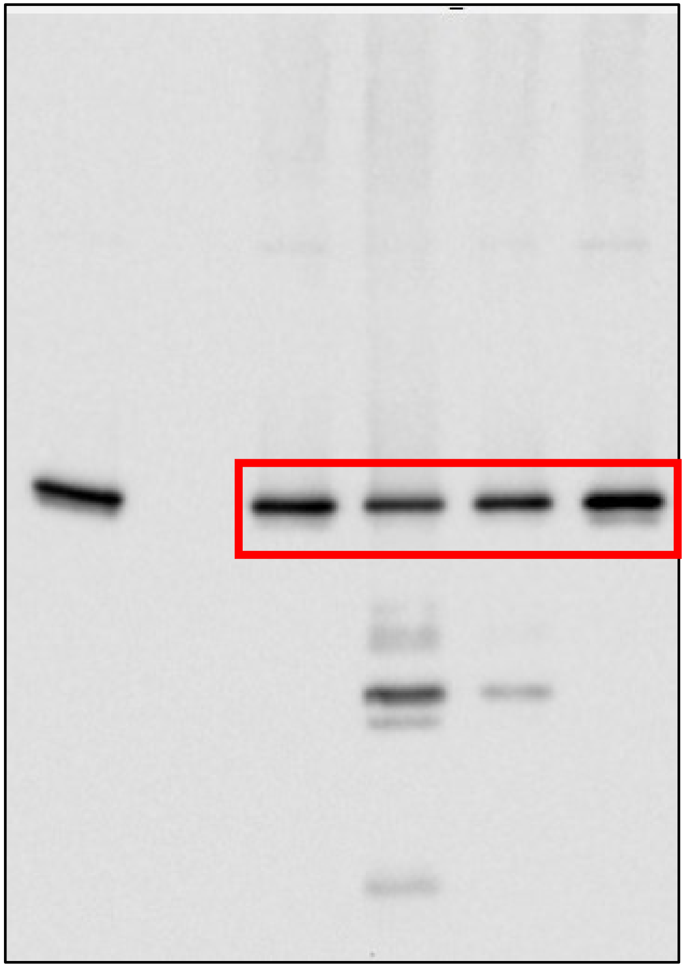

**Branscome et al. Supplementary Figure 11**

Glutamine Synthetase (Fig. 3a)

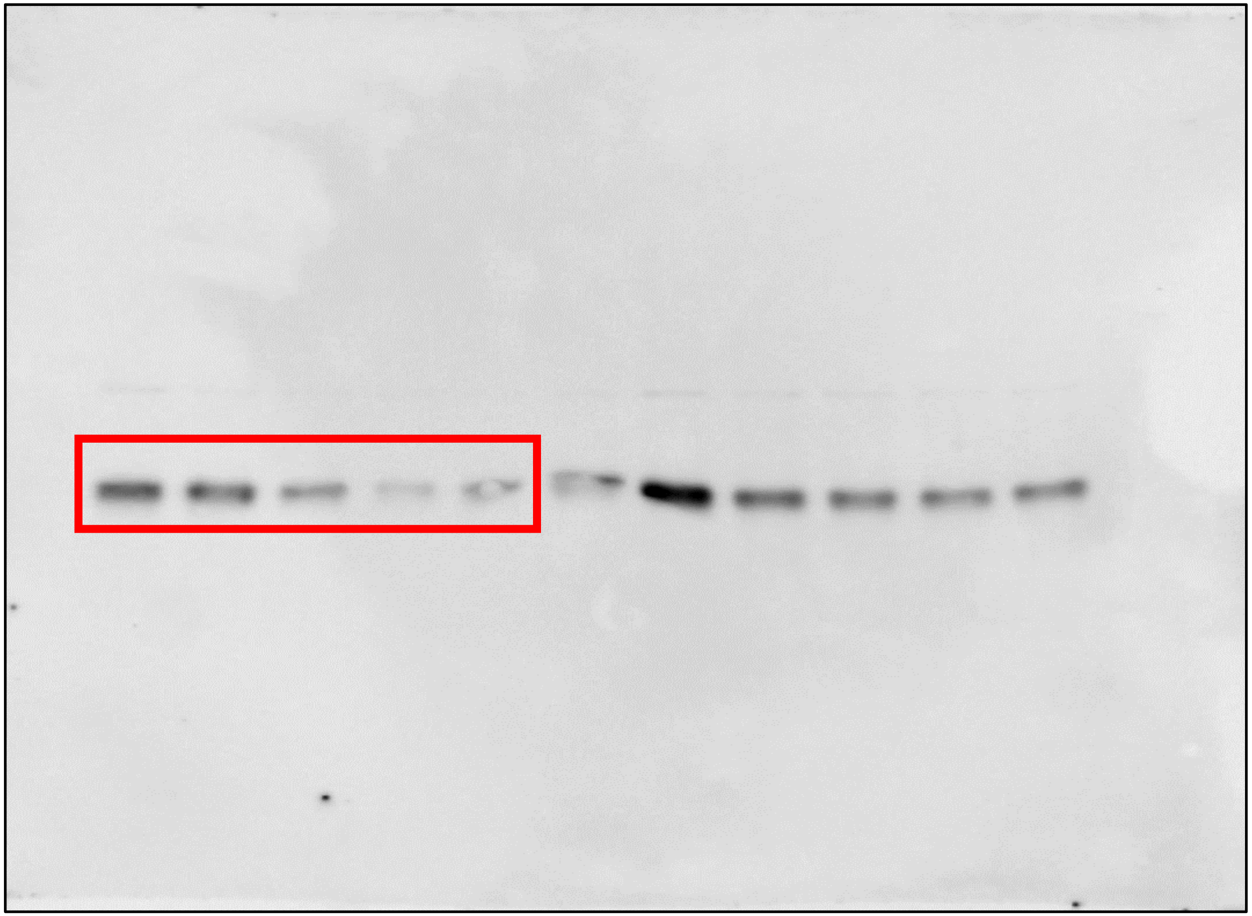

**Branscome et al. Supplementary Figure 11**

GFAP (Fig. 3a)

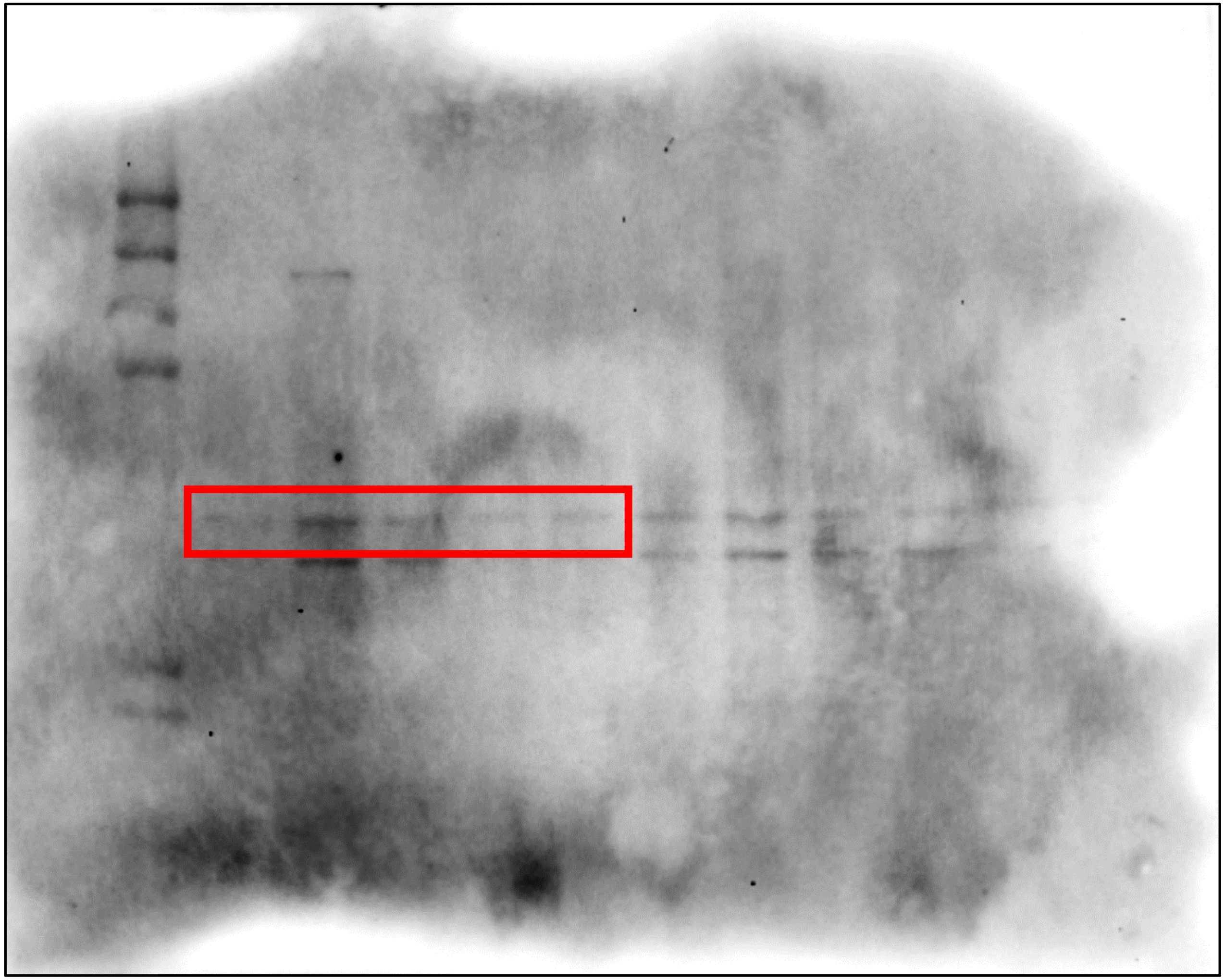

**Branscome et al. Supplementary Figure 11**

Tyrosine Hydroxylase (Fig. 3a)

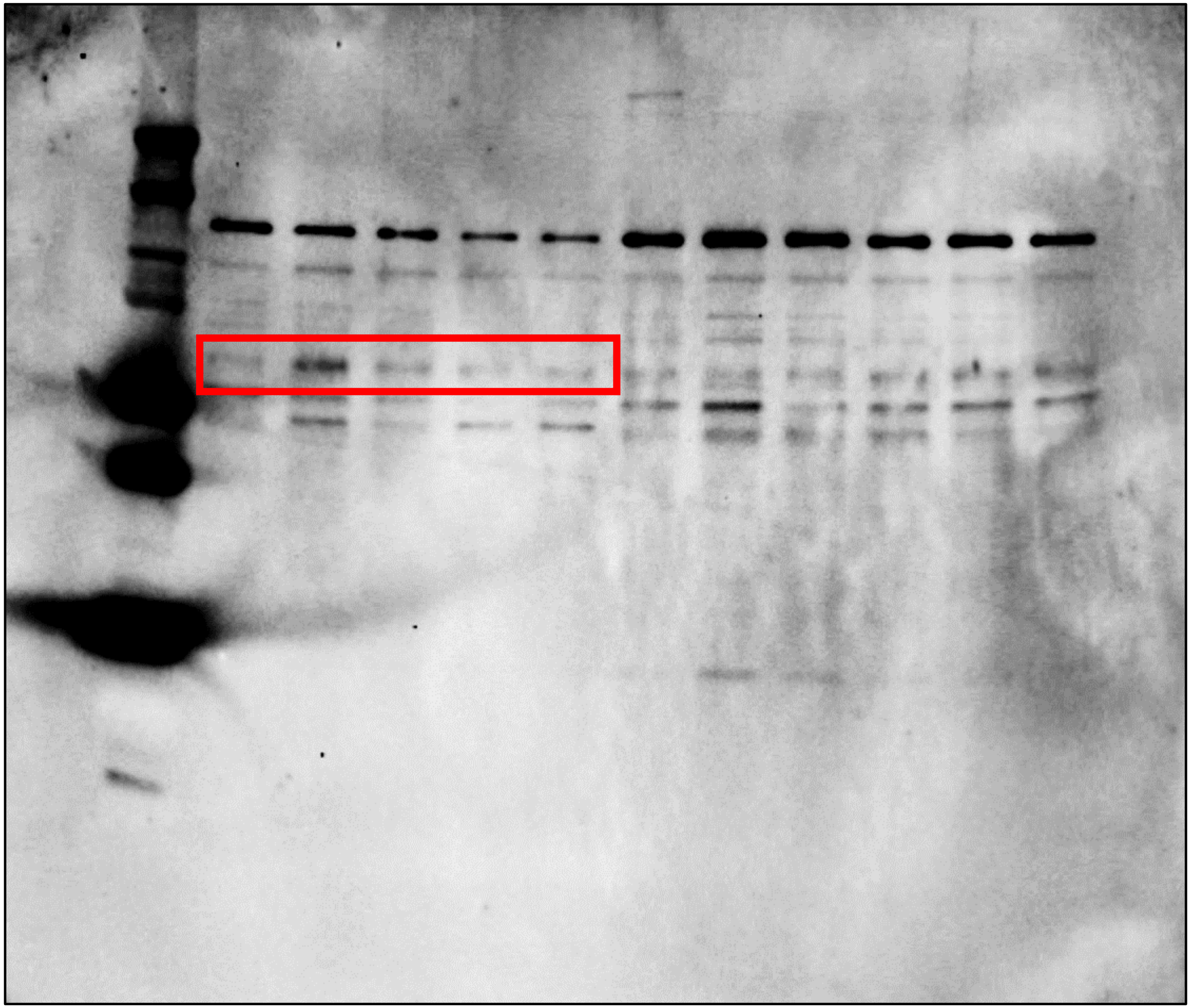

**Branscome et al. Supplementary Figure 11**

FOXA2 (Fig. 3a)

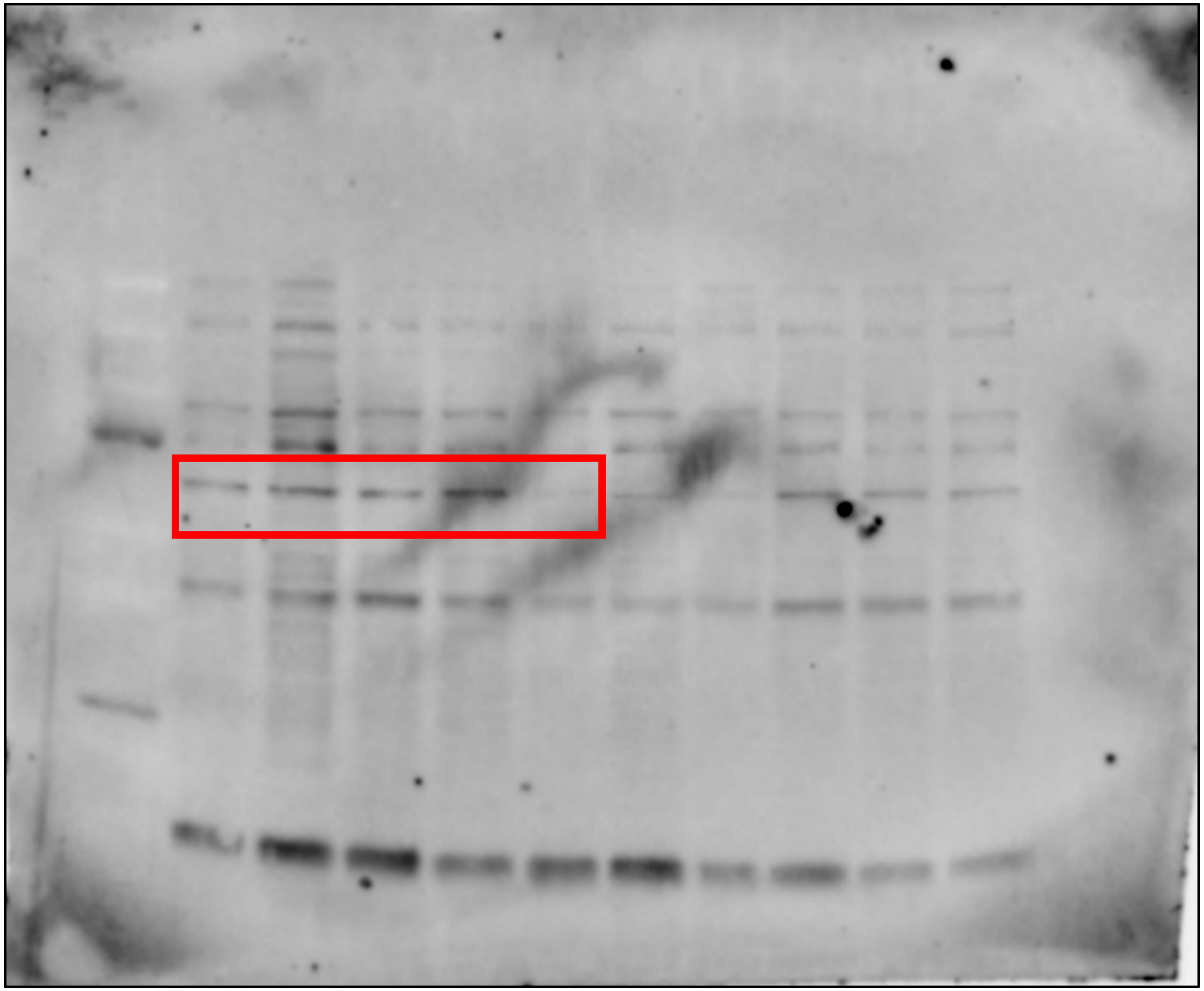

**Branscome et al. Supplementary Figure 11**

GAD65 (Fig. 3a)

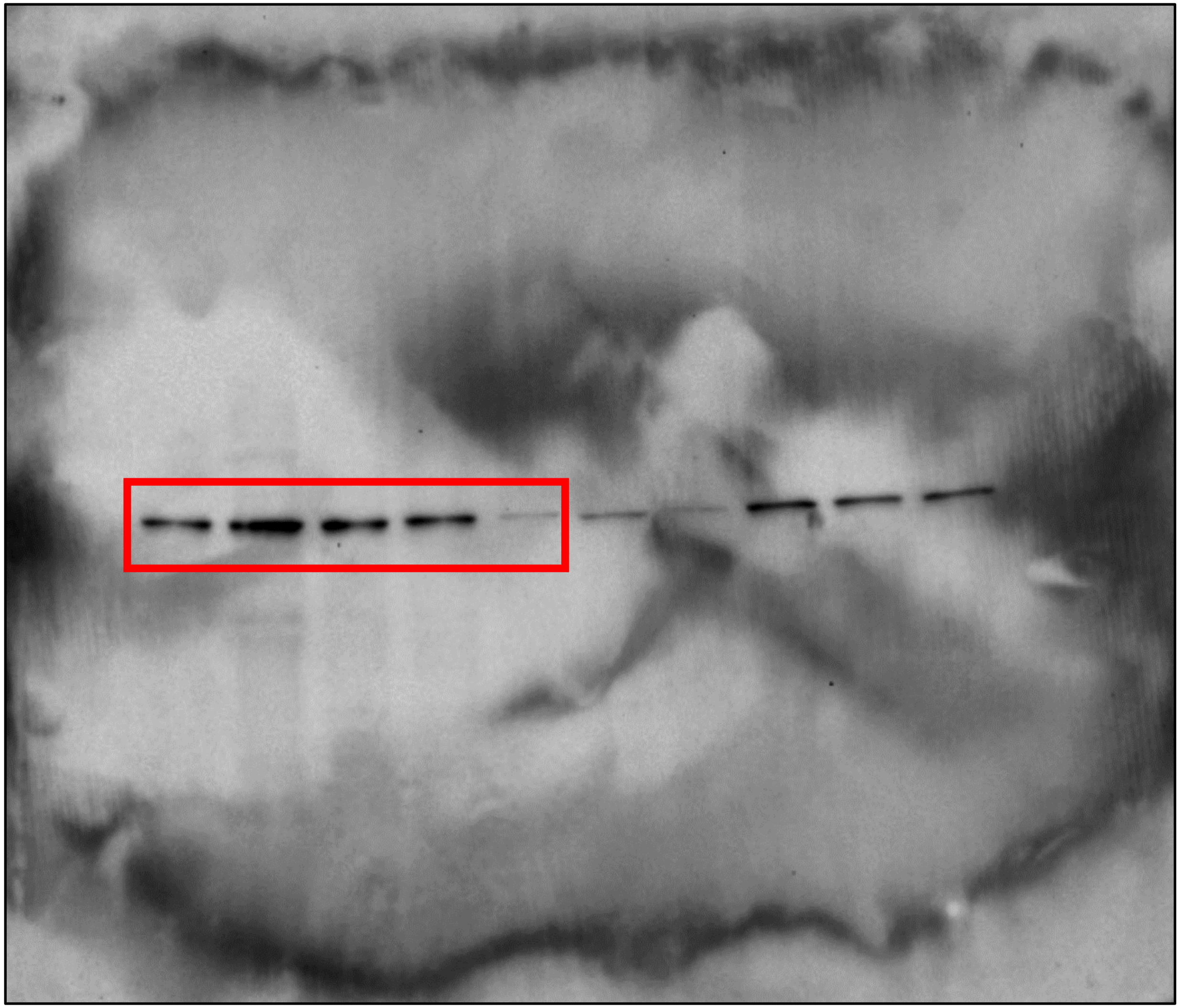

**Branscome et al. Supplementary Figure 11**

GAD67 (Fig. 3a)

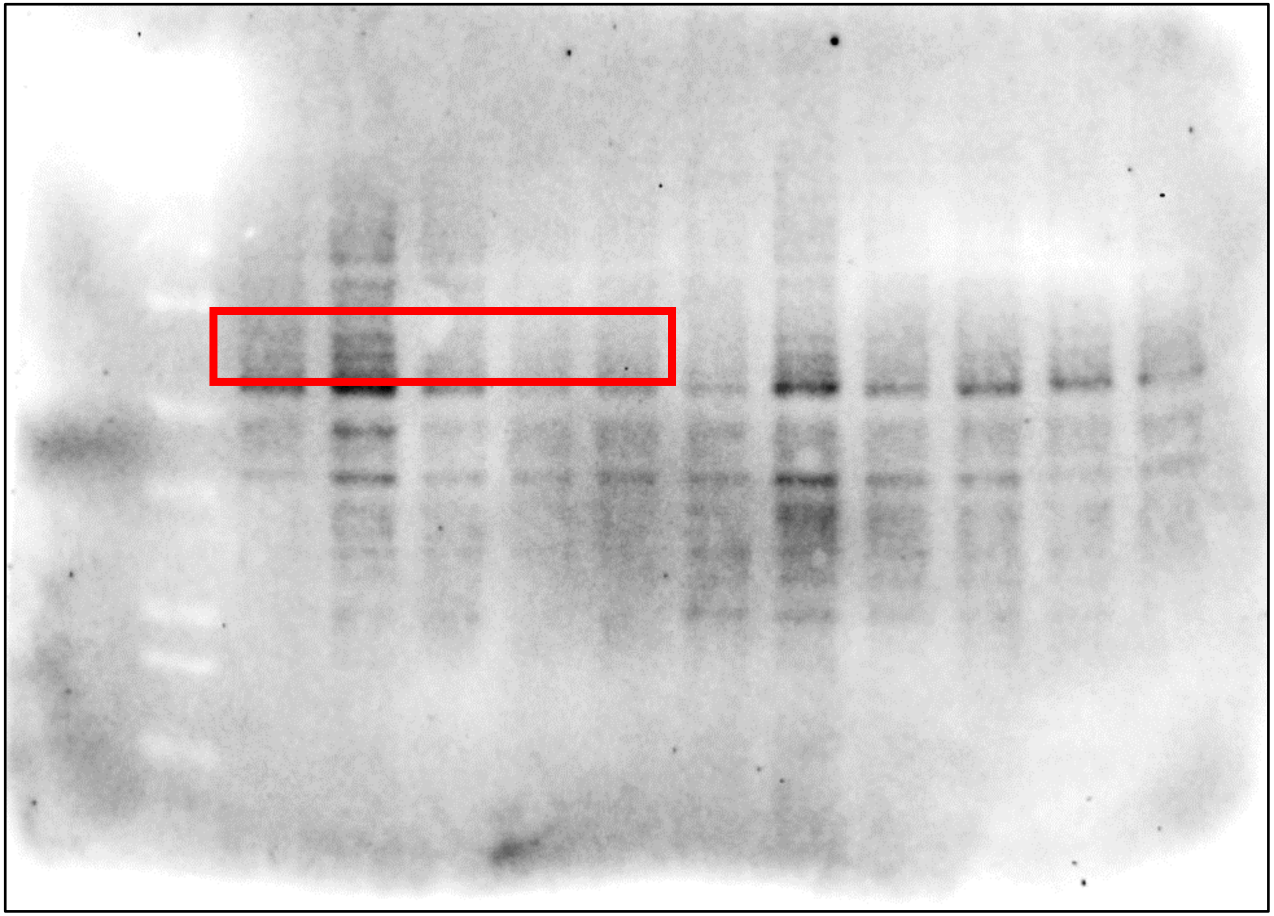

**Branscome et al. Supplementary Figure 11**

BNPI (Fig. 3a)

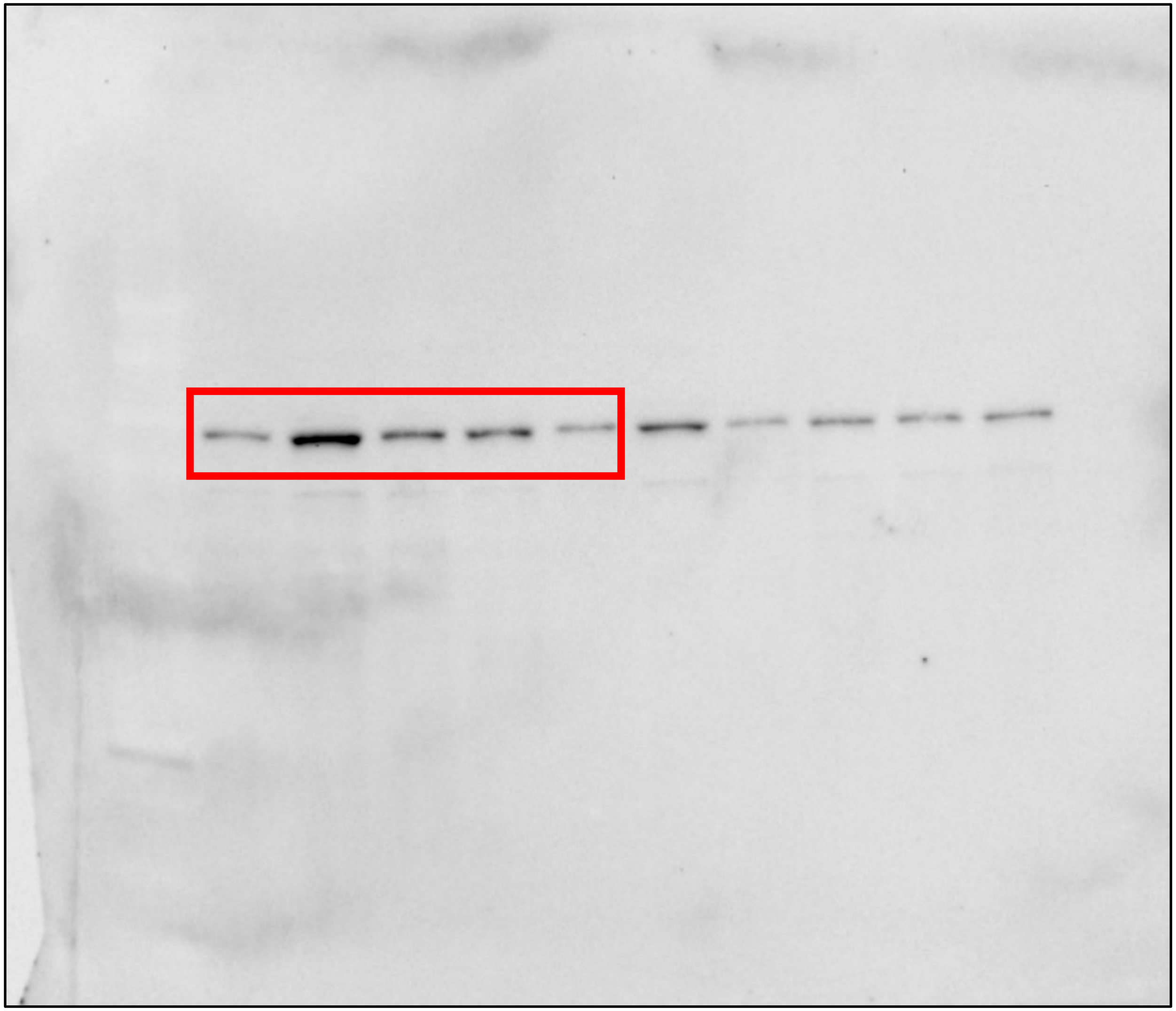

**Branscome et al. Supplementary Figure 11**

SOX2 (Fig. 3a)

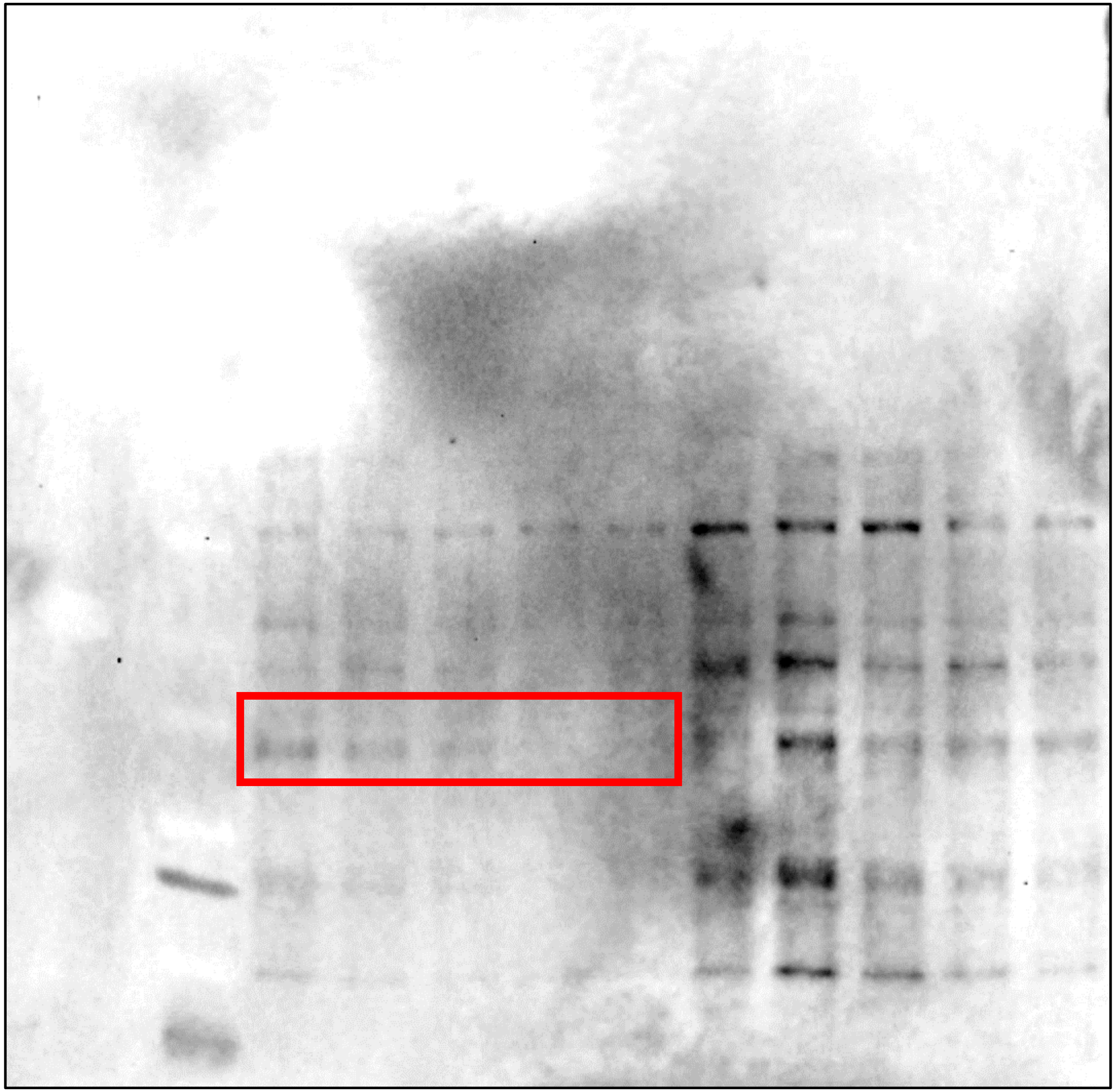

**Branscome et al. Supplementary Figure 11**

CD11b (Fig. 3a)

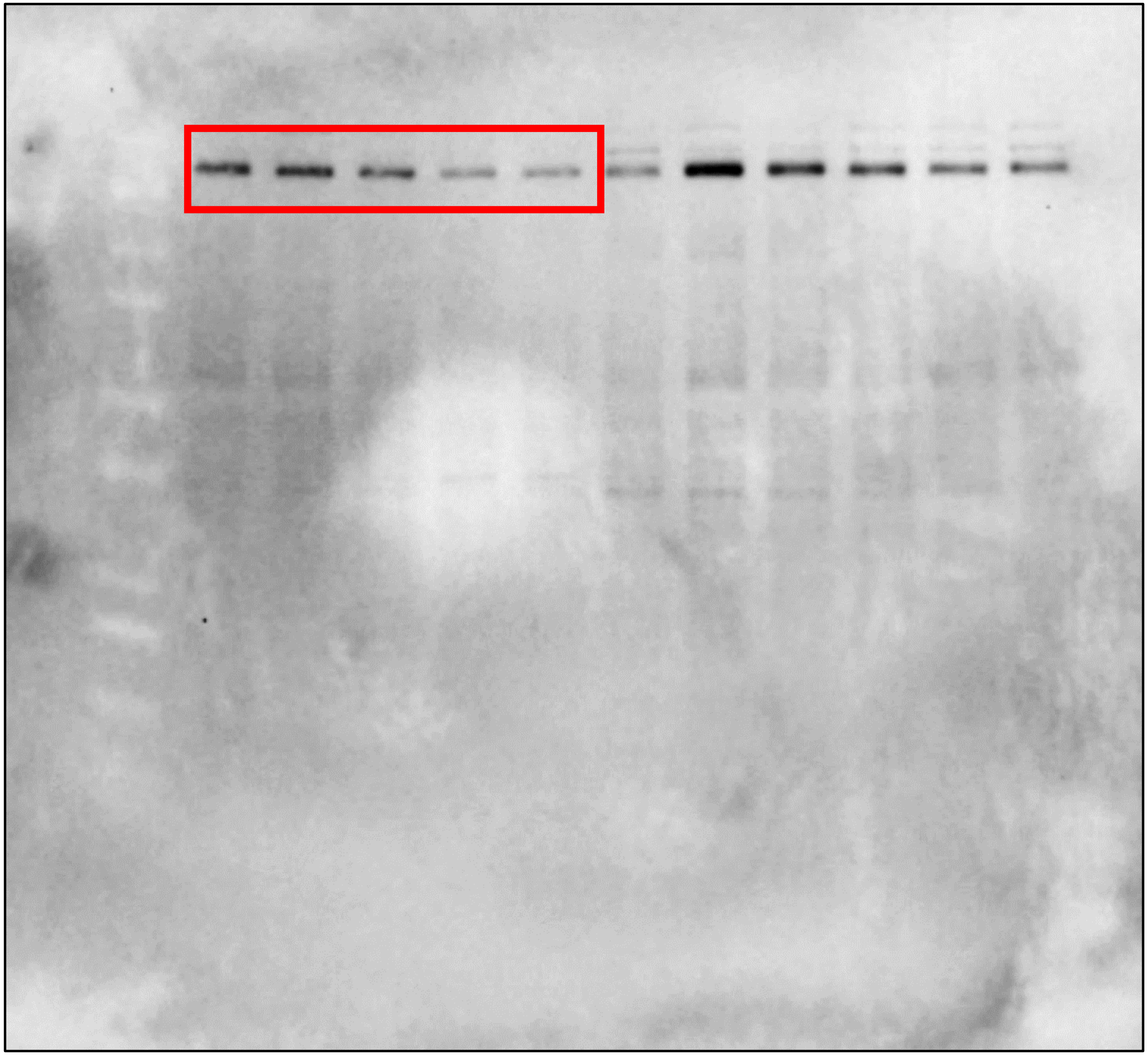

**Branscome et al. Supplementary Figure 11**

CD163 (Fig. 3a)

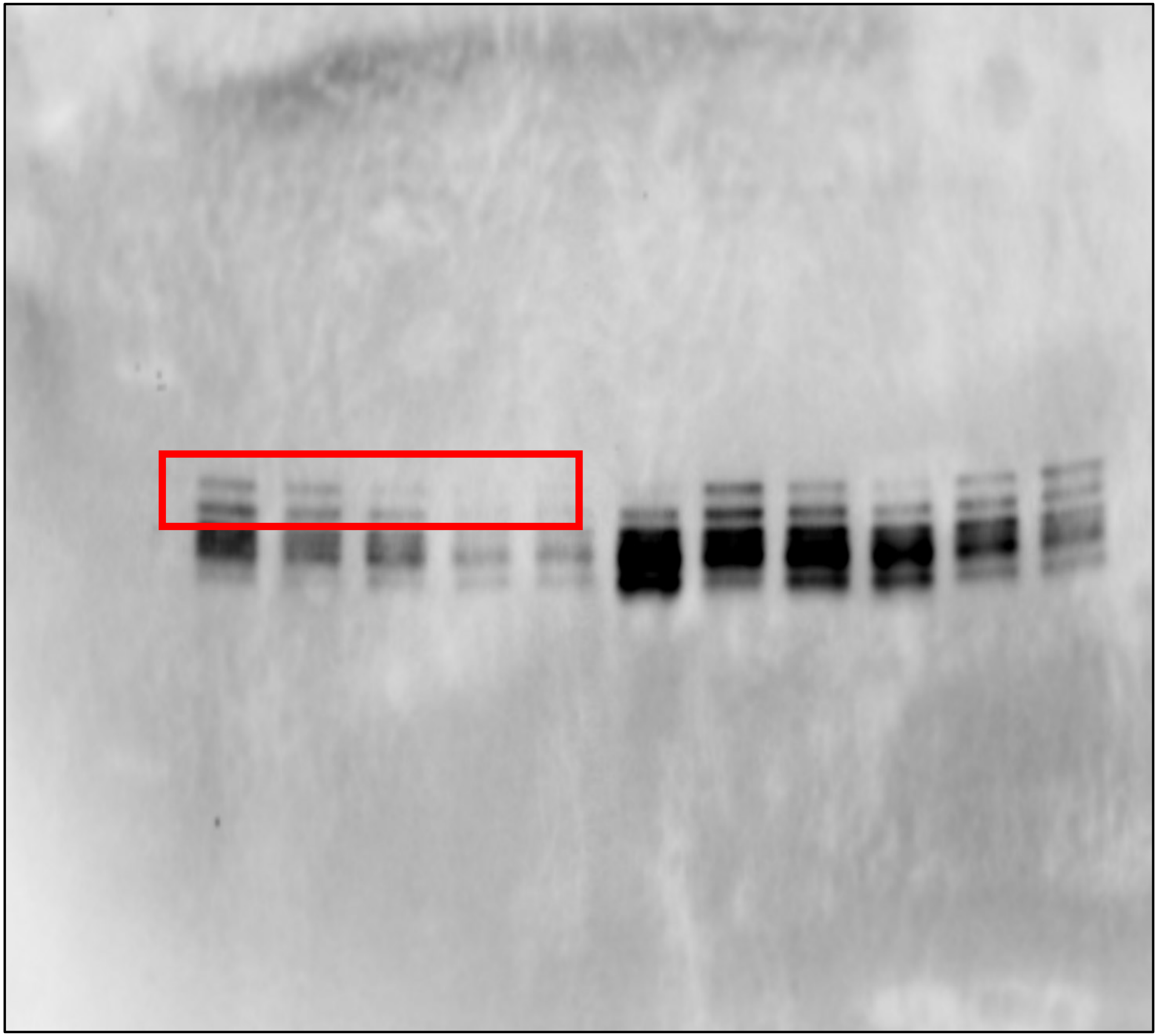

**Branscome et al. Supplementary Figure 11**

Actin (Fig. 3a)

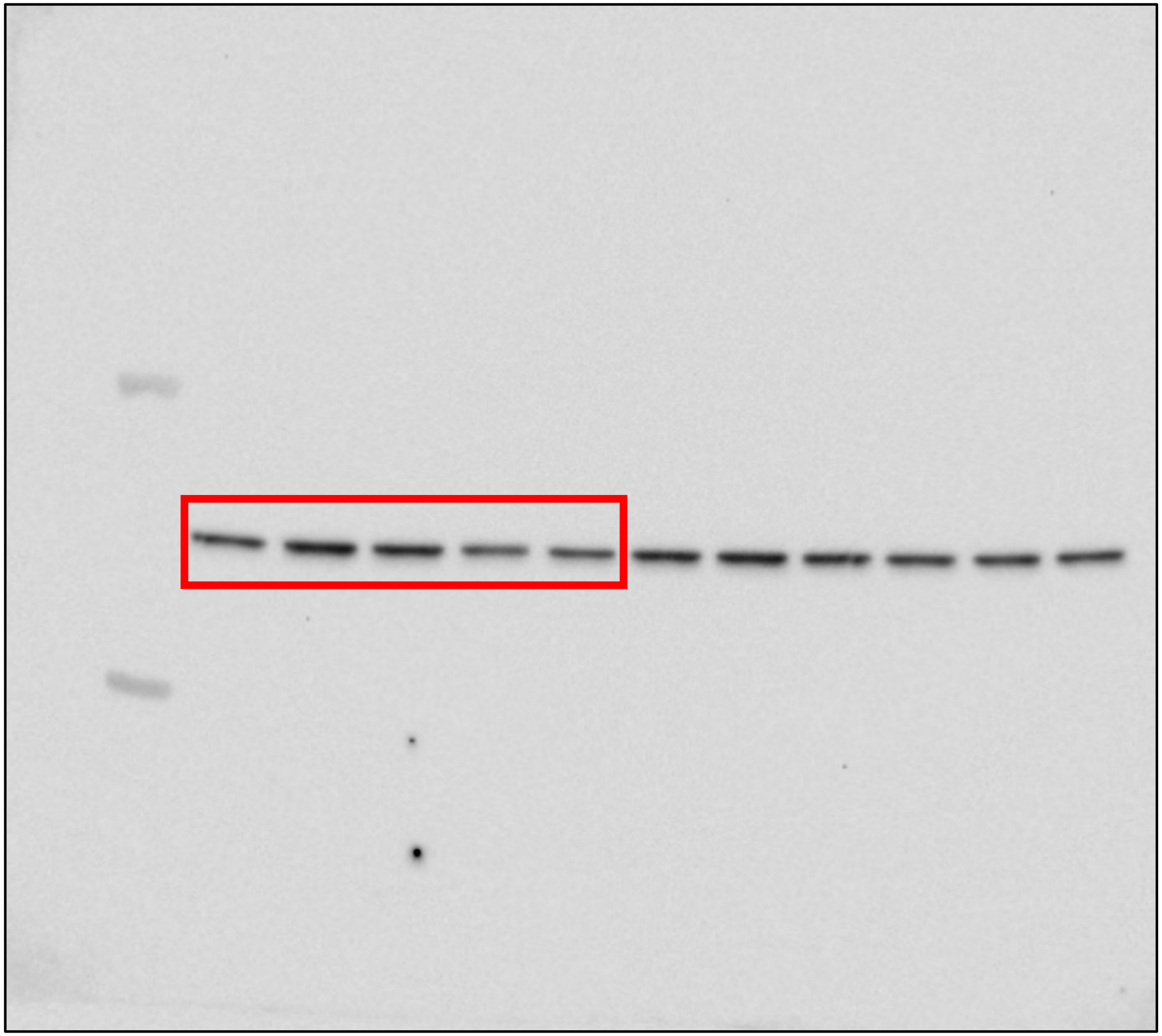

Branscome et al. Supplementary Figure 11

p19 (Fig. 3b)

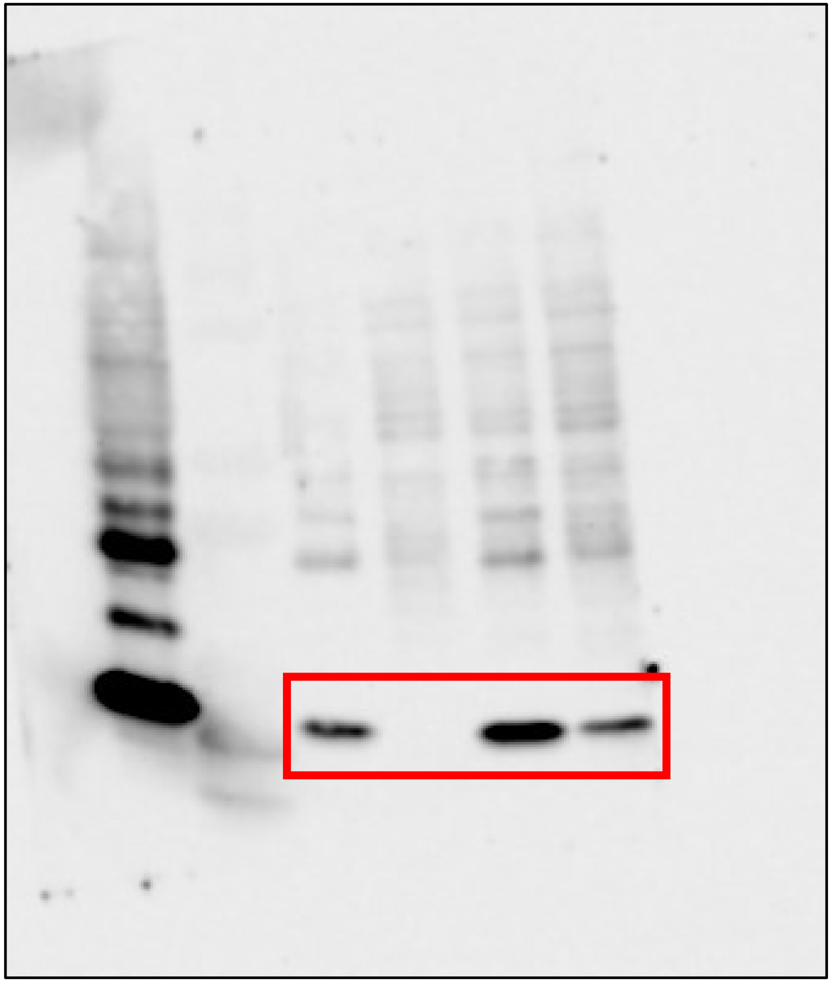

Tax (Fig. 3b)

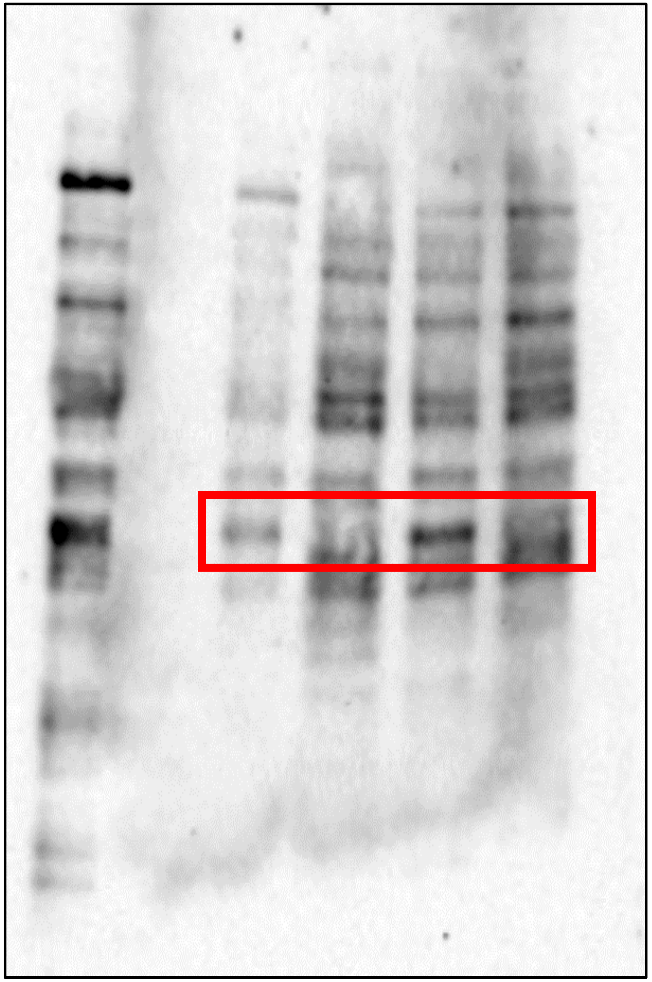

**Branscome et al. Supplementary Figure 11**

Actin (Fig. 3b)

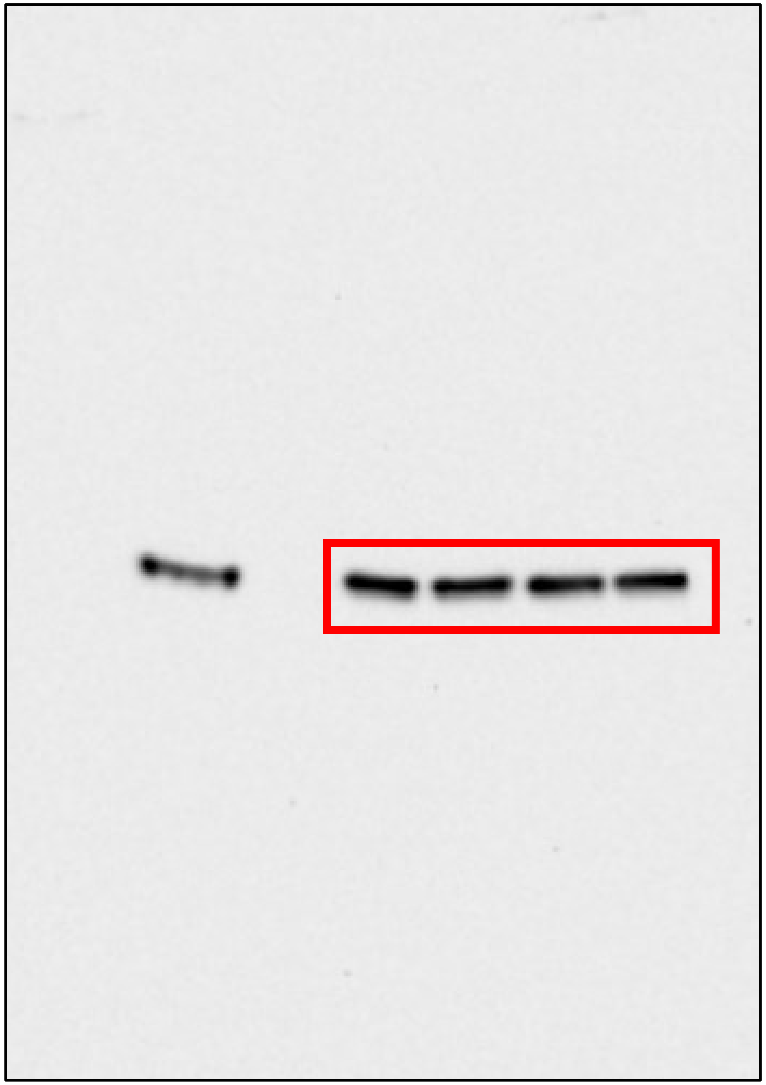

**Branscome et al. Supplementary Figure 11**

PARP-1 (Fig. 5a)

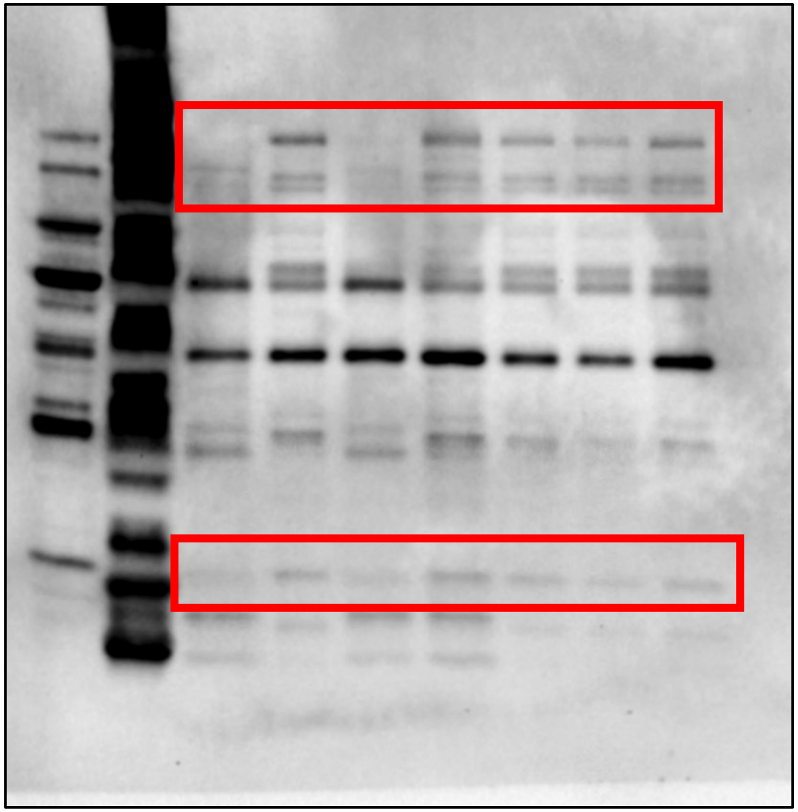

Caspase-3 (Fig. 5a)

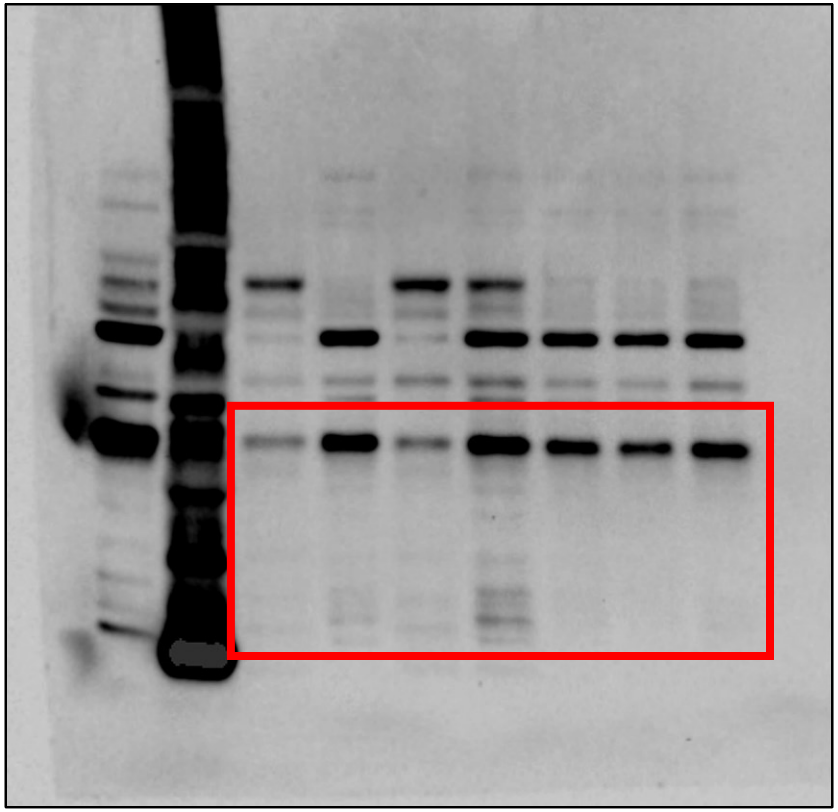

**Branscome et al. Supplementary Figure 11**

BAD (Fig. 5a)

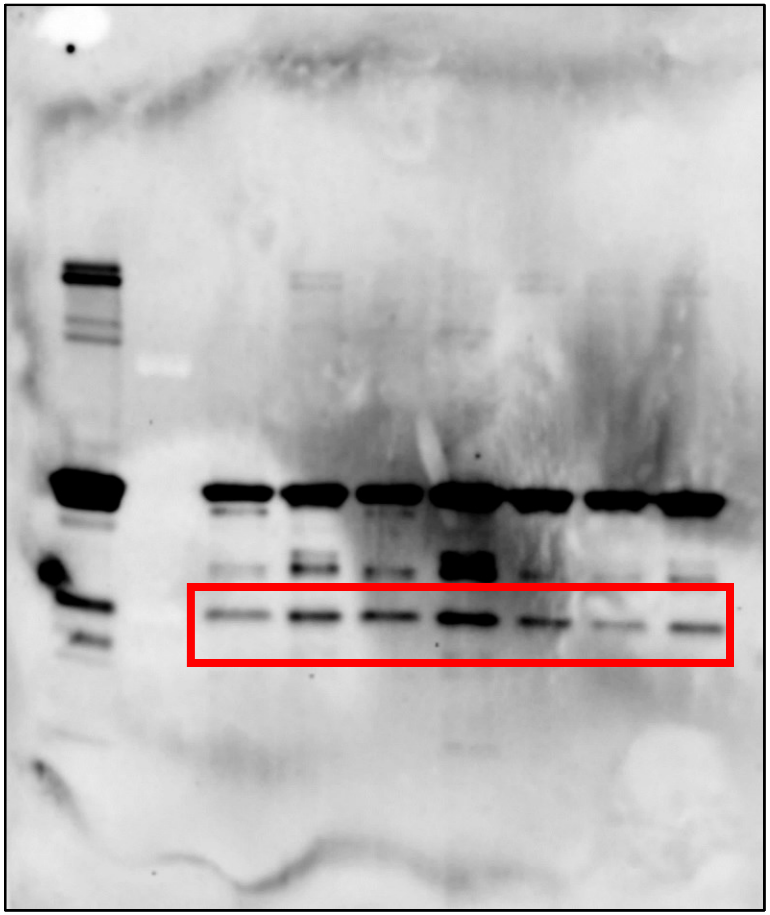

Actin (Fig. 5a)

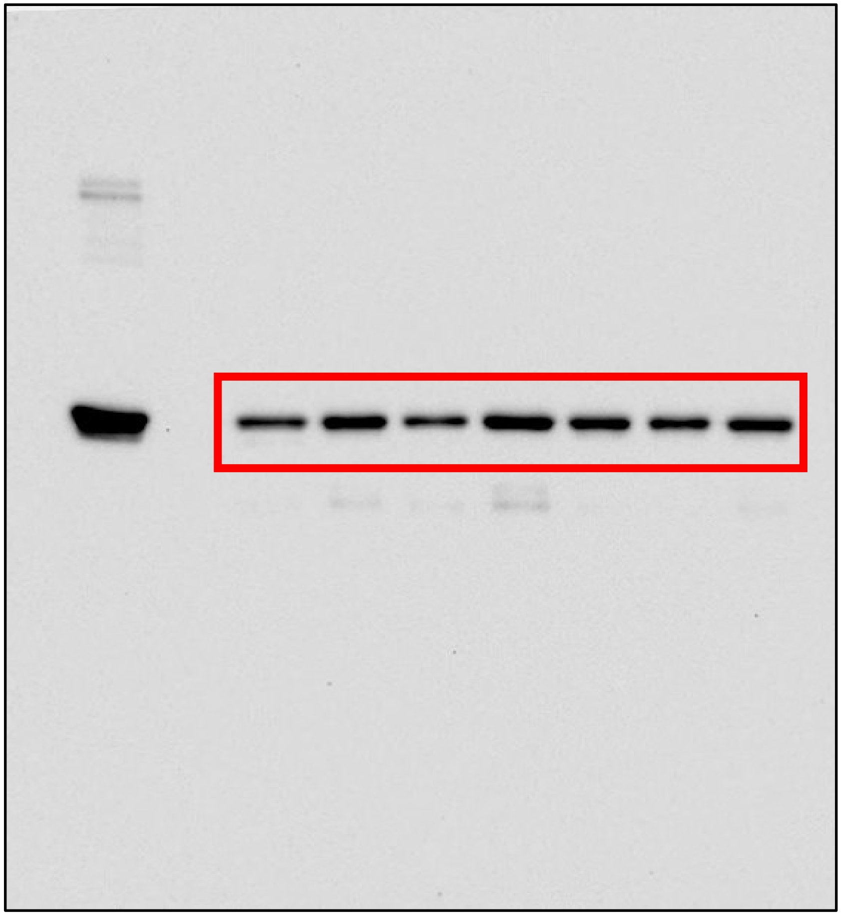

**Branscome et al. Supplementary Figure 11**

TNFα (Fig. 5b)

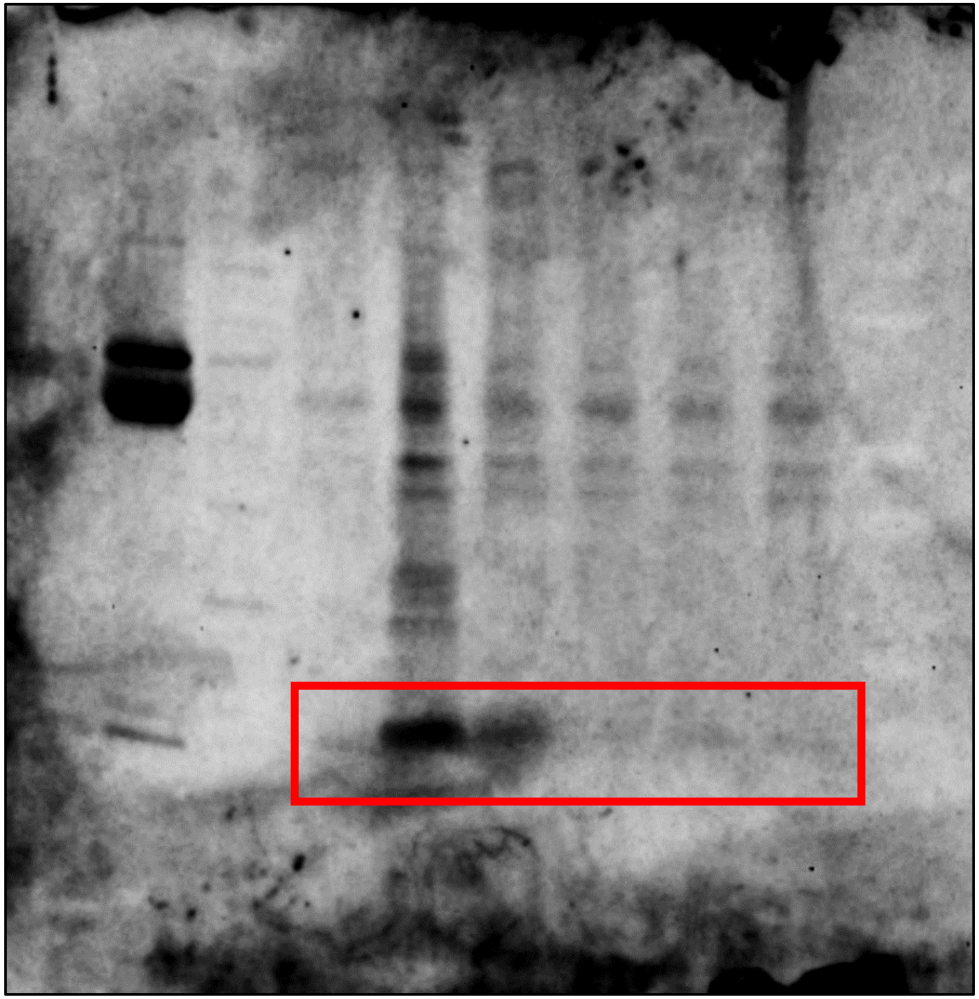

IL-8 (Fig. 5b)

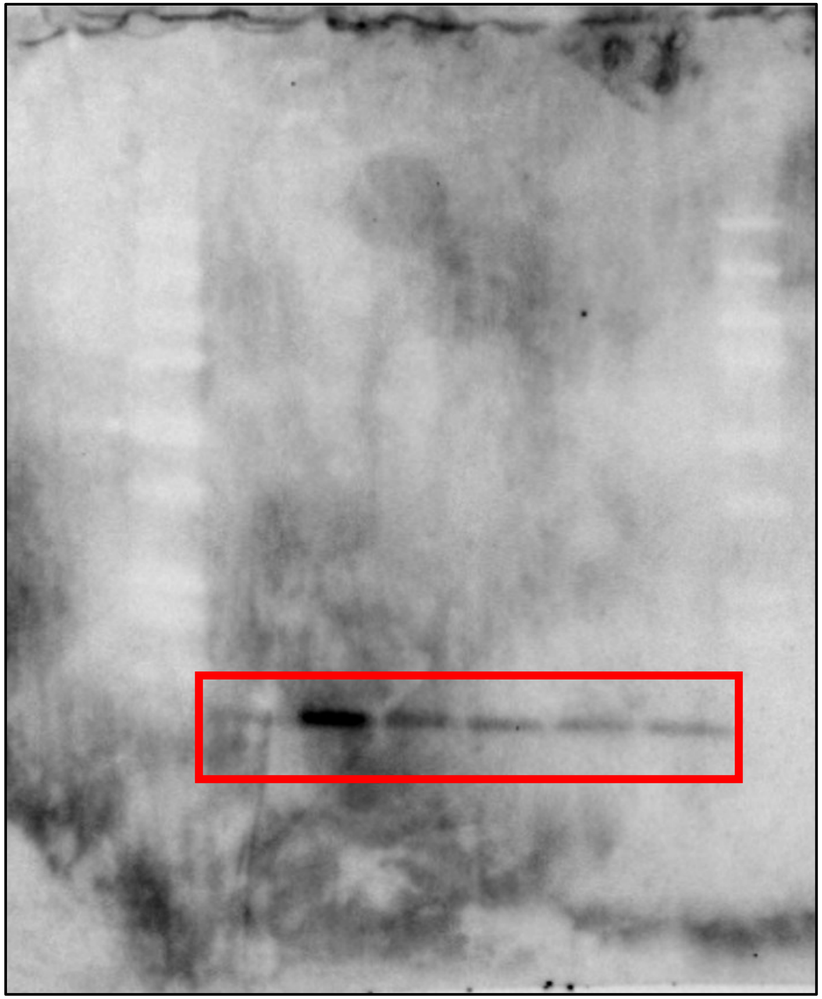

**Branscome et al. Supplementary Figure 11**

IL-1β (Fig. 5b)

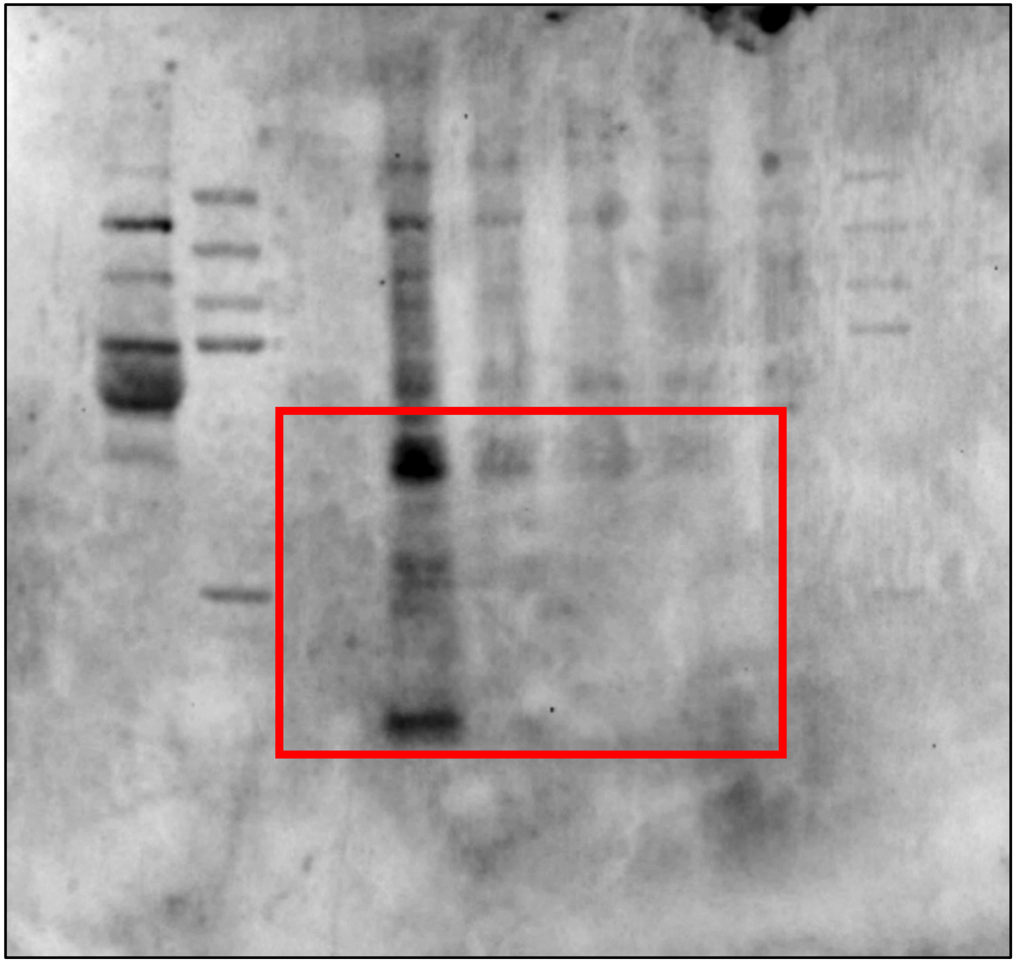

**Branscome et al. Supplementary Figure 11**

PKR  (Fig. 7c)

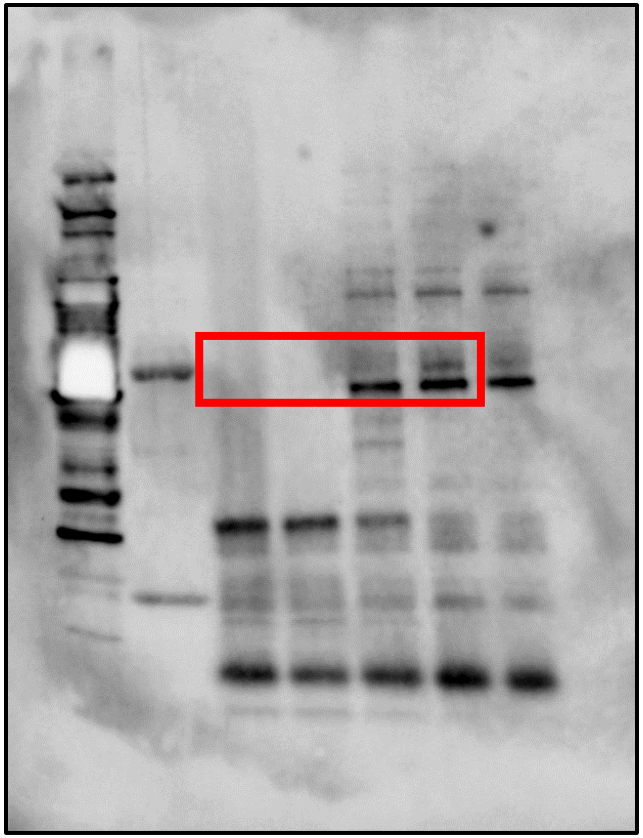

DICER (Fig. 7c)

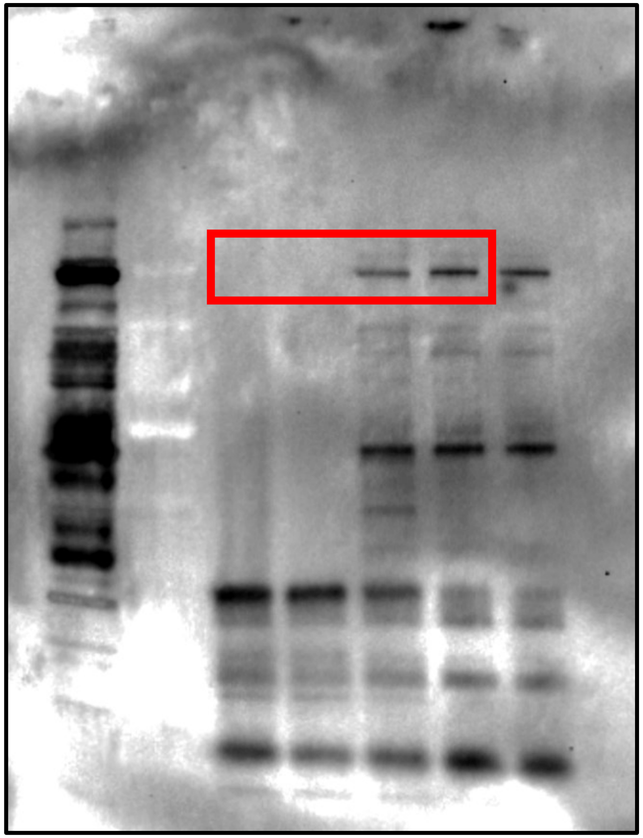

**Branscome et al. Supplementary Figure 11**

ADAR1  (Fig. 7c)

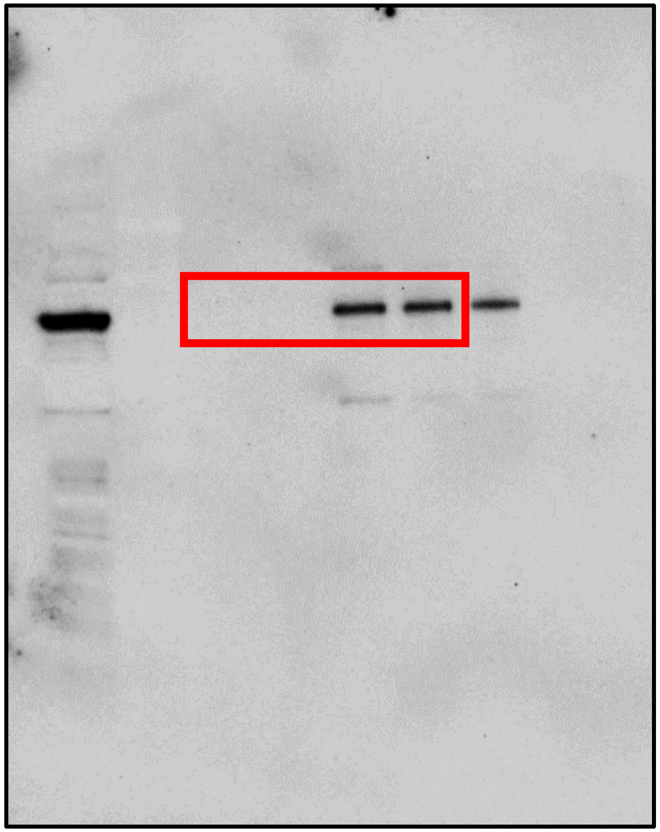

Branscome et al. Supplementary Figure 11

MMP9  (Supplementary Fig. 2d)

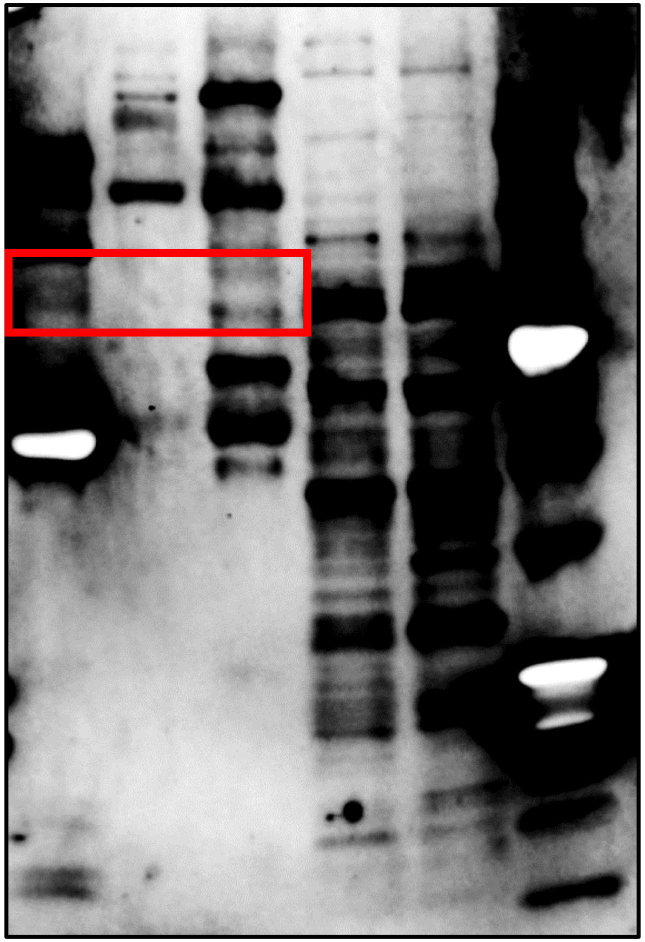

Actin (Supplementary Fig. 2d)

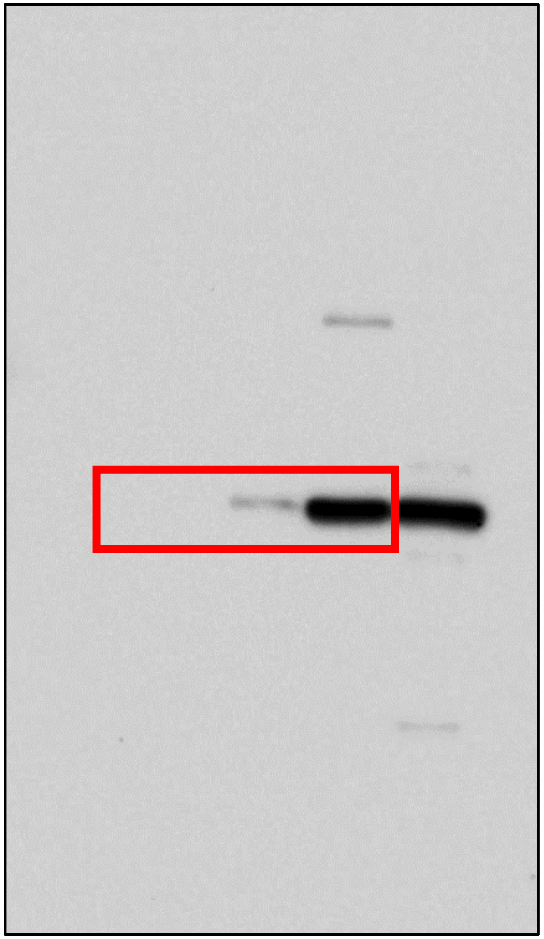

**Branscome et al. Supplementary Figure 11**

Supplementary Fig. 2e

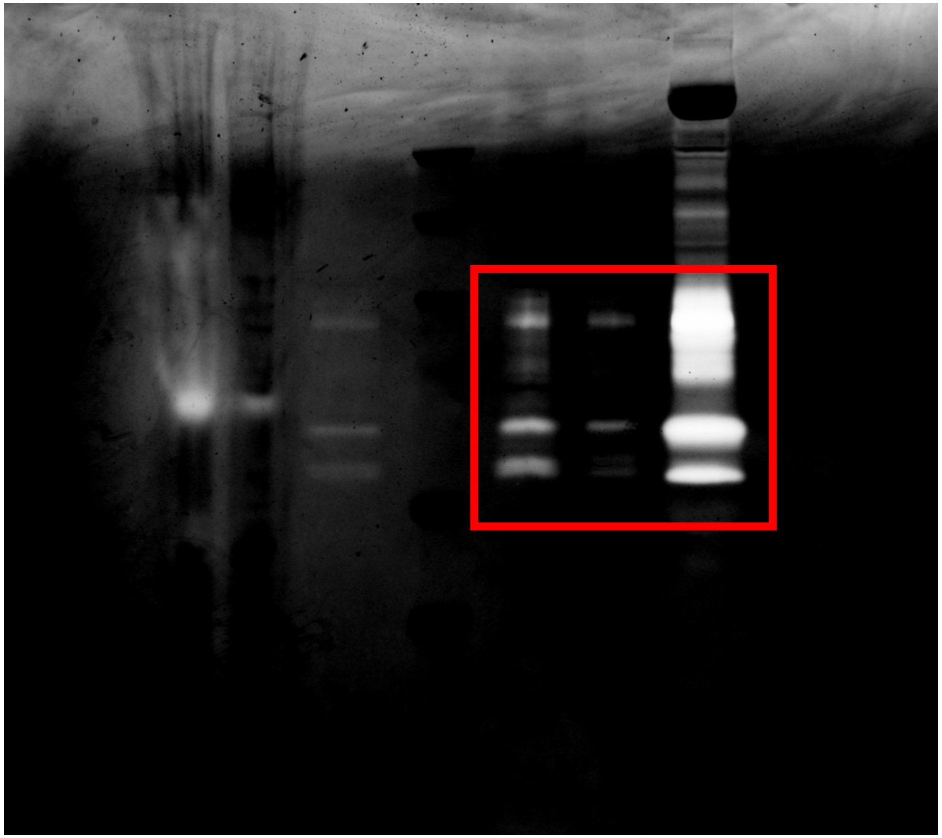

**Branscome et al. Supplementary Figure 11**

TNFα (Supplementary Fig. 3)

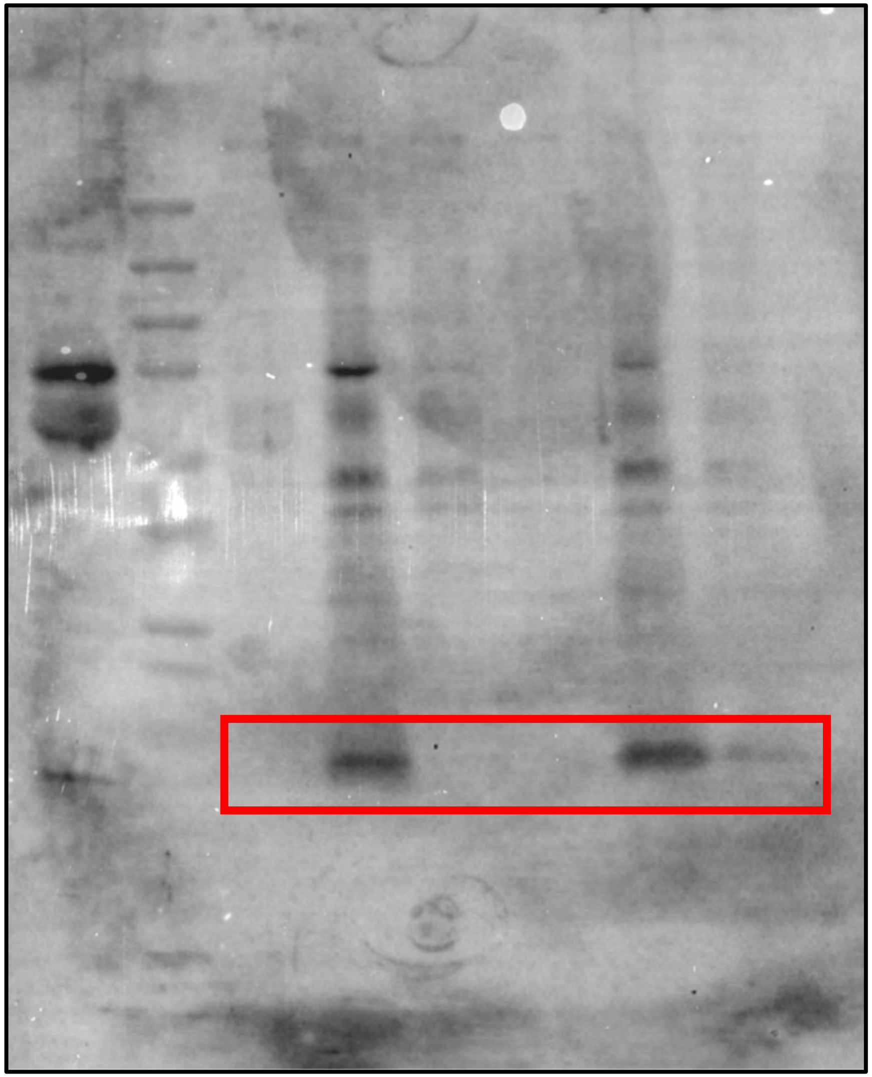

IL-8 (Supplementary Fig. 3)

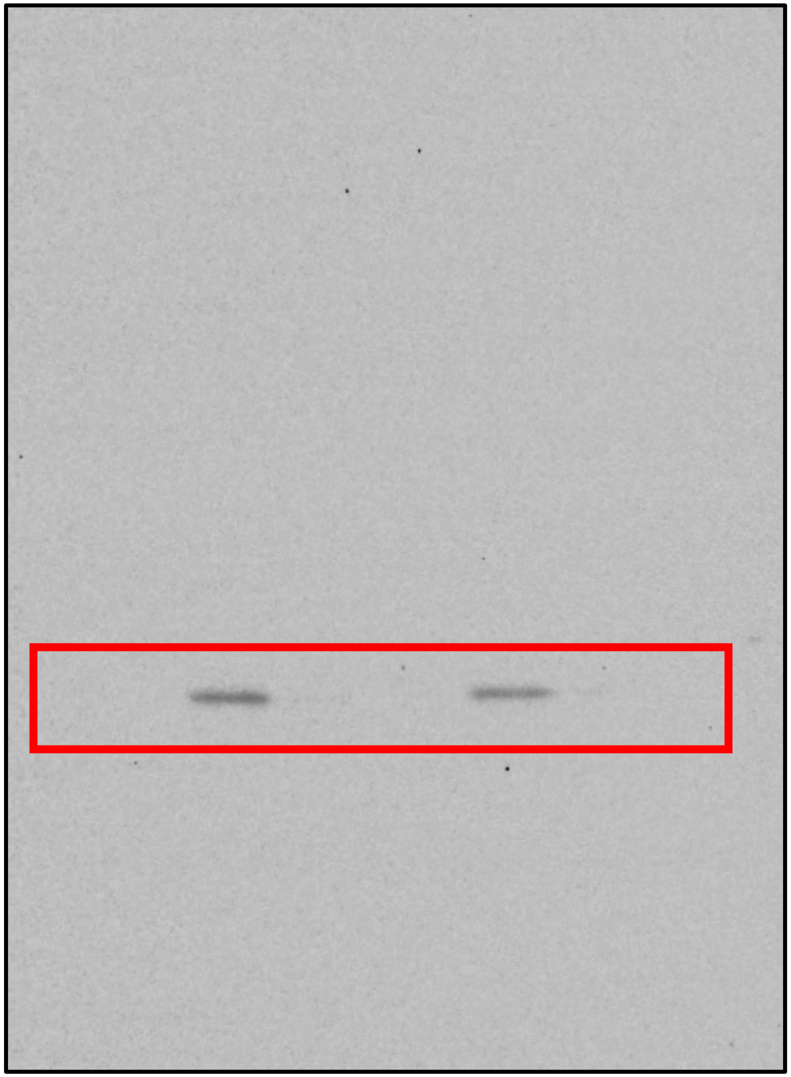

**Branscome et al. Supplementary Figure 11**

IL-1β (Supplementary Fig. 3)

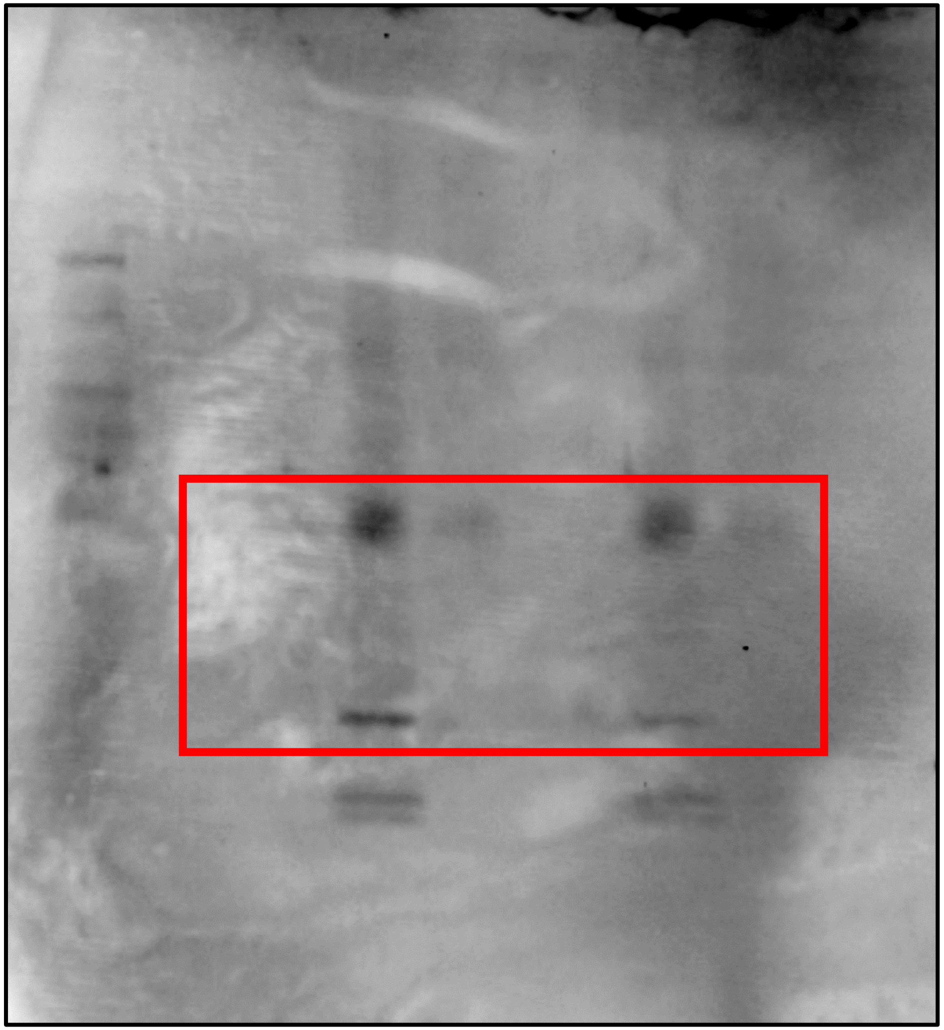

**Branscome et al. Supplementary Figure 11**

PKR (Supplementary Fig. 6)

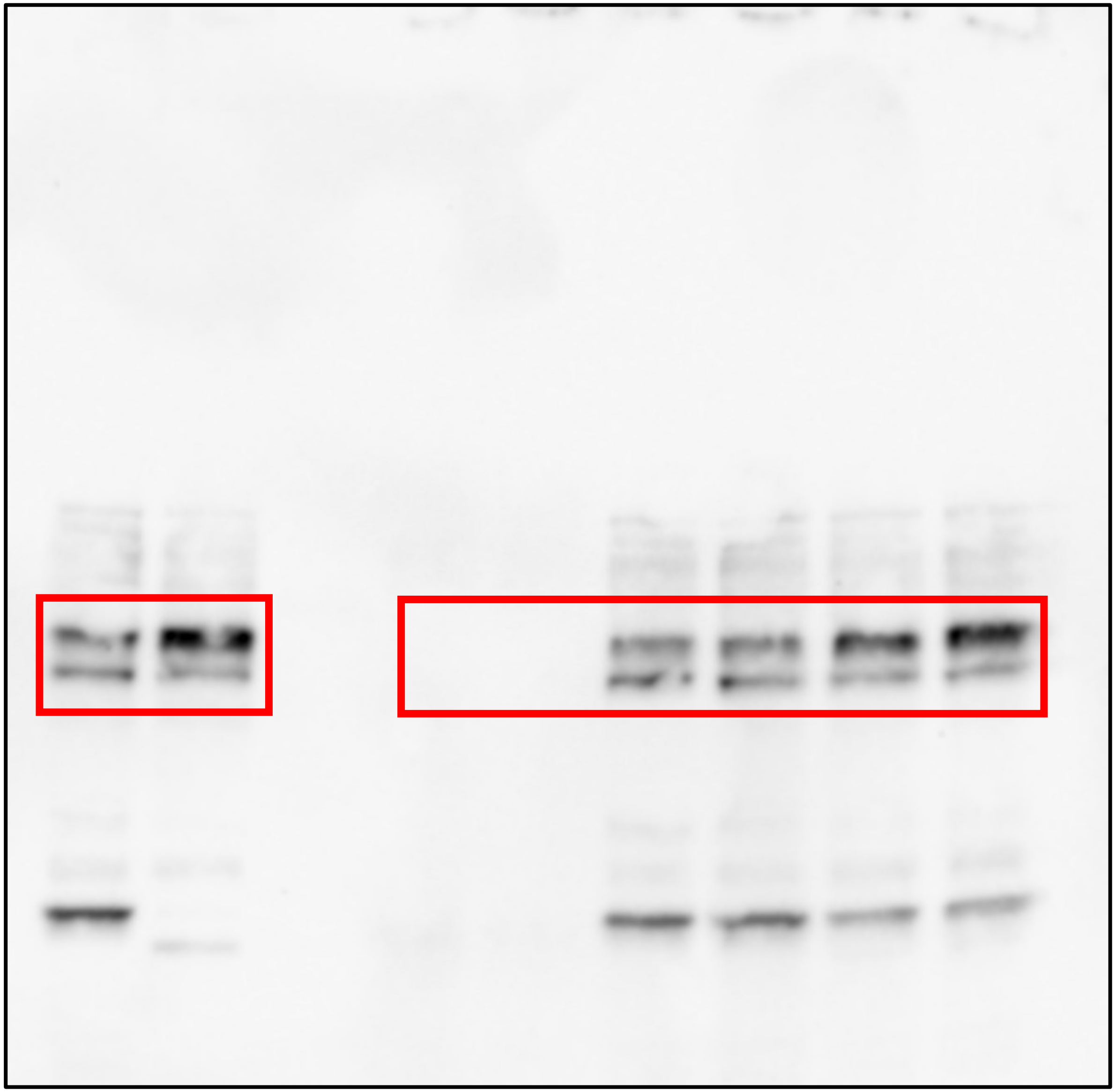

**Branscome et al. Supplementary Figure 11**

DICER (Supplementary Fig. 6)

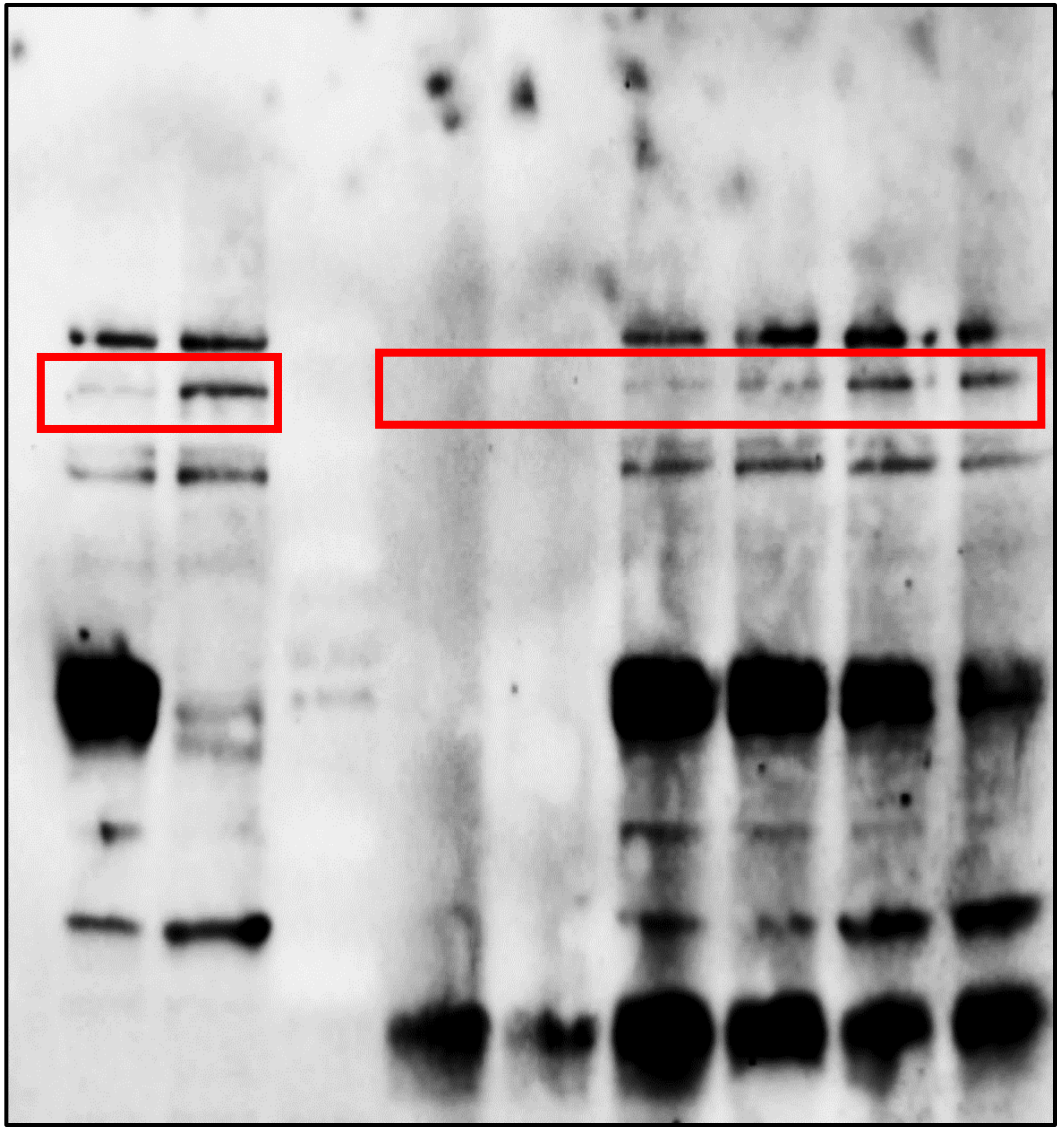

**Branscome et al. Supplementary Figure 11**

RIG-I (Supplementary Fig. 6)

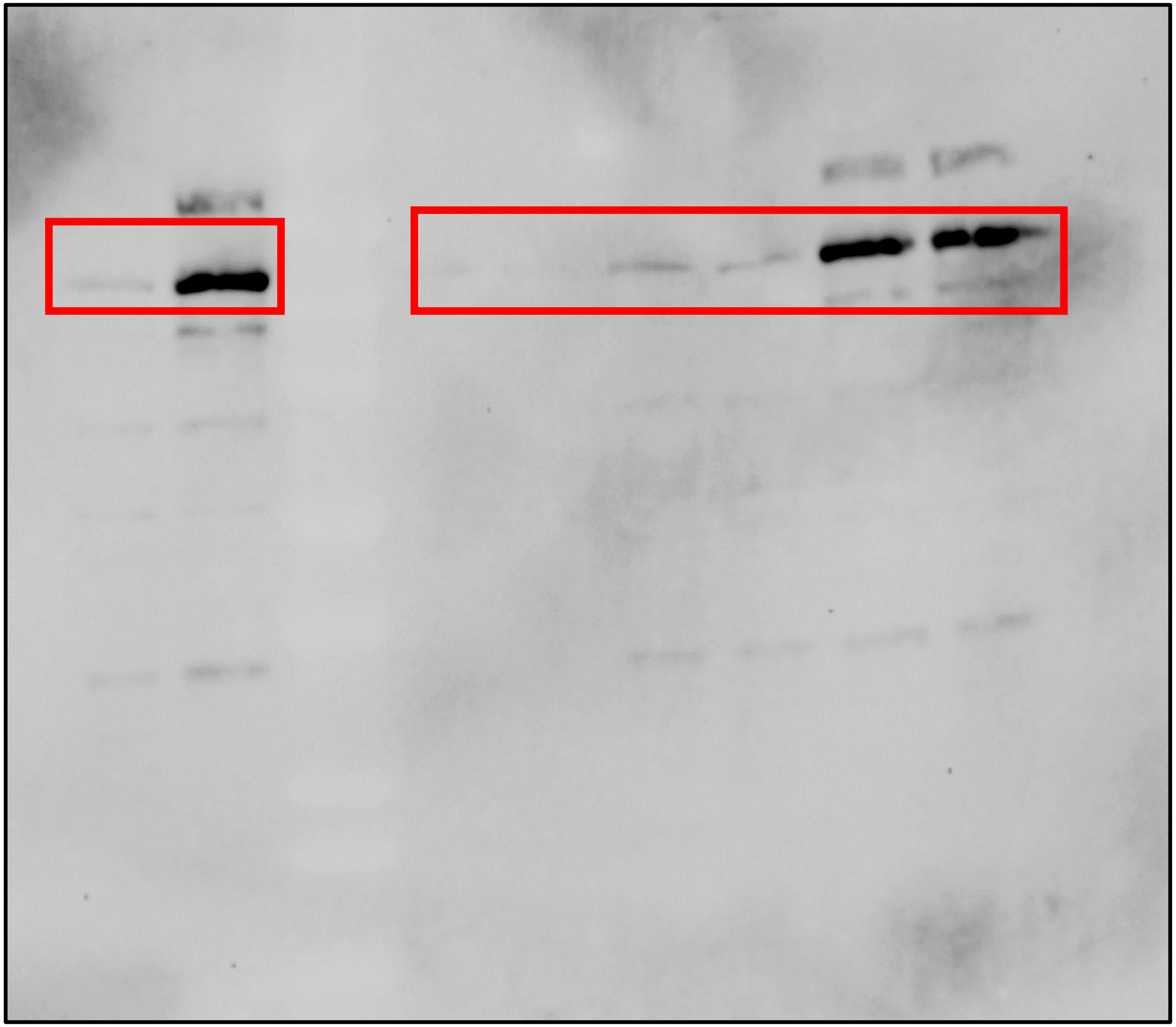

**Branscome et al. Supplementary Figure 11**

CD4 (Supplementary Fig. 8)

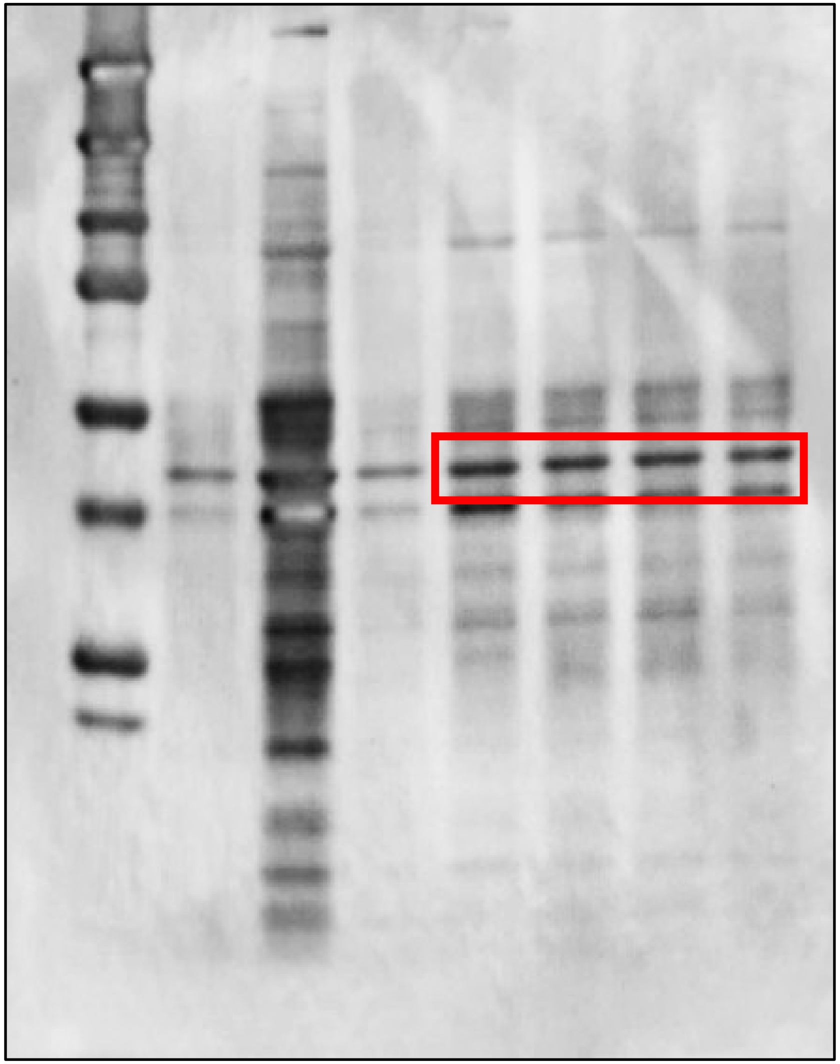

CCR5 (Supplementary Fig. 8)

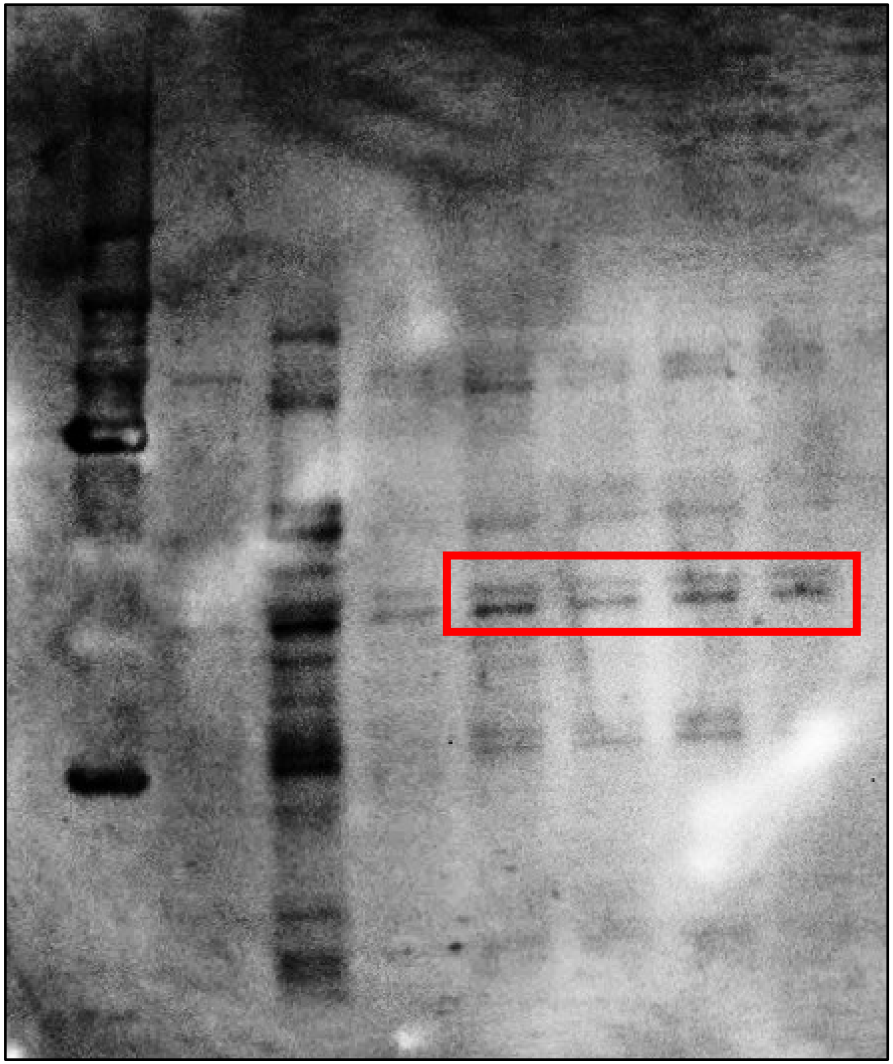

**Branscome et al. Supplementary Figure 11**

Actin (Supplementary Fig. 8)

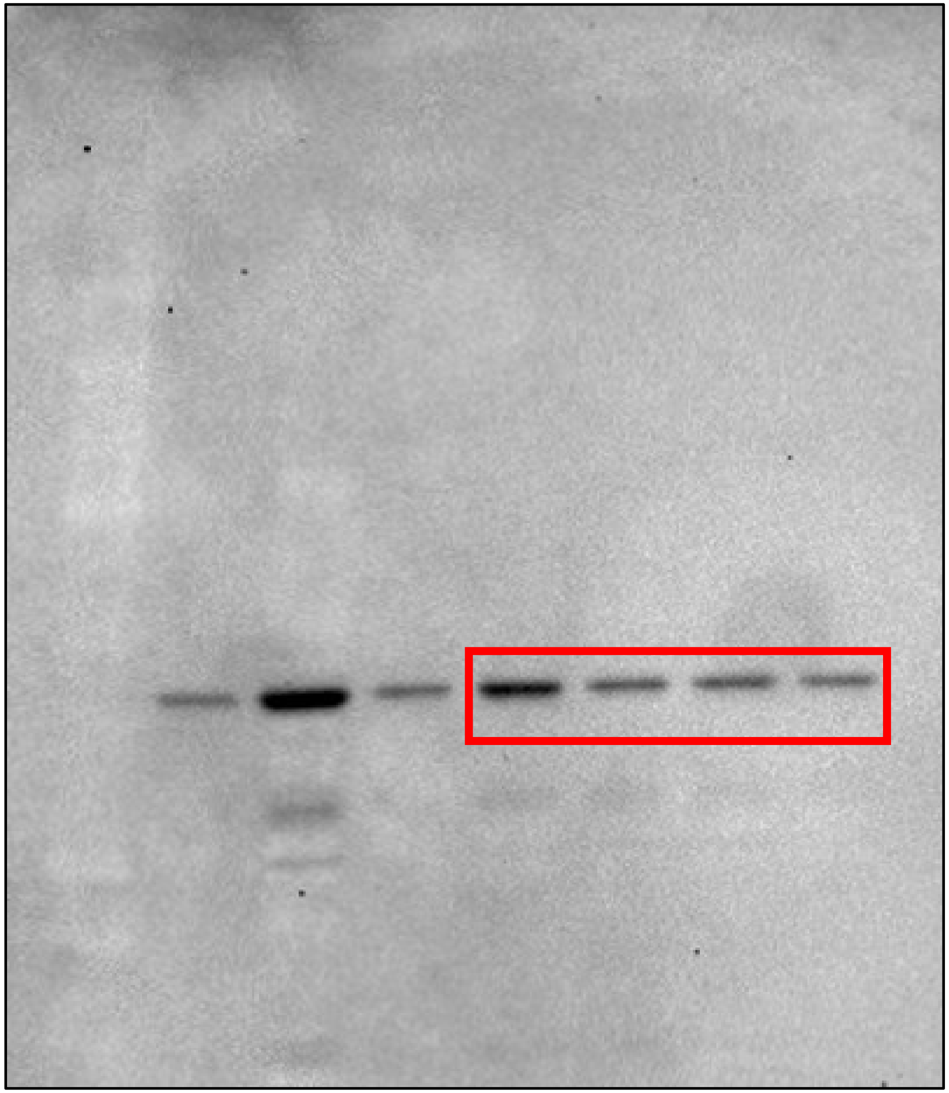

**Supplementary Figure 11**. Full-length blots corresponding to each western blot assay.

**Branscome et al. Supplementary Figure 11**
